# How Well Do Molecular Dynamics Force Fields Model Peptides? A Systematic Benchmark Across Diverse Folding Behaviors

**DOI:** 10.1101/2025.07.31.667969

**Authors:** Bhumika Singh, Yisel Martínez-Noa, Alberto Perez

## Abstract

Linear peptides play essential roles in biology and drug discovery, frequently mediating protein–protein interactions through short, flexible motifs. However, their structural plasticity—ranging from disordered to context-dependent folding—makes them challenging targets for molecular simulations. In this work, we benchmark the performance of eleven popular and emerging fixed-charge force fields across a curated set of twelve peptides spanning structured miniproteins, context-sensitive epitopes, and disordered sequences. Each peptide was simulated from both folded (200 ns) and extended (10 µs) states to assess stability, folding behavior, and force field biases. Our analysis reveals consistent trends: some force fields exhibit strong structural bias, others allow reversible fluctuations, and no single model performs optimally across all systems. The study highlights limitations in current force fields’ ability to balance disorder and secondary structure, particularly when modeling conformational selection. These results offer practical guidance for peptide modeling and establish a benchmark framework for future force field development and validation in peptide-relevant regimes.

## Introduction

Peptide epitopes—short regions within proteins that mediate molecular recognition—play central roles in biology and drug discovery. Many cellular interactions are driven by such short linear motifs^1,2^, which bind to specific partner proteins and often undergo disorder-to-order transitions in the process^3,4^. This structural plasticity is both biologically important and a significant modeling challenge.

Peptides are particularly attractive drug candidates because they can engage extended and shallow protein surfaces that are typically inaccessible to small molecules^5,6^. Advances such as peptide cyclization have improved their proteolytic stability and membrane permeability, further boosting their therapeutic potential^7^. Accurate modeling of peptide structure and binding is thus crucial for structure-based design of peptide therapeutics, peptidomimetics, and small molecules that mimic peptide–protein interactions.

Yet, modeling peptides remains difficult. Many peptides are intrinsically disordered in isolation and only fold upon binding to a target^8,9^. Others form defined secondary structures under specific solution conditions, but not others^10,11^. A singular protein receptor might recognize different peptide sequences via distinct binding modes (e.g., peptide binds as a helix or hairpin)^12^, while a given peptide sequence may bind different protein receptors by adopting different structures^13,14^. This environmental sensitivity places peptides near the edge of structural stability, where small changes in sequence, solvent, or receptor can trigger large conformational shifts. Unlike globular proteins, which are stabilized by dense networks of non-bonded interactions, peptides often lack such extensive stabilizing contacts^15,16^. As a result, even small inaccuracies in the force field can lead to significant errors in conformational preferences^17–19^.

From a computational standpoint, peptides are small enough that conformational sampling is not generally a limiting factor. However, their responsiveness to small perturbations makes them especially susceptible to force field biases^20–22^. For instance, modest secondary structure preferences encoded in a force field may not strongly affect the native fold of a protein but can dominate the ensemble of a flexible peptide^23^. Despite recent improvements, traditional force fields were largely developed and parameterized to describe well-folded proteins, often leading to over-compact ensembles for disordered systems.

Increasing computer time and access to better benchmark experimental data have shown several force field deficiencies, such as inaccurate helical propensities or incorrect compaction in disordered proteins^24,25^. This has led to an explosion in the number of groups participating in force field development as well as the number of available force fields.^26^ Yet, many of these developments have focused on large protein systems, and it remains unclear which force fields best describe peptides across different structural regimes^27,28^.

In this work, we systematically assess the performance of 11 modern force fields across a benchmark set of 12 peptides. These include peptides that are structured in solution, intrinsically disordered peptides, and peptides that remain disordered in water but fold in alternative solvent environments. We evaluate both structure stability (via simulations initiated from folded states) and folding behavior (via simulations initiated from extended conformations). While no force field performs optimally in all cases, our results reveal meaningful trends and biases, offering practical guidance for researchers modeling peptide-mediated interactions.

## Computational Methods

### Systems of study

A total of twelve peptides (see Table 1) were selected as model systems to evaluate and compare force fields.

**Table 1:**
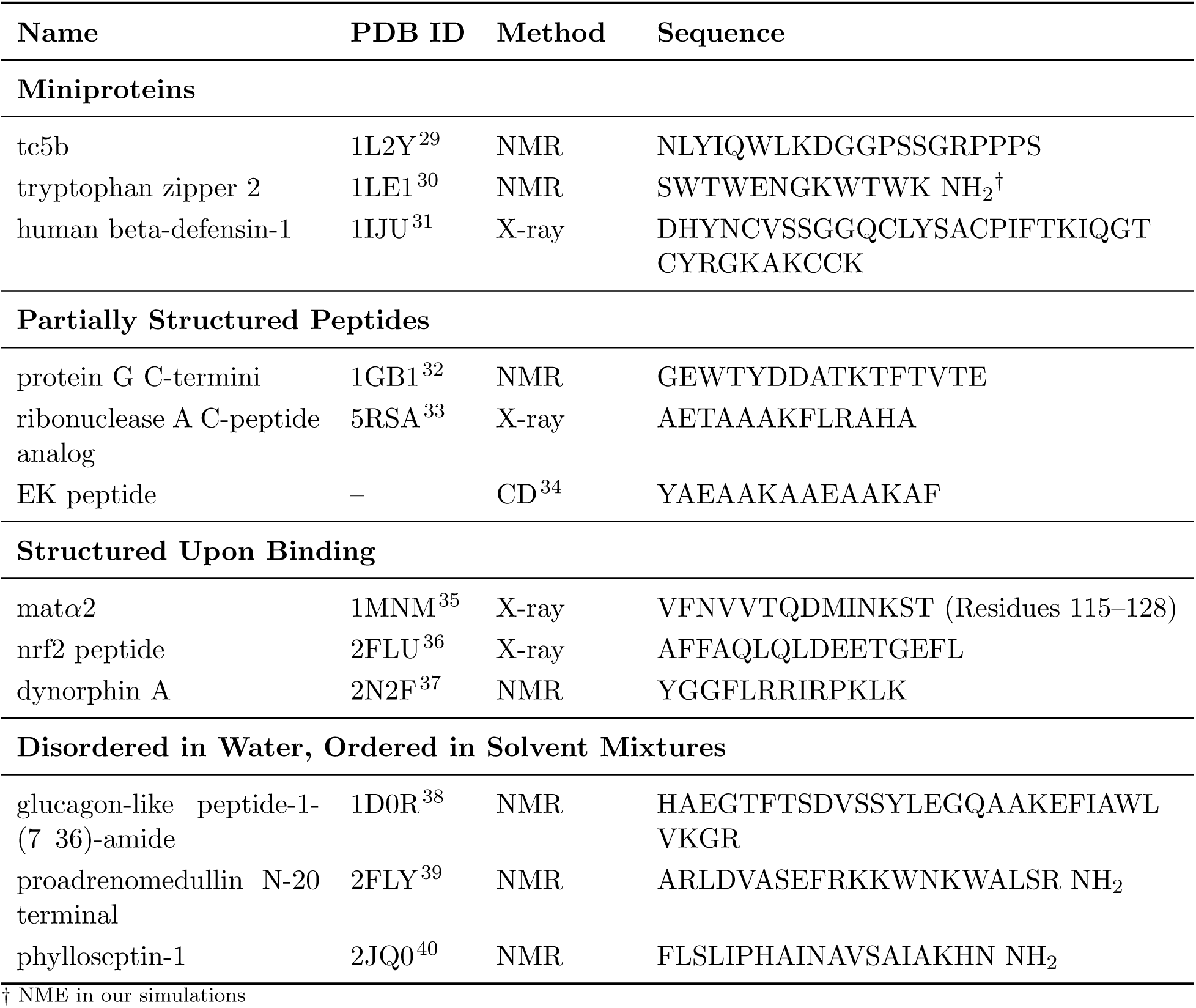
Benchmark systems used in this study.

### Force field choices

We took some traditional force fields ff19SB,^41^ ff99SB,^42^ and charmm36m^43^ commonly used in the field, as well as some IDP optimized force fields ff99IDPs,^44^ ff14IDPs,^45^ ff14IDPSFF,^46^ C36IDPSFF,^47^ OPLDIDPSFF,^48^ and some physics-enhanced modern force fields a99SBdisp,^17^ Des-Amber,^49^ Des-Amber-SF1.0.^49^ For each force field, we used the water model recommended for it (see table 2). While this is not an exhaustive list of force fields, it is representative of different trends in the field and readily available for running simulations.

**Table 2:**
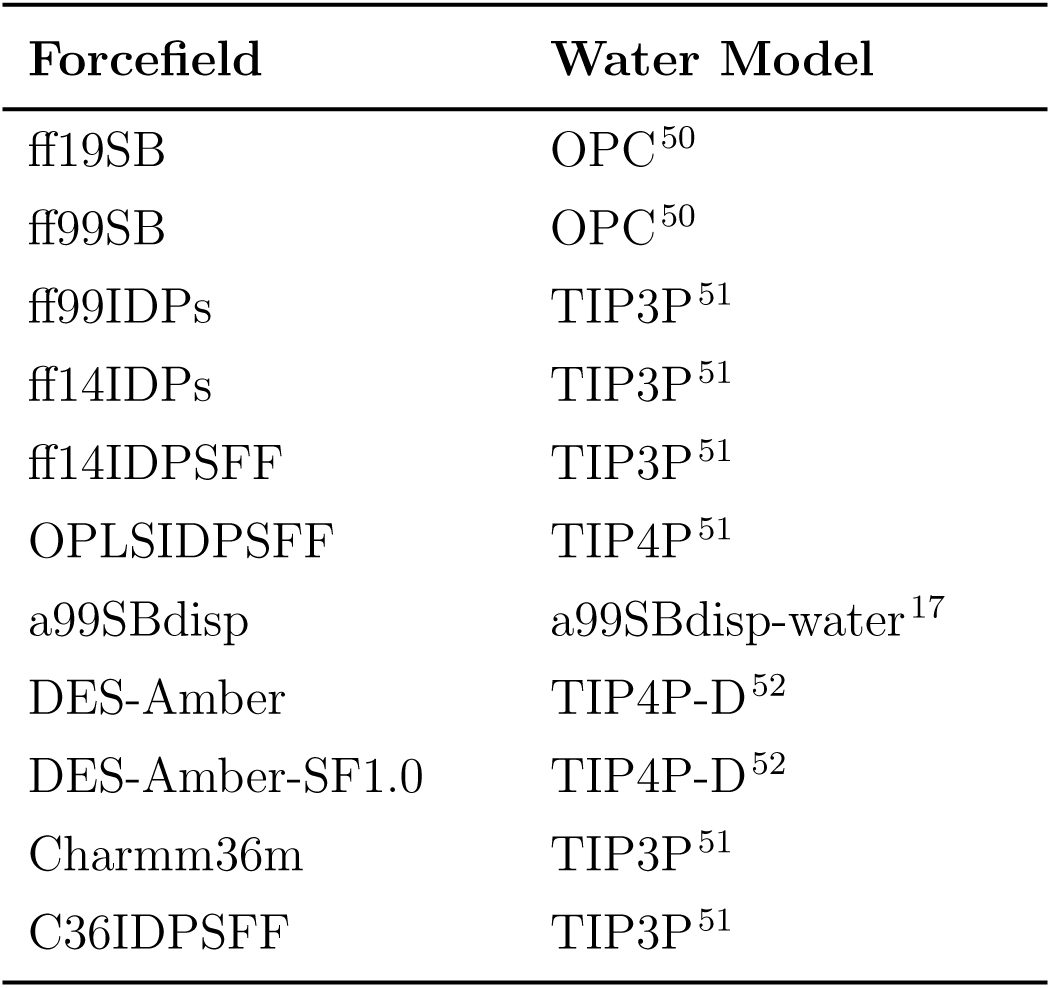
Summary of force fields and water models employed in this benchmark study.

| Forcefield | Water Model |
| --- | --- |
| ff19SB | OPC<sup>50</sup> |
| ff99SB | OPC<sup>50</sup> |
| ff99IDPs | TIP3P<sup>51</sup> |
| ff14IDPs | TIP3P<sup>51</sup> |
| ff14IDPSFF | TIP3P<sup>51</sup> |
| OPLSIDPSFF | TIP4P<sup>51</sup> |
| a99SBdisp | a99SBdisp-water<sup>17</sup> |
| DES-Amber | TIP4P-D<sup>52</sup> |
| DES-Amber-SF1.0 | TIP4P-D<sup>52</sup> |
| Charmm36m | TIP3P<sup>51</sup> |
| C36IDPSFF | TIP3P<sup>51</sup> |

### Simulation protocols

Molecular dynamics simulations were performed using twelve peptide force fields, including several variants incorporating CMAP corrections. The force fields and their recommended water models are summarized in Table 2. Simulations were conducted using AMBER^53,54^ and GROMACS^55^, depending on force-field availability. Simulations initiated from the unfolded state were generated from extended linear conformations constructed using the *xleap* module in AMBER, while folded-state simulations were initiated from experimentally determined structures retrieved from the RCSB Protein Data Bank (PDB)^56^. Simulations initiated from experimentally determined structures serve different interpretive roles depending on the peptide class. For miniproteins and partially structured peptides, the PDB structure represents a native-like state that is expected to remain stable in aqueous solution, and loss of structure indicates potential destabilization bias. In contrast, for peptides that are structured only upon binding or under specific solvent conditions, the PDB structure corresponds to a conformation stabilized in a different environmental context. For these systems, retention of the PDB-derived structure in water is not the expected outcome during simulations.

Each system was solvated in a truncated octahedral water box with at least 8 Å of padding between the solute and the box boundaries, and Na^+^ and Cl^−^ ions were added to neutralize the system and achieve a final salt concentration of 0.1 M. Hydrogen mass repartitioning^57^ was applied prior to production simulations to enable a 4 fs integration timestep, and trajectory frames were saved every 20 ps. For unfolded-state simulations, each system was simulated in triplicate, consisting of one 10 *µ*s trajectory and two additional 1 *µ*s trajectories. For folded-state simulations, five independent replicas of 200 ns each were performed.

#### Operational definitions

We use the term bias to denote a systematic tendency of a force field to preferentially stabilize specific secondary-structure motifs or compact conformational states that are inconsistent with the expected structural propensity class of a peptide (folded miniprotein, partially structured, structured upon binding, or solvent-induced). Evidence for bias is evaluated across multiple systems; recurring deviations from expected behavior are interpreted as indicative of force-field–specific tendencies rather than system-specific effects. Expected structural behavior is most clearly defined for well-structured peptides whose native conformations are stable in aqueous solution. For peptides whose experimentally determined structures are stabilized under different environmental conditions (e.g., binding partners or mixed solvents), the reference structure is not assumed to represent the dominant state in water. In such cases, sampling of a predominantly disordered ensemble is considered consistent with expected behavior, and persistent stabilization of ordered motifs is interpreted as potential overstabilization bias.

#### AMBER protocols

Molecular dynamics simulations using the ff19SB, ff99SB, ff99IDPs, ff14IDPs, ff14IDPSFF, and CHARMM36m force fields were performed with the AMBER MD suite. Systems with ff19SB and ff99SB were prepared directly using the standard AMBER parameter sets. Systems with ff14IDPs and ff14IDPSFF were first parameterized with ff14SB^58^ and then updated with the corresponding CMAP corrections, whereas ff99IDPs systems were initialized with ff99SB and modified using ff99IDPs-specific CMAP corrections. CHARMM36m systems were prepared using the CHARMM-GUI platform^59–61^.

Energy minimization was performed in six stages to relax the systems while preserving solute structure. In the first five stages, harmonic positional restraints were applied to solute heavy atoms (excluding hydrogens, solvent, and ions) with decreasing force constants of 50, 30, 20, 10, and 5 kcal mol^‒^1‒2 Å. This gradual reduction allowed solvent relaxation while preventing nonphysical solute distortions. In the final stage, all restraints were removed to allow full system relaxation prior to equilibration. Each stage consisted of 4000 steps, with the first 2000 using steepest descent followed by 2000 steps of conjugate gradient minimization. Systems were then heated from 0 K to 298.15 K over 5 ns using Langevin dynamics with a collision frequency of 1 ps*^‒^*^1^.

Equilibration was carried out in two stages under NPT conditions at 298.15 K and 1 atm, with SHAKE applied to all bonds involving hydrogens. In the first stage, positional restraints of 5 kcal mol^‒1^ Å^‒2^ were applied to solute heavy atoms (excluding hydrogens, solvent, and ions) for 10 ns using a Langevin thermostat (*γ* = 1 ps*^‒^*^1^). In the second stage, restraints were removed, and the system was equilibrated for an additional 10 ns with a higher collision frequency (*γ* = 2 ps*^‒^*^1^). Both stages used a 10 Å cutoff for nonbonded interactions.

Production simulations were conducted with a 4 fs timestep using hydrogen mass repartitioning. Systems starting from extended structures were simulated for 10 *µ*s, with two additional replicas of 1 *µ*s each, while folded systems were simulated in five independent replicas of 200 ns each.

#### GROMACS protocols

Simulations using the OPLSIDPSFF^48^, a99SBdisp^17^, DES-Amber^49^, DES-Amber-SF1.0^49^, and C36IDPSFF^47^ force fields were performed using GROMACS version 2019.2^55^. To ensure consistency with the AMBER-based simulations, similar protocols and conditions were applied across all systems to generate their topologies and coordinates.

Energy minimization was performed in two stages: an initial steepest descent minimization of up to 50,000 steps, followed by 2,000 steps of conjugate gradient minimization^62^. The systems were then gradually heated to 300 K over 10 ns in the canonical (NVT) ensemble, followed by a 20 ns equilibration in the isothermal-isobaric (NPT) ensemble. Covalent bonds involving hydrogen atoms were constrained using the LINCS algorithm^63^. Lennard-Jones and short-range electrostatic interactions were truncated at 10 Å. Long-range electrostatics were treated using the Particle Mesh Ewald (PME) method^64^. Temperature was controlled using the velocity-rescale thermostat set at 300 K, and pressure was maintained at 1 atm using the Parrinello-Rahman barostat^65,66^.

Production simulations were performed with a 4 fs timestep using hydrogen mass repartitioning. As in the AMBER protocol, systems starting from extended structures were simulated for 10 *µ*s, with two additional replicas of 1 *µ*s each, while folded systems were simulated in five independent replicas of 200 ns each.

Trajectory analysis and visualization for all the trajectories generated using AMBER and GROMACS were performed using CPPTRAJ, MDAnalysis^67,68^ and VMD^69^.

### Analysis

Prior to analysis, water molecules and ions were removed from all trajectories, and the coordinates were autoimaged to recenter the system at the origin of coordinates. The resulting processed trajectories were then used for all subsequent analyses.

#### Ramachandran Plots

To evaluate the stereochemical quality and conformational preferences of amino acid residues within the peptide sequences, `Ramachandran`^70^ plots were generated. These plots provide a visual representation of the backbone dihedral angles (*ϕ* and *ψ*), offering insights into the propensity of residues to adopt specific secondary structural elements such as *α*-helices, *β*-sheets, or random coils. The analysis was performed using the MDAnalysis toolkit, employing the Ramachandran class to compute *ϕ* and *ψ* angles across all residues over time. Stripped and aligned trajectories, along with the corresponding topology files, were used as input to ensure accurate dihedral angle calculations.

#### Clustering

To identify representative conformations and their populations sampled during the simulations, structural clustering was performed on each trajectory for every force field and system. Hierarchical agglomerative clustering with average linkage was applied using an RMSD-based metric with an *ɛ* cutoff of 3.0 Å, as implemented in `cpptraj`, part of the `AmberTools` suite. Clustering was based on the backbone heavy atoms (C, N, O, C*α*, and C*β*) of all peptide residues. Cluster centroids were selected as representative structures.

To enable robust comparison across force fields, clustering analyses were performed separately for three simulation categories: (i) meta-trajectories constructed by combining all short replicas from folded-start simulations, (ii) meta-trajectories constructed by combining the three unfolded-start trajectories of 1 *µ*s each, and (iii) the single 10 *µ*s unfolded-start trajectory. Rather than interpreting individual cluster identities, we focus on relative measures of conformational heterogeneity across force fields. In this context, Shannon entropy (see below), computed from cluster populations, serves as a compact proxy for distinguishing force fields that exhibit systematically different sampling behavior from the consensus. Entropy comparisons are restricted to a given peptide system and a single simulation category, and are not performed across categories, ensuring that differences in trajectory length or sampling density do not confound interpretation.

#### Secondary Structure preferences

Secondary structure content of the simulated peptides was analyzed using the `secstruct` command in `cpptraj`. Secondary structure assignment was carried out using the DSSP algorithm^71^ as implemented in `cpptraj`, which classifies structural elements—including *α*-helices, *β*-strands, turns, and coils—based on backbone hydrogen bonding patterns and geometric features.

#### Conformational Diversity and Structural Deviation Analysis

To quantify the conformational diversity sampled by each force field, Shannon entropy^72^ was calculated based on the fractional populations of clusters obtained from the hierarchical clustering analysis (as described in the *Clustering* section). This metric captures the degree of structural heterogeneity within a simulation, where higher entropy values indicate greater conformational diversity. Entropy, *S*, was calculated using the following equation:

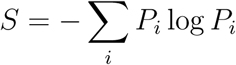

where *P_i_* is the population fraction of the *i*-th cluster. Calculations were performed with the pandas^73^ and `NumPy`^74^ Python libraries.

Because entropy values depend on clustering resolution, they are interpreted as relative, within-system indicators of heterogeneity rather than absolute thermodynamic quantities. In this work, entropy is used as a proxy to rapidly assess consensus versus divergence in sampling behavior across force fields for a given peptide system.

To quantify structural deviations from the reference state, root-mean-square deviation (RMSD) was computed for the most populated cluster for each force field in each system. The reference structure corresponded to the experimentally determined folded state of the peptide. RMSD was calculated using the backbone heavy atoms (C, N, O, C*α*, and C*β*) of all residues, employing `cpptraj` from the AmberTools suite.

When interpreted together with clustering entropy, RMSD enables discrimination between (i) low-entropy ensembles dominated by reference-like conformations, (ii) low-entropy ensembles stabilized in alternative, non-reference conformations, and (iii) high-entropy ensembles characterized by broad conformational sampling. This analysis strategy minimizes sensitivity to clustering artifacts while preserving the ability to detect systematic force-field biases across diverse peptide classes.

## Results

### Systems of study

The structural characterization of peptides presents a particularly stringent test for molecular force fields. While most peptide sequences are intrinsically disordered in solution, certain peptides have been experimentally observed to adopt defined secondary structures under specific conditions—such as binding to a partner protein, folding in a membrane-mimetic solvent, or being excised from a larger folded domain. These context-dependent peptides occupy an intermediate regime between fully folded miniproteins and disordered chains, making them sensitive probes for assessing force field accuracy and bias. We selected twelve experimentally characterized peptides that span four categories: (1) miniproteins with well-defined structures in solution, (2) partially structured peptides, (3) peptides that fold upon binding to a protein target, and (4) disordered peptides that adopt defined structure only in membrane-mimetic or mixed solvents. Categories (3) and (4) are intentionally near-threshold systems: they are disordered in water yet become structured under modest perturbations (binding or solvent composition). These systems are particularly diagnostic for bias because a force field that artificially stabilizes ordered motifs in water will appear as a deviation from expected IDP behavior. Together, this set (see Fig. 1) enables us to evaluate force field performance across a biologically meaningful spectrum of structural propensities.

**Figure 1:**
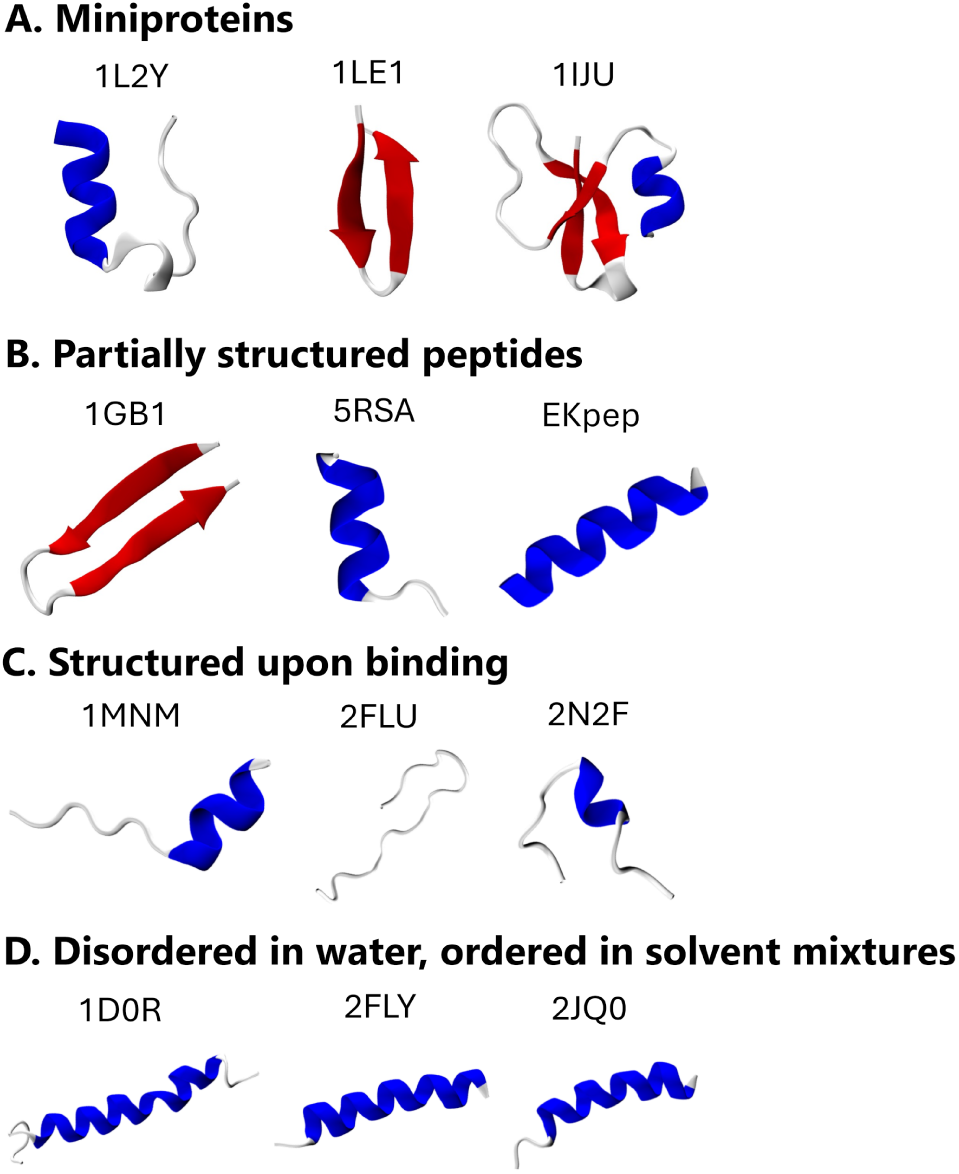
Experimentally folded structures of benchmark peptide systems used in this study, shown with their corresponding PDB IDs. The systems are grouped into four categories, each containing three representative systems: (A) miniproteins, (B) partially structured peptides in aqueous solvent, (C) peptides that adopt structure upon binding, and (D) peptides that are disordered in water but become ordered in solvent mixtures.

#### Miniproteins

These peptides adopt well-defined, compact tertiary folds under aqueous conditions and are considered stable, folded entities in solution. Trp-cage (1L2Y) is often classified as a miniprotein, as it exhibits over 95% folded population at physiological pH, forming a compact helical bundle stabilized by a hydrophobic core nucleated around a tryptophan residue.^29^ This 20 residue sequence was optimized starting from a 39 residue protein. Trp-zip (1LE1) is a 13-residue *de novo* designed sequence that folds as a *β*-hairpin. Four tryptophan residues create a hydrophobic side in the hairpin, while polar and charged residues are present in the other side. Trp-zip is highly stable with high hairpin populations (*>*95%) in water, as confirmed by NMR.^30^ Finally, *β*-defensin-1 (1IJU) is a natural antimicrobial peptide that forms stable monomers and dimers in crystallographic studies. NMR of a homologous defensin confirms that monomeric structure is preserved in dilute solution. ^31,75^ This 36 residue sequence is the longest in our dataset.

#### Partially structured peptides

These peptides exhibit some degree of structure in solution, often sampling a mixture of folded and unfolded conformations. Two of them are fragments derived from larger proteins, whereas the third one is a designed peptide. The C-termini from Protein G (*β*-hairpin, 1GB1) is a 16-residue segment from the immunoglobulin binding domain of streptococcal protein G. This hairpin is expected to be formed early in protein G folding^76^ and experimental studies suggests that this peptide should adopt ⇠20% hairpin population in solution.^30^C-peptide (helical, 5RSA) is a 13-residue analog from the RNase A N-terminal with moderate *α*-helical content in water. The original C-termini had a small helical population in solution and analogs produced with sequence optimization increased the helical stability with populations ranging from 40–60% depending on temperature and variant.^77,78^ The EK peptide is a designed helical peptide with favorable electrostatics (E,K at *i, i* +4 positions). Circular dichroism (CD) studies estimate 40% helical population in solution.^34^ The AlphaFold^79^ prediction is used as a reference.

#### Structured upon binding

These peptides are expected to be disordered or dynamic in isolation but adopt specific folded conformations upon binding to a protein receptor, making them ideal sensitive probes to force-field biases that might over-stabilize the free form (2FLU, 2N2F). The MAT*α*2 epitope (1MNM) is a context-dependent peptide from MAT*α*2 (residues 115-128) that has been observed binding as both a helix or a hairpin in the same crystal structure.^35^ This unique chameleon-like behavior is a practical test case for our force fields.

#### Disordered in water, ordered in solvent mixtures

These systems adopt helical structure in mixed solvent environments such as Trifluoroethanol (TFE)/water or membrane mimetics but remain disordered in pure water. They serve as sensitive probes for helical bias. The glucagon-like peptide-1 (GLP-1, 1D0R) is a peptide hormone that is unstructured in water but adopts a helix in 35% TFE. NMR experiments at different TFE concentrations shows that the C-terminal region has a higher helix propensity.^38^ The PAMP peptide (2FLY) also exhibits *α*-helicity in both membrane mimetic and TFE environments according to NMR studies.^39^ Finally, Phylloseptin-1 (2JQ0) is an antimicrobial peptide whose helical population increases with TFE concentration—from 53% at 30% to 70% at 60%.^40^ To evaluate force field performance for these systems, we conducted two sets of simulations for each system: 1) Short simulations (five independent replicas of 200 ns) initiated from the experimentally determined or AlphaFold-predicted (EK peptide) structure, to assess structural stability and strong biases. 2) Long simulations (10 *µ*s, with two additional replicas of 1*µ*s) initiated from extended conformations, to assess folding tendencies, sampling breadth, and subtle biases.

Each system’s simulation outputs— Ramachandran analysis, secondary structure propensities, cluster entropy, RMSD trajectories, representative conformers—are summarized in the main text and detailed in Supplementary Figures S1–S86. These results guide our analysis of force field accuracy and conformational bias.

### Modeling peptide stability

Short, 200 ns simulations starting from a folded conformation provide a practical means to detect strong force field biases. While unfolding proteins and nucleic acids in explicit solvent simulations in denaturing solvent mixtures requires long timescales, ^80,81^ peptides—due to their smaller size and limited stabilizing contacts—can unfold spontaneously within this timescale, especially when the native structure is marginally stable or environment-dependent.^80,81^ Thus, we use this protocol to detect whether force fields maintain starting structural features or quickly deviate toward alternative states.

#### Miniproteins

For the three miniproteins (1L2Y, 1LE1, 1IJU), which are experimentally stable in solution, most force fields correctly preserve the native fold over 200 ns simulations. Trp-cage (1L2Y) remains largely compact and helical across force fields, with only IDP-optimized variants showing occasional *α*-helix destabilization. The *β*-hairpin Trpzip (1LE1) is robustly maintained in nearly all force fields, with noticeable disruption only in ff99IDPs, indicating a tendency toward excessive disorder.

*β*-defensin (1IJU), the longest and most complex miniprotein in the set, is also generally preserved; however, IDP-biased force fields and, to a lesser extent, CHARMM36m show partial strand or N-terminal *α*-helix destabilization, consistent with previous observations for *β*-defensin-2.^31^ In the most pronounced case (ff99IDPs), loss of global *β*-sheet organization is observed despite retention of some local strand pairing.

Overall, structure-preserving force fields (e.g., ff19SB, a99SBdisp, des-amber variants) maintain native topology consistently, whereas IDP-oriented parameterizations display a systematic bias toward partial unfolding in systems that are experimentally stable.

#### Partially structured peptides

These peptides (1GB1, 5RSA, EK) are expected to adopt partially folded conformations in aqueous solution. On the 200 ns timescale, we anticipate either full or partial stabilization of native secondary structure. Extensive unfolding followed by refolding is not expected within this window.

For the Protein G C-terminal hairpin (1GB1), most force fields maintain the native *β*-hairpin over 200 ns, consistent with experimental evidence that folded conformations are present in the ensemble (SI Fig. 22–28). However, ff99IDPs consistently destabilizes the hairpin tail and samples transient helical turns in that region, indicating a mild secondary-structure redistribution. In several other force fields, partial fraying of the strand termini is observed, though without systematic conversion to alternative compact folds.

For C-peptide (5RSA), helicity is less stable. More than half of the force fields show early loss of *α*-structure (SI Fig. 29–35). Des-amber-SF1.0 exhibits the most consistent helical retention across replicas, while a99SBdisp and des-amber preserve helicity in a subset of trajectories. In contrast, ff99IDPs and portions of ff19SB trajectories display unfolding with only partial recovery. In rare cases, alternative *β*-like conformations are sampled late in the simulation, though without persistent stabilization.

EK, a designed helix, reveals clearer force-field distinctions (SI Fig. 36–42). Several models—including ff14IDPs, ff14IDPSFF, OPLSIDPSFF, and CHARMM36m—lose helicity rapidly and do not recover it within 200 ns. In contrast, ff19SB and ff99IDPs exhibit unfolding followed by partial refolding, suggesting a shallow but accessible helical basin. Notably, OPLSIDPSFF and C36IDPSFF intermittently sample *β*-hairpin-like conformations, indicating a strand-stabilizing tendency even in a sequence designed to remain helical.

Overall, while transient fraying and partial loss of structure are not interpreted as bias on this timescale, the repeated emergence of alternative *β*-like motifs or systematic helicity loss in certain force fields suggests imbalance in secondary-structure propensities.

#### Structured upon binding

These peptides (1MNM, 2FLU, 2N2F) fold only upon receptor binding and are therefore expected to exhibit low intrinsic stability as isolated monomers. Accordingly, both retention and loss of the initial structure in the five 200 ns replicates are reasonable outcomes. However, the spontaneous formation and stabilization of alternative folded conformations in isolation would indicate a force-field bias.

1MNM (SI Fig. 43–49) was initiated in a helical conformation, although it adopts a hairpin when bound. Four force fields rapidly lose the helix and predominantly sample disordered conformations (Fig. S44). Five others retain helical structure for longer times but lose it in at least one replicate. ff14IDPSFF and OPLSIDPSFF lose the helix rapidly but, in some replicates, locate and stabilize a *β*-hairpin. The repeated appearance of this alternative compact state suggests a strand-stabilizing tendency in these force fields.

NRF2 (2FLU) (SI Fig. 50–56) begins as a short C-terminal hairpin. In the 200 ns replicates, this hairpin flickers on and off but is generally maintained. ff19SB, ff99SB, ff99IDPs, and ff14IDPS show the strongest helical tendencies at short timescales. OPLSIDPSFF, which consistently favors hairpin formation, stabilizes an alternative *β*-hairpin distinct from the starting structure in multiple replicates (Fig. S51). C36IDPSFF occasionally samples the original hairpin but only sparsely. These results indicate competing secondary-structure preferences across force fields even on short timescales.

2N2F (SI Fig. 57–63) contains a central helix that is generally unstable in isolation. In the 200 ns replicates, the helix is rapidly lost in most force fields, with only isolated long-lived events in ff19SB and des-amber (Fig. S58). Across all force fields, structural heterogeneity dominates, consistent with the expectation that this peptide is structured primarily upon binding and should not stabilize a persistent folded basin in aqueous solution.

#### Solvent-sensitive helices

The last group comprises peptides (1D0R, 2FLY, 2JQ0) (SI Fig. 64–84) that form helices in membrane-mimetic or TFE-containing solvents but are disordered in water. Experimental data show that changing solvent composition modulates helix length, suggesting that certain sequence regions nucleate helix formation while others promote helix extension under favorable conditions. Thus, in aqueous simulations, rapid loss of helicity is expected, whereas persistent secondary structure or stabilization of alternative folds may indicate force field bias.

Across the three systems, ff19SB, a99SBdisp, des-amber, des-amber-SF1.0, and CHARMM36m generally maintain substantial helical structure over the five 200-ns replicas. For 1D0R and 2FLY, these force fields preserve *α*-helical conformations in all replicas, whereas less helical content is preserved for 2JQ0.

IDP-tailored force fields–including ff99IDPs, ff14IDPs, ff14IDPSFF, and OPLSIDPSFF– frequently lose helicity rapidly and exhibit a mild preference for *β*-hairpin conformations across all three peptides. Similarly, ff99SB and, to a lesser extent, C36IDPSFF disrupt helicity early across all systems.

Overall, these results indicate that force fields with strong secondary structure tendencies can affect the misfolding landscape even for peptides expected to remain disordered in water.

### Summary

Taken together, these 200 ns simulations reveal consistent trends across peptide classes (Fig. 2 and SI Table 1). Force fields such as ff19SB and a99SBdisp strike a balance between stability and flexibility, maintaining expected structure in miniproteins and partially structured peptides while allowing partial unfolding in binding-induced and solvent-sensitive systems. In contrast, ff99IDPs (and ff99SB) frequently destabilize folded helices and occasionally introduce *β*-structures in non-*β* contexts. OPLSIDPSFF, and to a lesser extent CHARMM36m and C36IDPSFF, show occasional strand bias, highlighting how tuning for disordered ensembles can influence the behavior of context-sensitive peptides.

**Figure 2:**
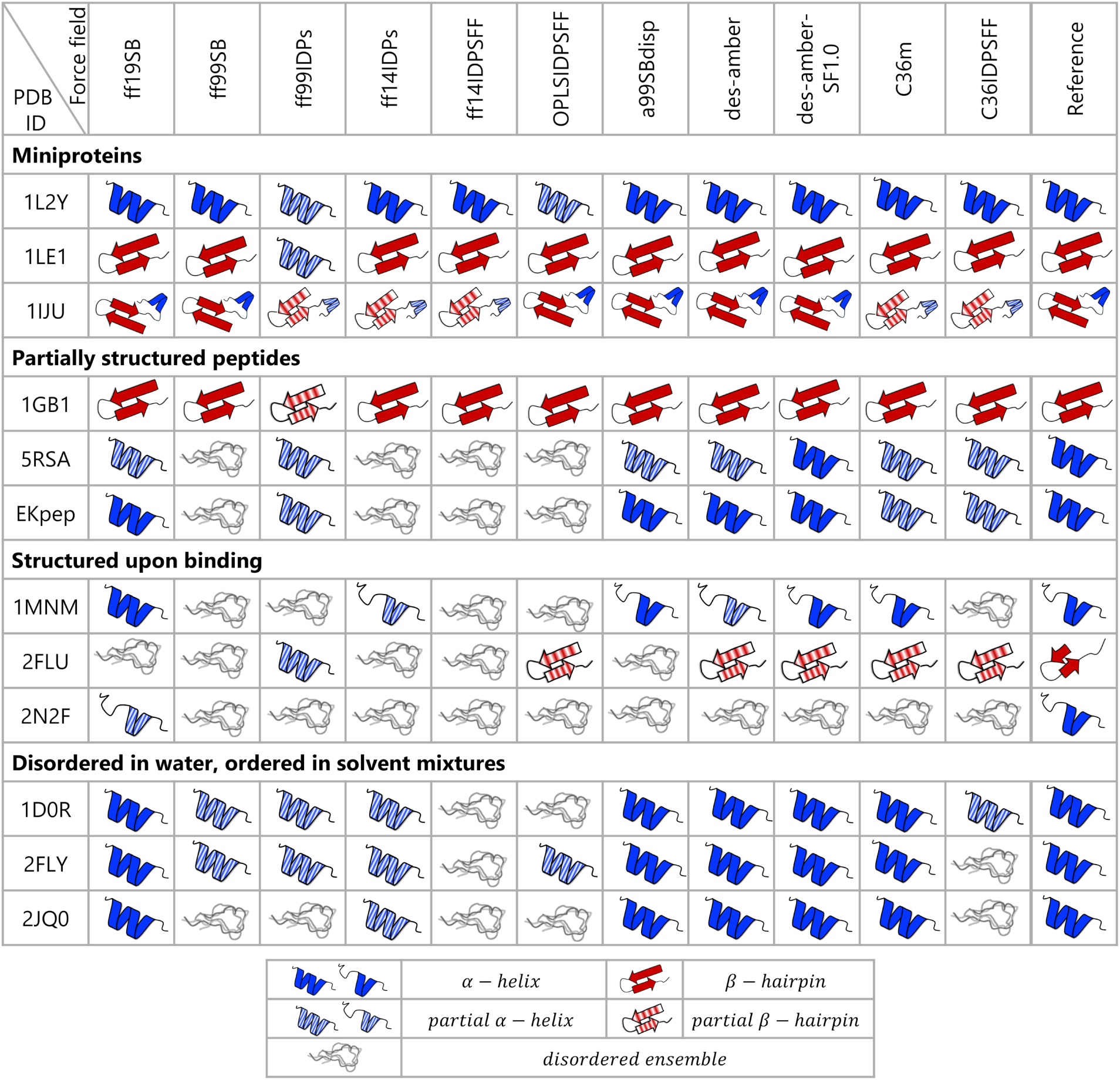
Overall secondary-structure trends across force fields in 200 ns simulations initiated from experimentally folded structures. For each peptide (rows), the dominant behavior across five replicas is summarized for each force field (columns). Solid helices or hairpins indicate stable native secondary structure; striped elements denote partial retention or stabilization of alternative structure; wavy lines indicate predominantly disordered sampling. The “Reference” column shows the experimental PDB structure used as the starting model (free, complexed, or determined in mixed solvent).

### Modeling peptide folding

While short simulations reveal whether force fields preserve native structures, they cannot assess whether force fields can discover such conformations from unfolded states. We therefore turn to long-timescale simulations initiated from extended conformations to examine folding behavior, landscape preferences, and potential biases. This approach helps detect dominant basins in the energy landscape and potential biases—e.g., force fields that prefer *β*-hairpins across many sequences or that stabilize incorrect folds once visited.

Some peptides that appeared disordered in the 200 ns simulations now display unexpected structure preferences, with certain force fields systematically promoting strand pairing or helix formation. Conversely, some force fields fail to reach native-like states for peptides known to be structured. Notably, OPLSIDPSFF frequently stabilizes *β*-hairpins early and maintains them, regardless of the peptide’s native fold. Force fields such as ff19SB and ff99IDPs often revisit native-like helices intermittently, suggesting balanced sampling.

#### Miniproteins

**For Trp-cage (1L2Y)**, composed of two *α*-helical segments packed around a hydrophobic core, none of the 1 µs triplicate simulations recover the native fold in any force field. Instead, several force fields—including ff99SB, ff14IDPs, ff14IDPSFF, OPLSIDPSFF, and des-amber-SF1.0—display a clear preference for *β*-structure. ff99IDPs samples partial *α*-helices but does not converge to the native topology, while the remaining force fields predominantly sample disordered ensembles without a well-defined folded state.

Only upon extending simulations to 10 µs does native-like folding emerge, and then only for ff19SB and a99SBdisp (SI Fig. 1–7). Of these, ff19SB stabilizes the folded structure for longer continuous periods. In contrast, OPLSIDPSFF and several IDP-tuned variants preferentially stabilize *β*-hairpin-like conformations once formed, indicating a persistent strand bias. Other force fields intermittently sample helical segments but fail to converge to the correct compact fold.

**For Trpzip (1LE1)**, a highly stable *β*-hairpin, the 1 µs triplicates already reveal clear force-field tendencies (SI Fig. 8–14). Strand-biased force fields—most notably OPLSIDPSFF, C36IDPSFF, and ff14IDPSFF—frequently sample and stabilize *β*-hairpin conformations within this timescale. Additional force fields (ff99SB, ff14IDPs, des-amber) recover the native hairpin in at least one replica, typically at later times. In contrast, ff19SB exhibits intermittent *β*-sheet formation with competing *α*-helical tendencies, while ff99IDPs and a99SBdisp favor *α*-helical conformations rather than converging to the native *β*-hairpin.

Extending the simulations to 10 µs sharpens these distinctions. Six force fields recover the correct *β*-hairpin topology within 2.5 µs, with OPLSIDPSFF, C36IDPSFF, and ff14IDPSFF showing the most consistent stabilization across replicas. Helix-prone models continue to sample alternative *α*-helical states, indicating a secondary-structure bias that competes with the native strand pairing. Overall, Trpzip exposes a clear separation between strand-stabilizing force fields that readily locate the correct basin and helix-favoring models that divert sampling toward non-native conformations.

**For** *β***-defensin-1 (1IJU)**, the longest peptide in this set, the 1 µs triplicates do not show convergence toward the native three-stranded *β*-sheet topology in any force field (SI Fig. 15–21). Instead, sampling is dominated by non-native secondary structures. OPLSIDPSFF and ff14IDPSFF frequently stabilize alternative *β*-hairpin or extended *β*-sheet motifs, while several other force fields sample persistent non-native elements.

Extending the simulations to 10 µs does not alter this qualitative picture. No force field folds the system to the native *β*-sheet structure. Rather than progressively forming isolated native-like elements, the trajectories predominantly stabilize incorrect folds. For example, ff19SB and des-amber repeatedly form a helix spanning residues 18–24, where the experimental structure contains a loop, indicating a misplaced helical basin. Across all force fields, the dominant clusters correspond to alternative compact structures rather than partial assembly of the native topology. Together, these results suggest that for this larger miniprotein, the native basin is either too shallow or kinetically inaccessible within the simulated timescale, and that several force fields instead favor competing non-native secondary-structure minima.

#### Partially structured peptides

For the Protein G C-terminal hairpin (1GB1), the 1 µs triplicates again reveal clear separation among force fields (SI Fig. 22–28). OPLSIDPSFF rapidly folds the *β*-hairpin within the first 100 ns and maintains it throughout the trajectory, reflecting a strong strand-stabilizing tendency. C36IDPSFF also captures hairpin conformations in select replicas. In contrast, ff14IDPs, ff99IDPs, and a99SBdisp frequently favor *α*-helical conformations, though without consistent convergence; cluster populations remain distributed, indicating competing basins rather than stable native assembly. The remaining force fields sample structurally diverse ensembles with limited persistent secondary structure.

In the 10 µs simulations, hairpin-like states become the most populated clusters in desamber, C36IDPSFF, and ff14IDPs, although these clusters represent a modest fraction of the total ensemble (*<* 15%). Conversely, ff19SB, ff99SB, and des-amber-SF1.0 occasionally sample helical conformations, but at low populations (*<*10%), indicating shallow competing basins rather than stable alternatives. Overall, 1GB1 highlights that even partially structured peptides can expose subtle secondary-structure biases, with strand-prone force fields rapidly stabilizing compact hairpins and helix-prone models diverting sampling toward alternative motifs. Since most force fields shift between some non-native secondary structure elements without stabilizing them for long periods of time, and given the stability of most force fields in the 5 replicates starting from native, it is possible that longer simulation times are required to sample native-like structures for this system.

EK and C-peptide (5RSA) both exhibit moderate *α*-helical populations experimentally. In the 1 µs triplicates, only ff19SB and ff99IDPs consistently sample and stabilize helical conformations for both systems (SI Fig. 29–42). ff99SB also stabilizes *α*-structure for EK, though less consistently across replicas. In contrast, OPLSIDPSFF frequently samples *β*-hairpin-like states, indicating a competing strand basin. The remaining force fields predominantly explore disordered ensembles or transient secondary structure without convergence. Extending simulations to 10 µs reinforces these trends rather than altering them. ff19SB and ff99IDPs intermittently recover and stabilize helical conformations, suggesting that the native-like basin is accessible but shallow. Other force fields either remain structurally disordered or preferentially populate alternative non-native motifs. We do not interpret transient loss of helicity alone as bias; instead, bias is identified when trajectories converge to persistent, low-entropy, non-native basins across replicates. By this criterion, helix-recovering force fields show balanced sampling, whereas strand-prone models exhibit systematic diversion from experimentally supported helicity.

#### Structured upon binding

These peptides are disordered in isolation and adopt defined conformations only when bound to a protein partner. Accordingly, extended-start simulations in water are not expected to converge to a single stable fold. Instead, the key diagnostic is whether force fields artificially stabilize persistent secondary-structure basins in the absence of binding.

For the MAT*α*2 chameleon peptide (1MNM), the 1 µs triplicates reveal divergent force-field tendencies (SI Fig. 43–49). ff99IDPs and ff14IDPs sample both helical and hairpin conformations transiently, without persistent stabilization. In contrast, ff19SB, ff14IDPs, and a99SBdisp show a preference for helical conformations, while OPLSIDPSFF and ff14IDPSFF favor *β*-hairpin formation. Notably, ff14IDPSFF exhibits reversible hairpin formation and loss, whereas OPLSIDPSFF stabilizes hairpin conformations once formed.

Extending simulations to 10 µs sharpens this distinction. OPLSIDPSFF maintains *β*-hairpin states without unfolding events, indicating a deeper strand-stabilizing basin. ff14IDPSFF continues to display reversible *β*-hairpin sampling, suggesting a shallower competing minimum. Helix-prone force fields intermittently sample *α*-structure but do not converge to persistent folded states. Overall, 1MNM highlights that certain force fields promote stable alternative folds in isolation, whereas others maintain a dynamically disordered ensemble.

For NRF2 (2FLU), the 1 µs triplicates again expose clear secondary-structure tendencies (SI Fig. 50–56). OPLSIDPSFF consistently stabilizes *β*-hairpin conformations distinct from the bound-state structure. In contrast, ff19SB, ff99SB, ff99IDPs, and ff14IDPSFF exhibit helical tendencies, with ff19SB and a99SBdisp sampling *α*-helical conformations at lower populations. The remaining force fields show limited persistent structure.

These qualitative behaviors persist in the 10 µs simulations. OPLSIDPSFF samples multiple distinct hairpin states with sustained populations, whereas helix-prone models sample *α*-structure intermittently without clear convergence. C36IDPSFF occasionally samples a native-like hairpin but only sparsely. Thus, 2FLU further illustrates how strand-biased force fields may overstabilize compact *β*-structures even when the peptide is expected to remain dynamic in water.

For 2N2F, which contains a central helix that is unstable in isolation, the 1 µs triplicates show minimal secondary-structure stabilization across force fields (SI Fig. 57–63). ff19SB samples native-like helical motifs at low population, while most other models predominantly explore disordered conformations. Extending to 10 µs does not alter this qualitative picture: structural heterogeneity dominates, and no force field stabilizes a persistent folded basin. Clustering reveals disordered ensembles across force fields, consistent with the expectation that this peptide requires binding interactions to access its native structure.

#### Solvent-sensitive helices

Peptides such as GLP-1 (1D0R), PAMP (2FLY), and phylloseptin-1 (2JQ0) adopt *α*-helical conformations in TFE or membrane-mimetic environments but are expected to remain largely disordered in pure water. In extended-start simulations performed in aqueous conditions, persistent stabilization of secondary structure therefore indicates potential over-structuring bias.

In the 1 µs triplicates, clear force-field tendencies emerge (SI Fig. 64–84). ff14IDPSFF and OPLSIDPSFF frequently stabilize *β*-hairpin-like motifs across all three peptides, indicating a recurring strand preference. To a lesser extent, C36IDPSFF and ff99IDPs show similar behavior. In contrast, ff19SB, des-amber, and des-amber-SF1.0 promote *α*-helical sampling in 1D0R and 2FLY, particularly in the C-terminal region. Although not expected, formation in this region maps with experimental observations showing that this segment retains helicity at lower cosolvent concentrations while other helical regions are lost. a99SBdisp also promotes helicity in 1D0R, as observed in multiple replicas. For 2FLY, ff99SB samples *β*-hairpin conformations more frequently than ff99IDPs, whereas a99SBdisp favors *α*-helical sampling, with des-amber and des-amber-SF1.0 showing weaker helical tendencies. In 2JQ0, secondary-structure formation is generally less pronounced, though strand-biased models still sample *β*-hairpins intermittently.

Extending simulations to 10 µs does not substantially alter these qualitative trends. Strand-prone force fields continue to stabilize compact *β*-like states with moderate persistence, while helix-promoting models intermittently populate *α*-helical segments without converging to long-lived folded basins. Across all three peptides, ensembles remain heterogeneous overall; however, the repeated emergence of *β*-hairpin motifs in multiple systems highlights a systematic strand bias in specific force fields.

Taken together, these solvent-sensitive peptides underscore how secondary-structure propensities encoded in a force field can reshape the conformational landscape even when disorder is experimentally expected. Over-stabilization of either *α*- or *β*-structure in this regime reflects imbalance in the underlying energy surface rather than successful folding behavior.

### Summary

Folding simulations highlight distinct tendencies among force fields (Fig. 3 and SI Table 2). OPLSIDPSFF displays a strong and consistent *β*-bias across systems, often producing dominant clusters regardless of peptide identity. ff19SB and ff99IDPs show context-sensitive behavior, folding miniproteins and moderately structured peptides while avoiding strong misfolding. IDP-optimized force fields vary widely in behavior: some (ff14IDPSFF) promote *β*-structure, while others (ff99IDPs) show flexibility and helix recovery. These simulations emphasize the need to assess force field performance across multiple structural regimes and not assume generality based on protein-focused benchmarks.

**Figure 3:**
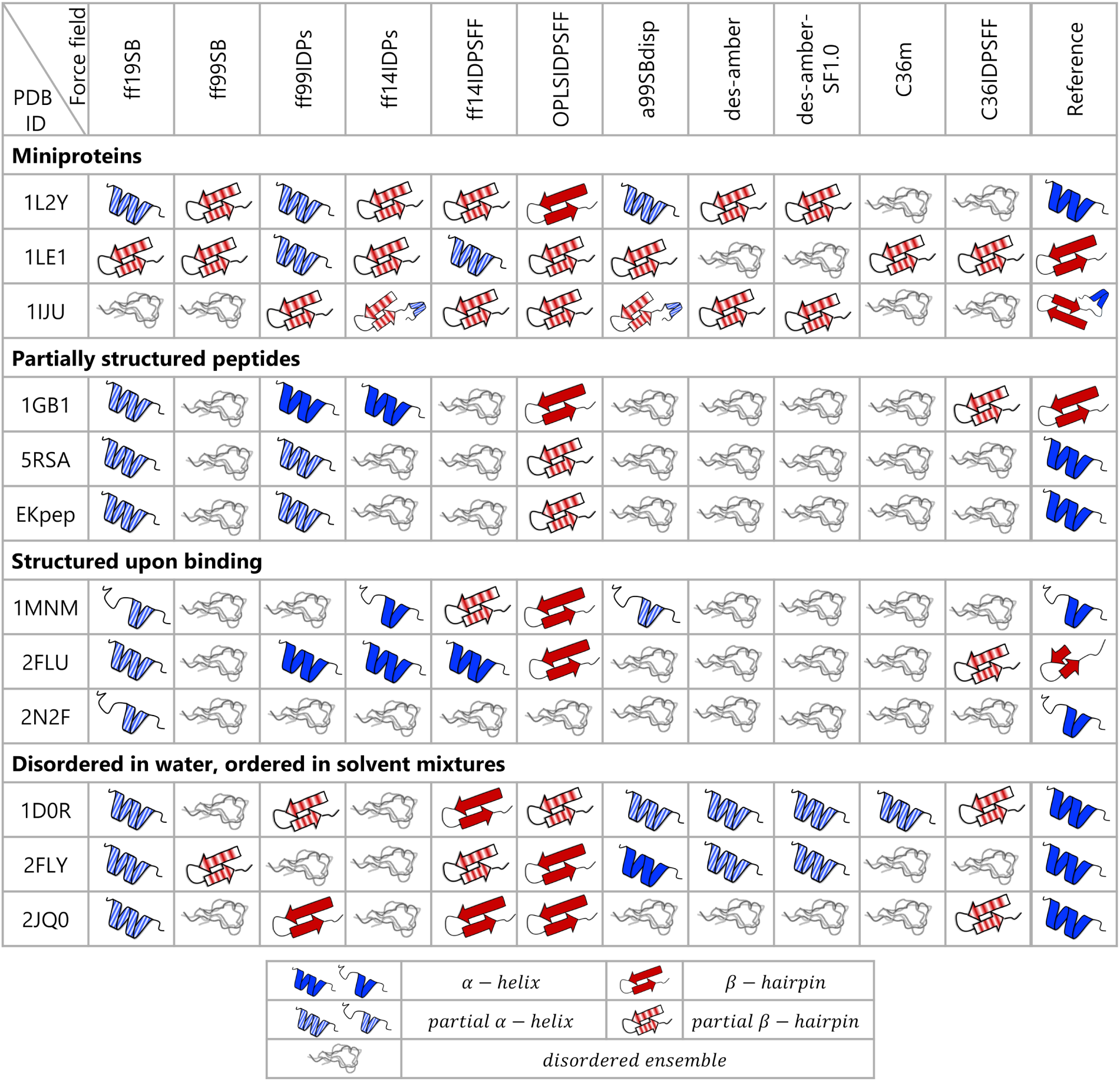
Summary of overall trends observed for each force field across different systems during 10 *µ*s simulations initiated from linear peptide sequences. Solid-colored helices or hairpins represent the formation of native secondary structures, while striped helices or hairpins indicate partial formation of native structures or stabilization of non-native secondary structures. Wavy lines denote disordered ensemble sampling. The “Reference” column shows the experimental PDB structure used as the starting model (free, complexed, or determined in mixed solvent).

Across nearly all peptides, OPLSIDPSFF shows a consistent *β*-hairpin preference, often early in the trajectory (e.g., 1GB1, EK, 1D0R), leading to dominant cluster populations (e.g., 1GB1: *>* 80%). In contrast, ff19SB and ff99IDPs intermittently recover native-like helices in marginal systems (e.g., EK, 5RSA), suggesting balanced behavior.

### Entropy as a proxy for conformational sampling across force fields

Entropy derived from population-weighted cluster occupancies using Shannon’s entropy provides a compact metric to compare the breadth and distribution of conformational space explored by different force fields. Because system size and trajectory length influence absolute entropy values, we analyze the 200 ns native-start, microsecond-extended, and 10 µs extended-start ensembles separately (Fig. 4 and SI Figs. 85–86).

**Figure 4:**
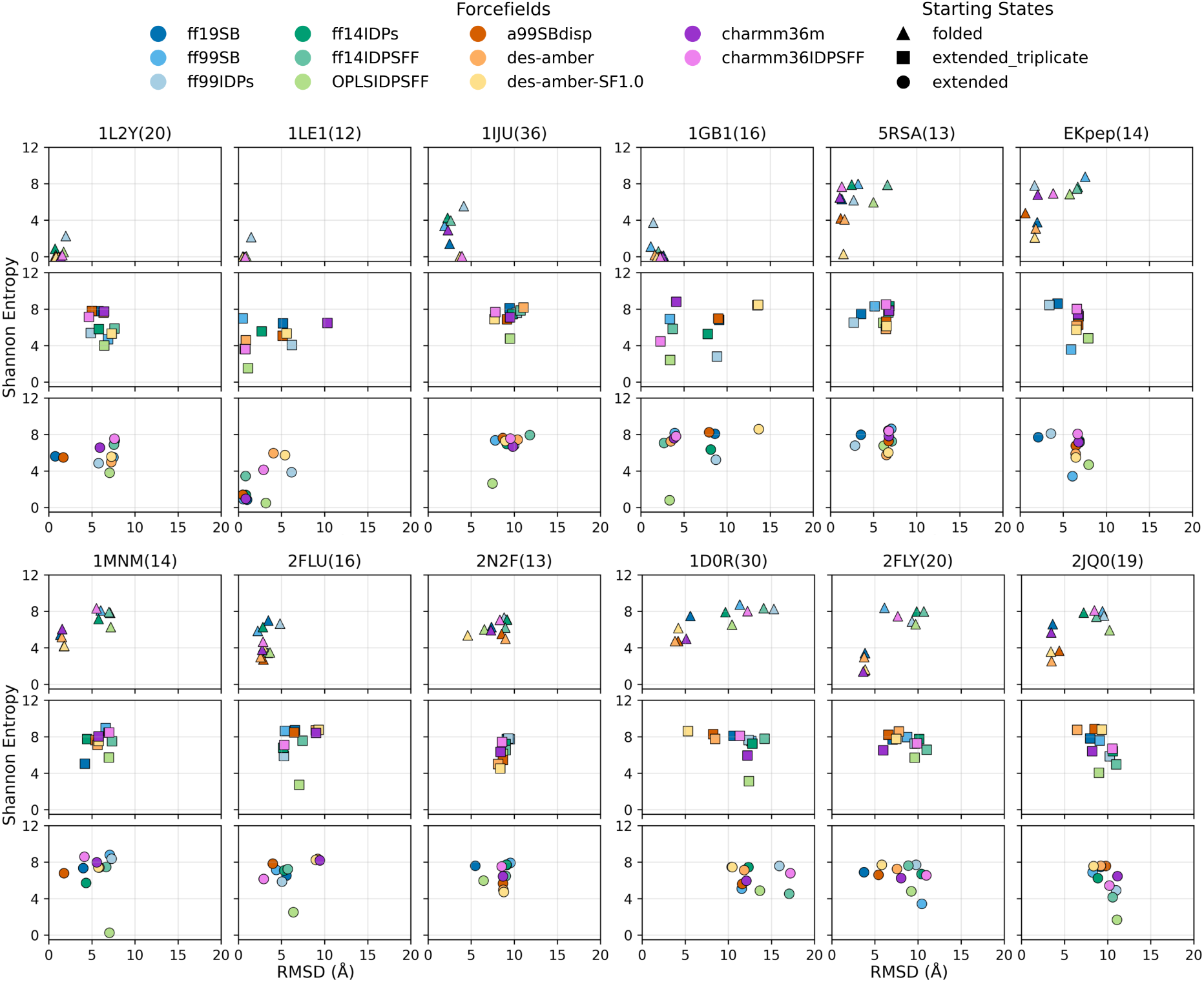
Relationship between Shannon entropy and RMSD across different peptide systems for each force field. For each system, triangle markers (top panels) represent data from five 200 ns simulations replicas initiated from experimentally folded structures, while square markers (middle panel) correspond to three 1 *µ*s simulations, and circle markers (bottom panels) correspond to 10 *µ*s simulations starting from linear peptide sequences. Numbers in parentheses indicate the length of each peptide sequence.

Our objective is not to assess exhaustive sampling, nor to compare entropy across simulation lengths. Rather, entropy quantifies whether a force field broadly explores conformational space (high entropy) or concentrates population into a limited number of states (low entropy). RMSD analysis complements this measure by determining whether low-entropy states correspond to experimentally supported structures or to alternative conformations indicative of bias.

Because the same clustering protocol, RMSD cutoff, and trajectory length are applied uniformly across all force fields within each simulation class, entropy serves here as a comparative diagnostic rather than an absolute thermodynamic quantity. While absolute values depend on clustering definitions, relative differences under identical analysis conditions primarily reflect differences in the underlying conformational distributions. This framework enables identification of force fields that systematically over-stabilize or over-disperse conformational ensembles relative to the consensus behavior.

We interpret the entropy-RMSD landscape (see Fig. 4) using three qualitative regimes:

1. Stable native basin (low entropy, low RMSD); 2) Non-native traps (low entropy, high RMSD); 3)Broad exploration (high entropy, any RMSD). Entropy values are not normalized across systems and should be interpreted within each system’s context.

#### Miniproteins (1L2Y, 1LE1, 1IJU)

In short simulations (200 ns) from native conformations, all three systems retain low RMSD (4Å) and low entropy (1.0) under most force fields. FF99IDPs consistently shows higher RMSD and entropy, suggesting destabilization. In long simulations from extended chains, only ff19SB and a99SBdisp achieve partial folding of Trp-cage (1L2Y) from extended conformations. In contrast, the Trpzip (1LE1) fold is readily sampled by several force fields, resulting in low RMSD (1.5Å) and low entropy (2.0) for several force fields. Interestingly, OPLSIDPSFF finds a non-native hairpin that is readily stabilized as seen by its lowest entropy, indicating that its *β*-bias might be unbalanced with respect to subtle sequence-dependent properties. None of the force fields successfully folds the larger *β*-defensin-1 (1IJU) within 10 *µ*s, where even the top clusters have very low population (SI Fig. 19). Examination of secondary structure timeseries (SI Fig. 18) reveals long-lived hairpins, particularly in OPLSIDPSFF, but as most of the system remains disordered, this leads to overall low population clusters. Interestingly, most of the secondary structure sampled is incompatible with native-like secondary structure preferences.

#### Partially structured peptides (1GB1, 5RSA, EKpep)

1GB1 remains stable when starting from the reference state (low entropy and RMSD). OPLSIDPSFF shows especially low entropy, consistent with its known hairpin preference. Approximately half of the force fields recover folded-like clusters from extended states. In contrast, 5RSA and EKpep show higher entropies even when RMSD remains low, indicating dynamic exchange between folded and unfolded states. Only ff99IDPs and ff19SB recover native-like clusters for these two systems starting from extended. Interestingly, neither of these two force fields was among those that fold 1GB1, suggesting there might be a slight bias towards helical states.

#### Structured upon binding (1MNM, 2FLU, 2N2F)

In native-start simulations, 1MNM and 2FLU retain native-like clusters but with moderate-to-high entropy, consistent with transient helicity in the absence of a binding partner. Few force fields succeed in refolding these systems from extended chains. For the chameleon 1MNM, OPLSIDPSFF and ff14IDPSFF favor the hairpin conformation, with OPLSIDPSFF sampling it at low entropy (contrasting with the high entropy in runs starting from the helical conformation). 2N2F remains disordered independently of the starting point, with high RMSD and moderate entropy, confirming that no dominant folded state is sampled. This contrasts with 1MNM and 2FLU supports a nuanced view of conformational selection versus induced fit. The latter two sample bound-like structures as top clusters (albeit with low populations and high entropy), implying that a complementary binding site could stabilize these conformations. In contrast, 2N2F appears to require interaction with its binding partner to access the native-like fold as the top state.

#### Solvent-sensitive helices (1D0R, 2FLY, 2JQO)

Native-start trajectories preserve helical structure to varying degrees (RMSD < 5Å) with moderate entropy. Surprisingly, extended-start simulations show lower entropy despite longer duration, often converging to compact, non-reference states (RMSD 6–10Å, entropy < 2) – with some force fields (ff19SB, charmm36m, des-amber, des-amber-SF1.0) stabilizing partial helices at one termini, with intrachain hydrophobic interactions collapsing the remainder of the sequence. These compact states are not unexpected: starting helices are stabilized in membrane-mimicking environments and expose hydrophobic side chains. In aqueous solvent, collapse likely reflects an entropic and enthalpic drive to shield hydrophobics from water, which in turn leads to low-entropy conformations that are structurally distinct from the input helix.

## Discussion

Peptides sit at the edge of stability: they lack the extensive tertiary contacts that stabilize globular proteins. Some peptides adopt stable structures under physiological conditions and are often classified as miniproteins. Others become structured only in specific contexts, such as in non-aqueous solvents, macromolecular crowding, or upon binding to a partner. This intermediate regime, between intrinsically disordered and stably folded, makes peptides a sensitive benchmark to detect and characterize force field biases. The peptides chosen for this study span this continuum, including well-folded systems, disordered peptides, and context-dependent examples.

As peptide therapeutics continue to gain traction in pharmaceutical research, it is critical to assess what improvements are needed in computational pipelines to reliably predict peptide binding. A useful decomposition of the binding free energy is:

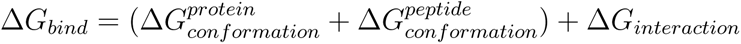

In practice, the protein’s conformational penalty between the apo and holo state 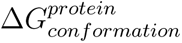 often cancels in relative binding comparisons (ΔΔ*G_bind_*) between two peptides. However, the peptide’s conformational free energy depends strongly on the force field’s ability to model its unbound and bound ensembles accurately. When two peptides are differentially affected by force field errors—such as differing helix propensities or hairpin lengths—the resulting Δ*G_bind_*can be significantly biased.

Our interest in this topic stems from efforts to predict peptide epitopes, their binding modes, and complex structures using data-guided simulations.^12,82,83^ In this context, sparse experimental data guide the peptide to approach the protein binding site (typically within ∼8 Å, avoiding diffusion times that hinder binding simulations), allowing the force field and sampling to determine the binding pose and whether the peptide folds upon binding. We observe a consistent trend where weaker binders (micromolar and millimolar) require stronger guiding data to reproduce expected binding poses, whereas nanomolar binders typically recover experimental binding modes with significantly less data. In low-affinity binders, where the energetic contributions from folding and binding are comparable, even minor force field biases may disrupt the delicate balance required for productive complex formation – requiring higher guiding power.^84^ Identifying such trends is important for going from qualitative relative binding affinities (e.g., from competitive binding simulations^82,85^) into more quantitative and robust measurements. The use of implicit solvent in previous methodologies inherits deficiencies from this force field selection. Thus, the current work helps users make better decisions for choosing force fields for transferring these methodologies into explicit solvent environments.

The force fields selected include both traditional parameterizations and more recent variants designed to better balance sampling of folded and disordered states. While modern biomolecular force fields are rarely parameterized directly against folded protein structures, they are commonly validated and iteratively refined to reproduce folded-protein observables (e.g., stability of benchmark folds, NMR observables, secondary-structure propensities). As a result, residual biases may persist when these models are applied to small peptides that sit near disorder–order boundaries. The peptide systems examined here are therefore well suited to amplify such residual biases.

Force field developers already benchmark their methods using metrics tailored to their development goals, often comparing simulated observables to experimental data such as NMR chemical shifts or scalar couplings,^86^ with agreement quantified via ^2^ statistics. Additionally, studies that compare across an array of force fields are usually intended to identify alternative metrics or benchmarks that support more informed force field selection choices.^28,87,88^ Such studies provide valuable insight into force field performance and underscore the complexity of assessing accuracy in molecular simulations.

One interesting note is that traditionally it has been easier to assess helicity than it is *β* propensity. Thus, many force field assessments have focused on whether the force field is too helical or not – looking at helical vs coil balance. However, it is entirely possible that a force field has either too much propensity to form both helices and strands,^22,89^ or too little of both of them compared to coil conformations.

Our paired simulation design (starting from native or extended) allows us to distinguish between a force field’s ability to preserve native conformations and its ability to discover them from unfolded states. Several force fields (e.g., ff14IDPSFF, OPLSIDPSFF) maintain hairpins once formed but fail to reach them from extended structures—suggesting shallow but narrow basins. Others (e.g., ff19SB, ff99IDPs) allow reversible transitions, consistent with broader landscape exploration.

RMSD alone cannot distinguish between realistic flexibility and biased trapping. By analyzing Shannon entropy over cluster populations, we reveal whether force fields sample broad ensembles (suggesting realistic disorder) or artificially narrow basins (indicative of structural bias). For example, OPLSIDPSFF often shows low entropy across peptides, consistent with early stabilization of compact, non-native basins.

Our results indicate that no single force field performs best across all systems. Nonetheless, clear trends emerge. Some force fields exhibit a strong bias toward specific secondary structures, stabilizing helices or strands even when these are not experimentally favored. Others tend to sample transient structure without stabilizing it long enough to be biologically relevant. A third group largely suppresses structure formation, even in peptides expected to fold. These behaviors are informative of underlying biases and limitations.

In our simulations, OPLSIDPSFF consistently promotes *β*-hairpins even in systems like EK or GLP-1 that are not *β*-structured, highlighting a consistent strand bias. In contrast, ff99IDPs frequently destabilizes helices, even in stable miniproteins like Trp-cage, and occasionally introduces *β*-structure in disordered or helical systems. ff19SB and a99SBdisp, by contrast, maintain expected secondary structure in folded systems and allow context-sensitive folding in marginal peptides. Notably, force fields optimized for disordered ensembles do not consistently outperform traditional models. For example, ff14IDPSFF introduces non-native *β*-hairpins in several peptides, while OPLSIDPSFF frequently stabilizes strand-pairing even in helically biased systems. These results caution against assuming that IDP-tuned force fields generalize to all flexible peptide systems.

Based on stability, folding, and entropy behavior across twelve peptides, ff19SB and a99SBdisp consistently show the most balanced performance—preserving native folds, avoiding over-stabilization, and sampling relevant ensembles. These force fields represent a strong starting point for modeling flexible peptides. Ongoing improvements in force field design, expanded benchmark sets, and longer simulations, combined with FAIR (Findable, Accessible, Interoperable, Reproducible) principles, offer new opportunities to detect and correct structural biases. Our benchmark shows that peptides, as sensitive probes between order and disorder, expose biases not always evident in globular protein models. We advocate for routine inclusion of diverse peptide benchmarks—covering miniproteins, disordered sequences, and context-dependent folders—in future evaluations.

## Conclusion

Peptides inhabit a structurally diverse and biologically significant regime that challenges current molecular mechanics force fields. Our systematic benchmark, combining short simulations from native structures and long folding trajectories from extended chains, reveals that while many force fields are capable of maintaining folded states in select systems, few are unbiased across the full peptide landscape. Force fields such as ff19SB and a99SBdisp strike a useful balance—retaining native structure in well-folded systems while allowing transitions in marginal or context-sensitive peptides. Others, such as ff99IDPs or OPLSIDPSFF, show clear tendencies toward particular secondary structures, often stabilizing non-native folds.

These findings have important implications for peptide modeling in drug discovery, where Δ*G* predictions depend not only on interaction energies but also on accurate modeling of conformational ensembles. When comparing different peptides, force field-dependent errors in the conformational component of Δ*G* can bias binding predictions. Our results underscore the need for caution when interpreting peptide simulations and highlight the value of ensemble-based and comparative strategies.

We anticipate that this benchmark will serve both as a practical resource for force field users and as a testbed for future force field development targeting intrinsically disordered and conformationally adaptive systems. The dataset, simulation protocols, and analysis framework are available at https://github.com/PDNALab/peptFF-benchmark to facilitate further refinement and standardization in the community.

## Acknowledgement

The authors thank the support from the National Institutes of Health grant R01GM149646.

## Supporting Information Available

Additional Figures can be found in the Supplementary information.

Summary of structural stability and folding behavior from 200 ns and 10 *µ*s simulations (Tables S1, S2); miniprotein analyses, including Ramachandran plots, per-residue secondary structure time series (200 ns, 1 *µ*s, 10 *µ*s), clustering with representative structures, per-residue secondary structure preferences, and 2 *µ*s interval-based secondary structure analysis (Figures S1–S21); analogous analyses for partially structured peptides (Figures S22–S42), peptides structured upon binding (Figures S43–S63), and peptides disordered in water but ordered in solvent mixtures (Figures S64–S84); Shannon entropy analysis and log-transformed cluster size distributions for each force field and system (Figures S85, S86)

## TOC Graphic

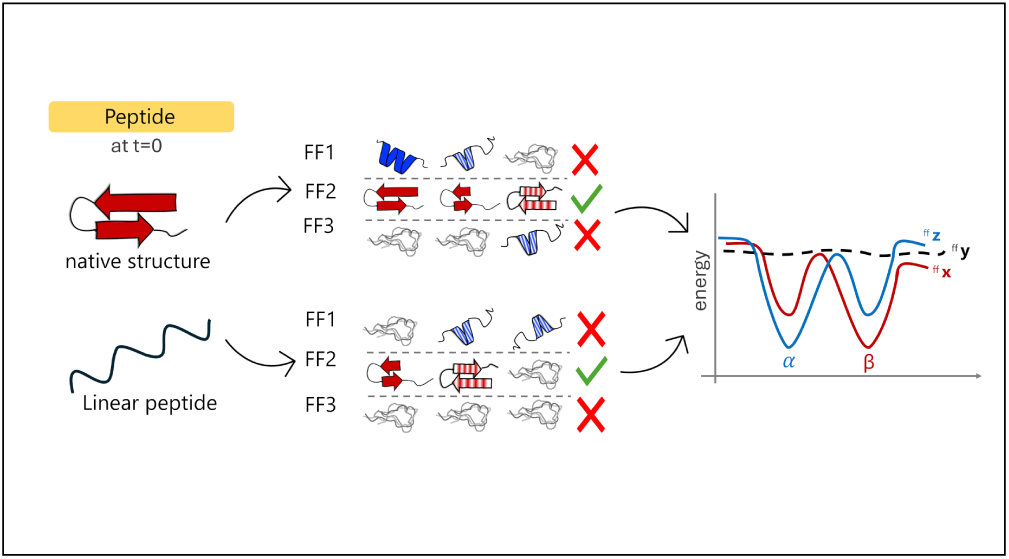

