## Supplementary material for "How Well Do Molecular Dynamics Force Fields Model Peptides? A Systematic Benchmark Across Diverse Folding Behaviors": SI Figures

Table S1: Summary of structural stability observed in 200 ns simulations initiated from folded peptide conformations. Simulations reveal force field-dependent tendencies to preserve or disrupt native-like structures.

| Peptide | Expected Structure | Retention of Expected Structure | Force Field Deviation |
|---|---|---|---|
| **Miniproteins** | | | |
| 1L2Y (Trp-cage) | Helix (>95%) | Structure preserved in most FFs throughout the simulation | Helicity disrupted in ff99IDPs, ff14IDPSFF, and OPLSIDPSFF |
| 1LE1 (Trpzip) | $\beta$-hairpin (>95%) | Hairpin maintained in all FFs except ff99IDPs | ff99IDPs fails to retain native fold; shows partial helix |
| 1IJU ($\beta$-defensin-1) | Folded $\beta$-structure | Most FFs maintain fold; secondary elements persist | ff99IDPs, ff14IDPs (at C-terminal), ff14IDPSFF (at N-terminal) and C36m/C36IDPSFF ( at N-terminal) show partial unfolding |
| **Partially Structured Peptides** | | | |
| 1GB1 (Protein G) | $\beta$-hairpin ($\sim$20%) | Hairpin preserved across all FFs over 200 ns | Minor loss in ff99IDPs; others maintain structure |
| 5RSA (C-peptide) | Partial helix | Refolding observed in ff99IDPs and a99SBdisp; partial helix retention in des-amber, C36IDPSFF, ff19SB | Helicity maintained in des-amber-SF1.0; C36m and OPLSIDPSFF favors $\beta$-structure |
| EK peptide | Helix ($\sim$40%) | Helix stabilized in a99SBdisp, des-amber, des-amber-SF1.0, ff19SB; partial refolding in ff99IDPs, C36IDPSFF | OPLSIDPSFF shifts toward $\beta$-structure |
| **Structured Upon Binding** | | | |
| 1MNM (MAT$\alpha$2) | Helix/hairpin (context-dependent) | Helical content preserved in ff19SB, a99SBdisp, des-amber, des-amber-SF1.0,C36m | Helix lost in other FFs; ff99IDPs and ff14IDPs show brief recovery; hairpin-like motifs were sampled by ff14IDPSFF and OPLSIDPSFF |
| 2FLU (Nrf2) | $\beta$-hairpin (binding-induced) | Native fold not retained in any FF | OPLSIDPSFF, des-amber, des-amber-SF1.0, C36m, and C36IDPSFF show weak $\beta$-bias; ff99IDPs favors helix instead |
| 2N2F (Dynorphin) | Helical motif (binding-induced) | Helical content transiently recovered in ff19SB, des-amber, and des-amber-SF1.0 | All other FFs rapidly disrupt helicity |
| **Disordered in Water** | | | |
| 1D0R (GLP-1) | Disordered ensemble | Helicity maintained in ff19SB, a99SBdisp, des-amber, des-amber-SF1.0, and C36m | C36IDPSFF shows partial helicity; IDP FFs and ff99SB disrupt helicity |
| 2FLY (PAMP) | Disordered ensemble | Helix retained in ff19SB, a99SB, des-amber, des-amber-SF1.0, and C36m | IDP FFs sample unfolded or $\beta$-hairpin states; ff99SB and C36IDPSFF disrupts helicity |
| 2JQ0 (Phylloseptin-1) | Disordered ensemble | Helicity largely maintained in ff19SB, a99SB, des-amber, and C36m (except one des-amber-SF1.0 replica) | ff99SB and IDP FFs favor unfolded or $\beta$-hairpin states |

Table S2: Summary of structural behavior in 10 µs simulations starting from extended peptide conformations. Convergence to native-like states reflects force field accuracy, while stabilization of alternative or non-native folds suggests potential force field bias.

| Peptide | Expected Structure | Retention of Expected Structure | Force Field Deviation |
|---|---|---|---|
| **Miniproteins** | | | |
| 1L2Y (Trp-cage) | Helix-forming | Only ff19SB and a99SBdisp sample helical structure; ff99IDPs shows partial helix | Most FFs fail to form helices; OPLSIDPSFF favors $\beta$-hairpins; ff14IDPSFF, des-amber and des-amber-SF1.0 partially form $\beta$-hairpins |
| 1LE1 (Trpzip) | Fast folding $\beta$-hairpin | Folded by 5 FFs; 2 FFs achieve partial hairpin structures | ff99IDPs forms partial helix; ff14IDPSFF transitions from hairpin to helix; des-amber and des-amber-SF1.0 show disordered behavior |
| 1IJU ($\beta$-defensin-1) | Complex $\beta$-fold | No FF fully recovers native fold; partial non-native $\beta$-hairpins sampled by several FFs | ff14IDPs partially folds the protein; OPLSIDPSFF stabilizes non-native 3-strand $\beta$-sheet |
| **Partially Structured Peptides** | | | |
| 1GB1 (Protein G) | Weak $\beta$-hairpin (∼20%) | OPLSIDPSFF folds early into dominant hairpin cluster | a99SBdisp, ff19SB folds partially towards helix; ff99IDPSFF and ff14IDPs show folding-unfolding-refolding helix dynamics |
| 5RSA (C-peptide) | Partial helix | Only ff19SB and ff99IDPs stabilize helical segments | OPLSIDPSFF favors $\beta$-hairpin formation |
| EK peptide | Moderate helicity (∼40%) | ff99SB, ff19SB and ff99IDPs retain helical segments | OPLSIDPSFF favors $\beta$-hairpin structures |
| **Structured Upon Binding** | | | |
| 1MNM (Chameleon) | Contextual helix/hairpin duality | Most FFs prefer either helix or hairpin; ff99SB and ff99IDPs transiently sample both | OPLS-IDPSFF favors $\beta$-hairpins |
| 2FLU (Nrf2) | Binding-induced $\beta$-hairpin | Only OPLSIDPSFF stabilizes hairpin; C36IDPsFF favors partial $\beta$-hairpin | ff19SB, ff99IDPs, ff14IDPs, and ff14IDPSFF show helix-biased behavior |
| 2N2F (Dynorphin) | Binding-induced helix | Only ff19SB samples helical structure | All other FFs show low secondary structure content |
| **Disordered in Water** | | | |
| 1D0R (GLP-1) | Coil in water; helix in TFE | Most FFs maintain disordered ensemble; des-amber briefly samples helix before unfolding; ff99IDPs and C36IDPs forms $\beta$-hairpin but destabilize | ff19SB and des-amber-SF1.0 stabilize C-terminal helices; ff14IDPSFF and OPLSIDPSFF favor $\beta$-hairpins |
| 2FLY (PAMP) | Coil in water; helix in TFE | Most FFs retain disordered conformations | ff19SB, a99SBdisp and des-amber-SF1.0 retain sample helices; ff99SB, ff14IDPSFF, OPLSIDPSFF, and C36IDPSFF favor $\beta$-hairpins |
| 2JQ0 (Phylloseptin-1) | Coil in water; helix in TFE | Most FFs retain disordered conformations | ff99IDPs, ff14IDPSFF, and OPLSIDPSFF form $\beta$-hairpins; C36IDPSFF forms $\beta$-hairpins partially; ff19SB shows partial helix formation |

### Miniproteins

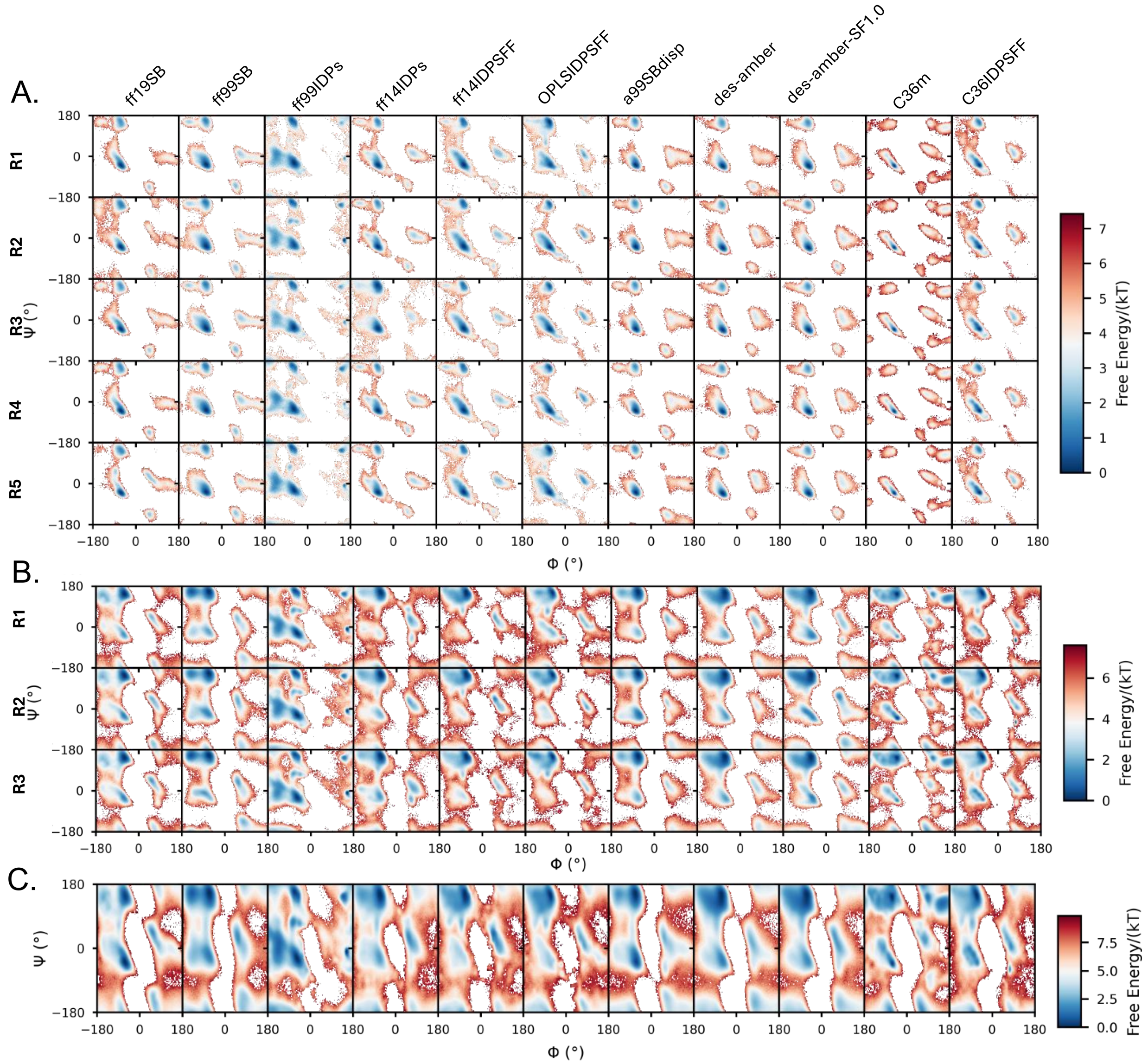


Figure S1: **Ramachandran plots for Trp-cage (PDB ID: 1L2Y)**. (A) Five 200-ns simulation replicas initiated from the experimentally folded structure. (B) Three 1-μs simulation replicas initiated from the linear peptide sequence. (C) 10-μs simulation initiated from the linear peptide sequence.

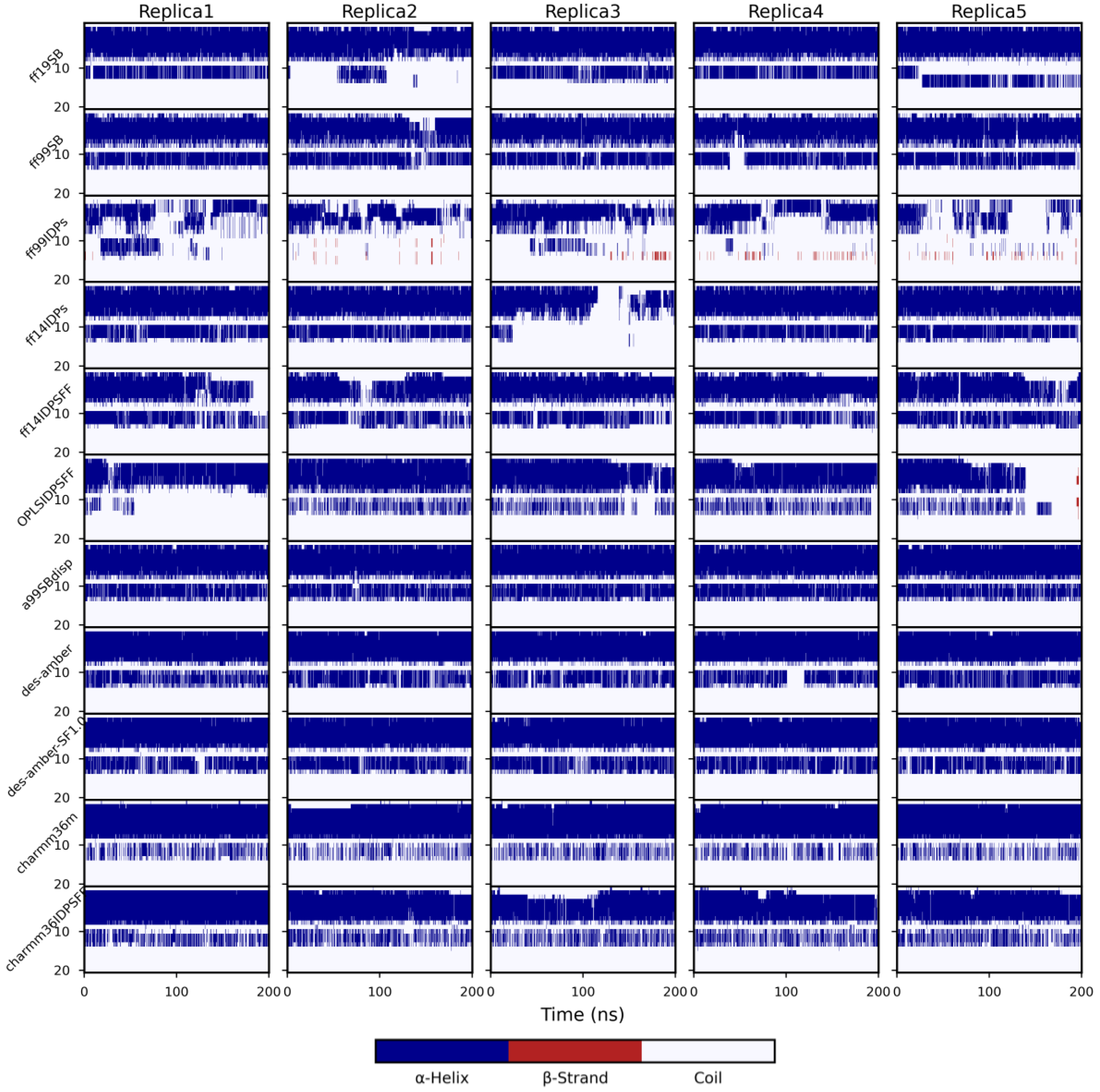


Figure S2: **Per-residue secondary structure time series for Trp-cage (PDB ID: 1L2Y)** for five 200-ns simulation replicas initiated from the folded structure. Red denotes $\beta$-strand, and blue denotes $\alpha$-helix or 3-10 helix, and the absence of color corresponds to coil or turn regions.

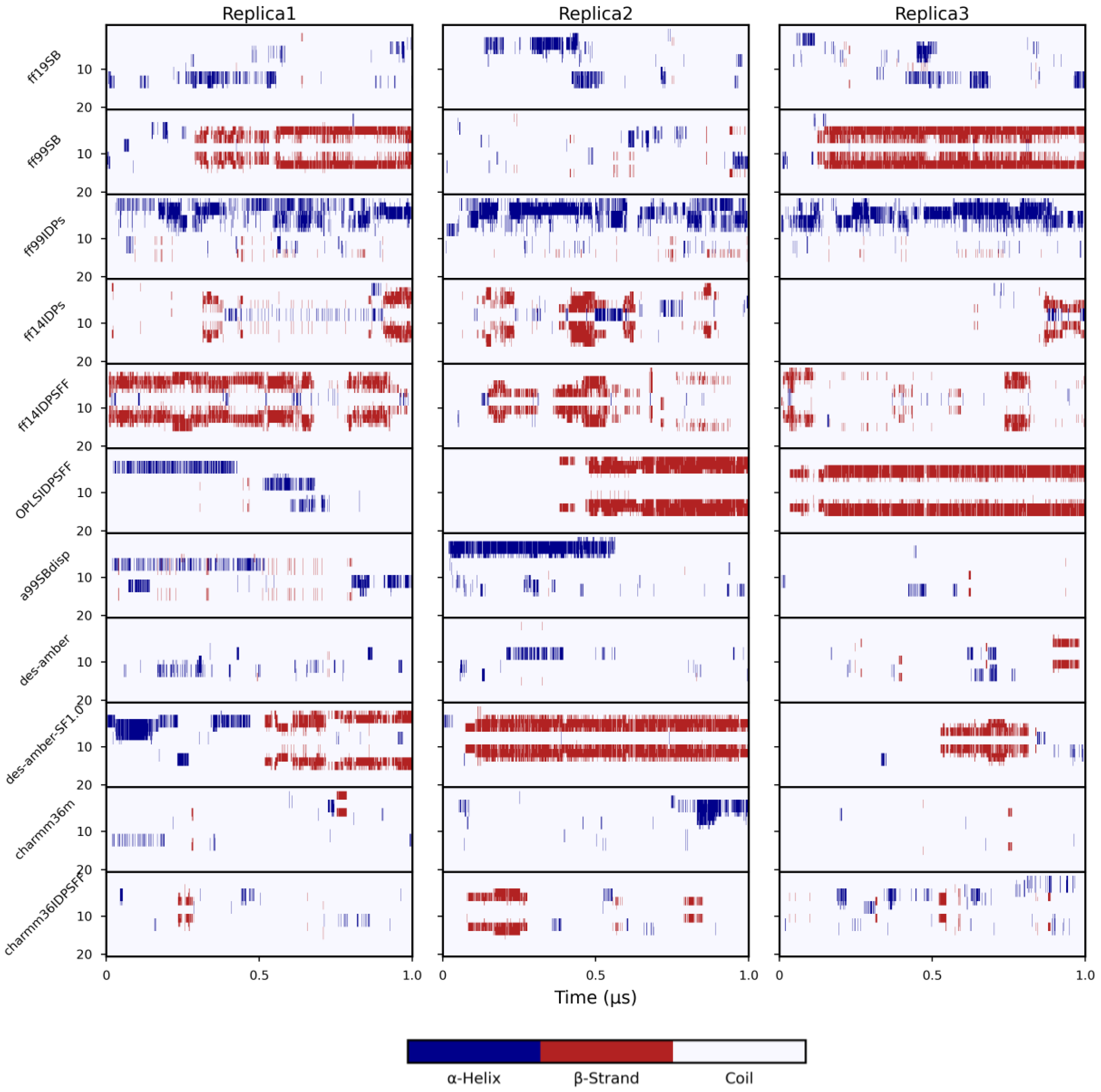


Figure S3: **Per-residue secondary structure time series for Trp-cage (PDB ID: 1L2Y)** for three 1 $\mu$s simulation replicas initiated from the folded structure. Red denotes $\beta$-strand, and blue denotes $\alpha$-helix or 3-10 helix, and the absence of color corresponds to coil or turn regions.

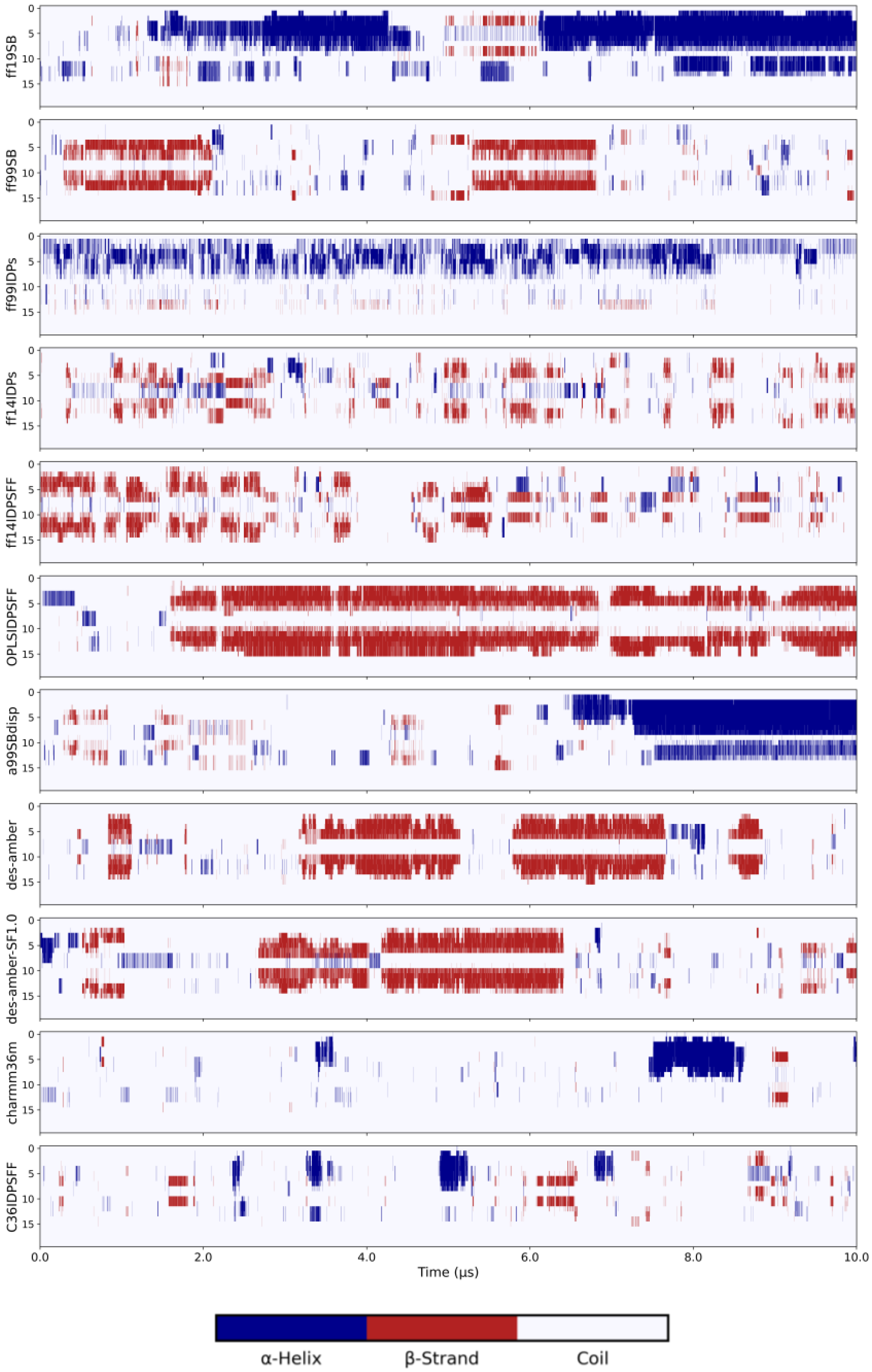


Figure S4: **Per-residue secondary structure time series for Trp-cage (PDB ID: 1L2Y)** for the 10 $\mu$s simulation replicas initiated from the folded structure. Red denotes $\beta$-strand, and blue denotes $\alpha$-helix or 3-10 helix, and the absence of color corresponds to coil or turn regions.

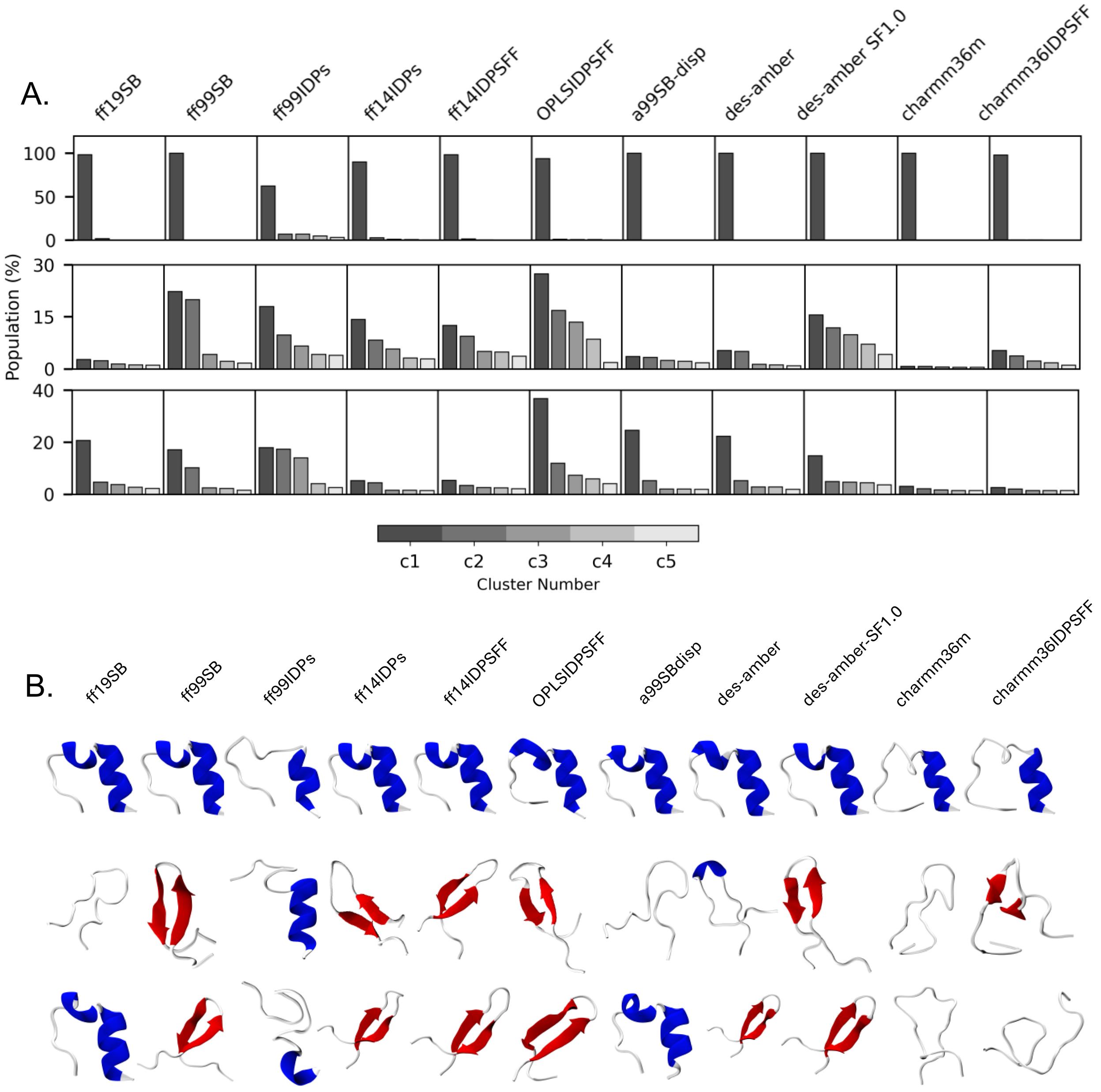


Figure S5: **Clustering results showing (A) cluster populations and (B) representative structures (centroids) of the most populated cluster for Trp-cage (PDB ID: 1L2Y).** The top panel shows clustering from five 200-ns stability simulations initiated from the folded structure; the middle panel shows clustering from three 1-µs simulation replicas initiated from the linear peptide sequence; and the bottom panel shows clustering from a 10-µs folding simulation initiated from the extended peptide conformation. Red indicates $\beta$-strand, blue indicates $\alpha$-helix or 3-10 helix, and white indicates coil or turn regions.

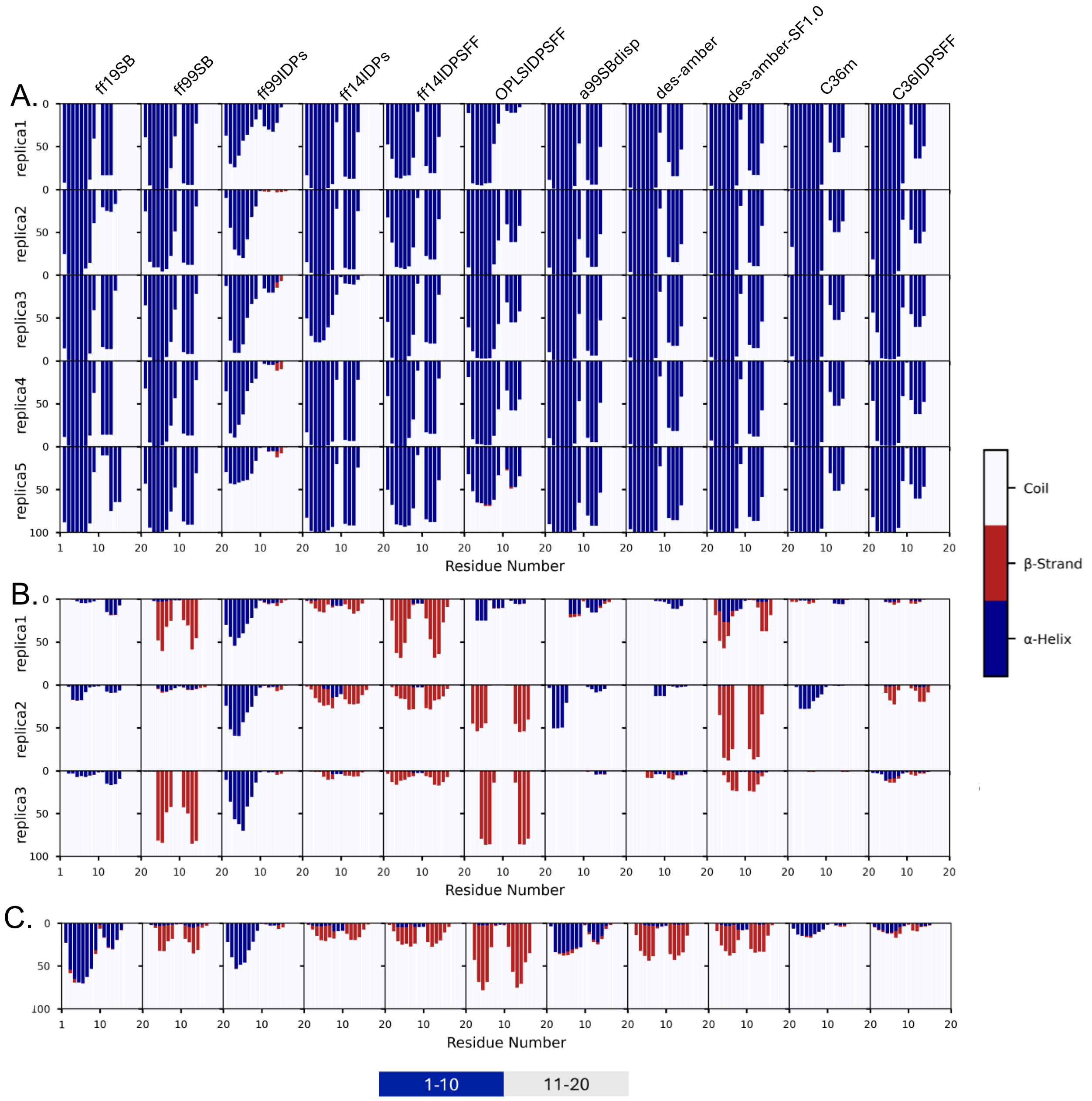


Figure S6: **Per-residue secondary structure preferences for Trp-cage (PDB ID: 1L2Y)** across simulations. Bars represent the percentage occurrence of each secondary structure type per residue in (A) five 200-ns replicas, (B) three 1-µs replicas, and (C) a 10-µs simulation. The secondary structure assignment in the experimentally folded structure is shown for reference. Red denotes $\beta$-strand, and blue denotes $\alpha$-helix or 3-10 helix, and the absence of color corresponds to coil or turn regions.

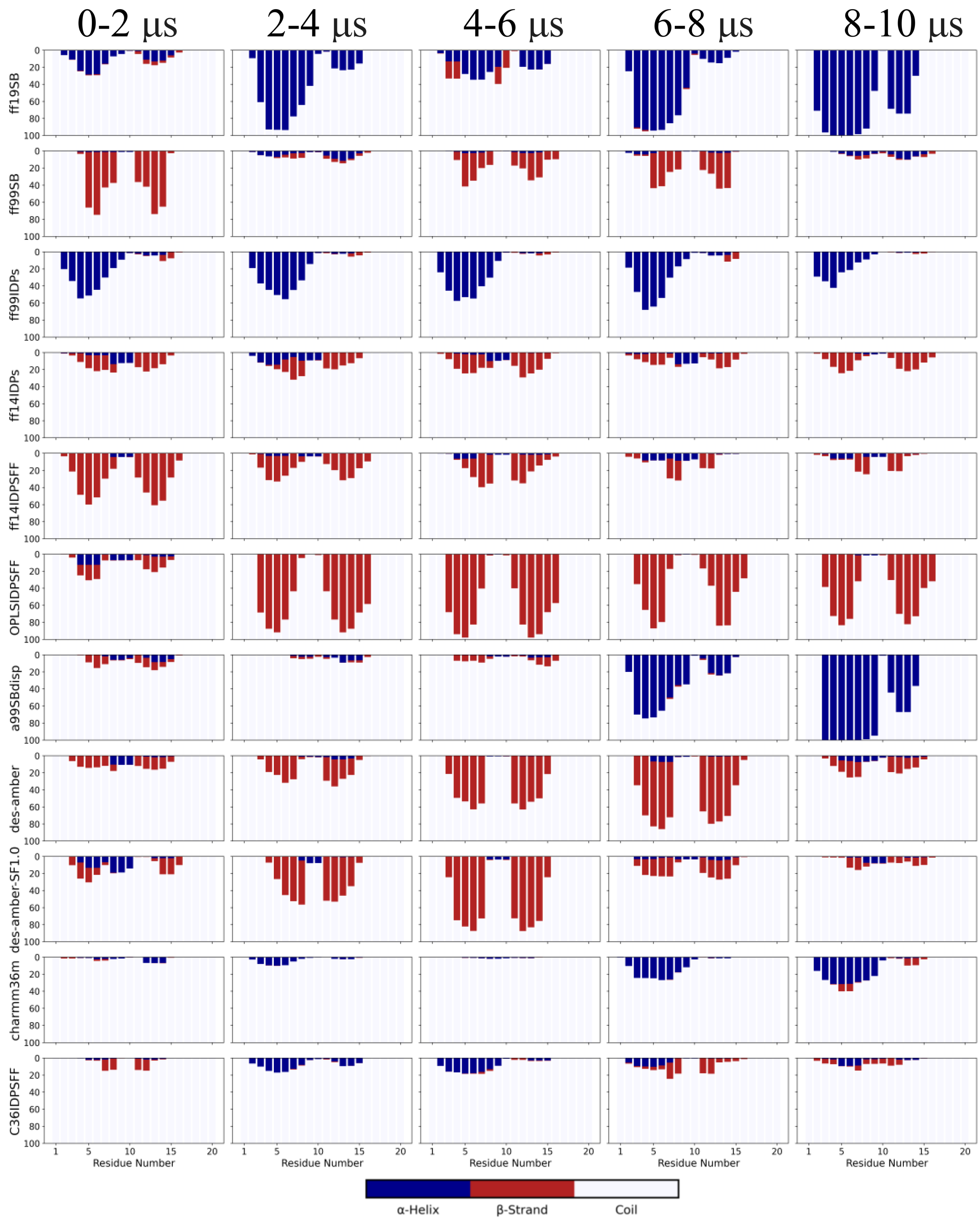


Figure S7: **Per residue secondary structure preferences for Trp-cage (PDB ID: 1L2Y) during the 10 $\mu$s simulation, shown as blocks of 2 $\mu$s intervals.** Bars represent the percentage occurrence of each secondary structure type per residue. Red indicates $\beta$-strand, blue indicates $\alpha$-helix or 3-10 helix, and the absence of color corresponds to coil or turn regions.

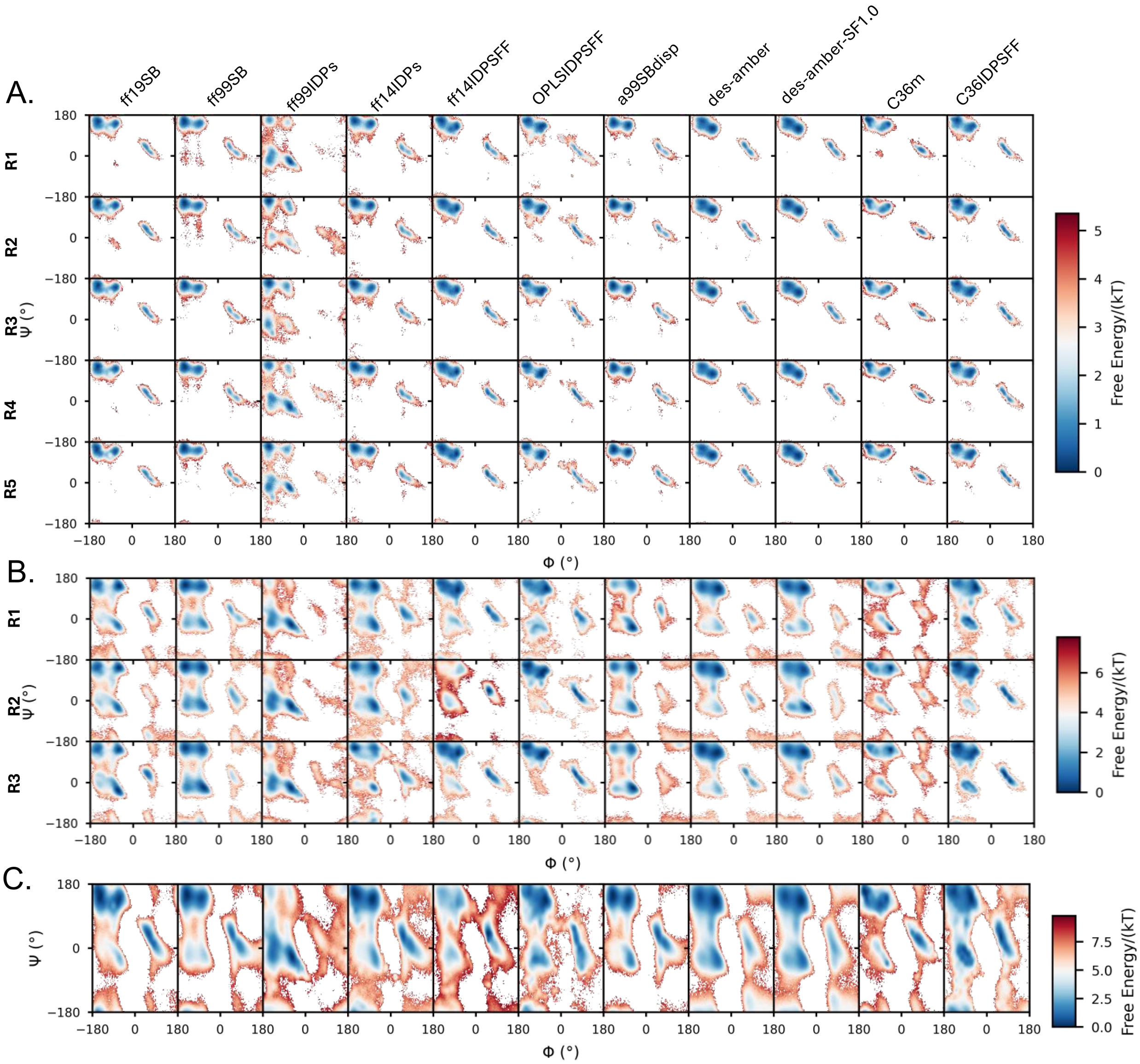


Figure S8: **Ramachandran plots for Trp-zip (PDB ID: 1LE1)**. (A) Five 200-ns simulation replicas initiated from the experimentally folded structure. (B) Three 1-µs simulation replicas initiated from the linear peptide sequence. (C) 10-µs simulation initiated from the linear peptide sequence.

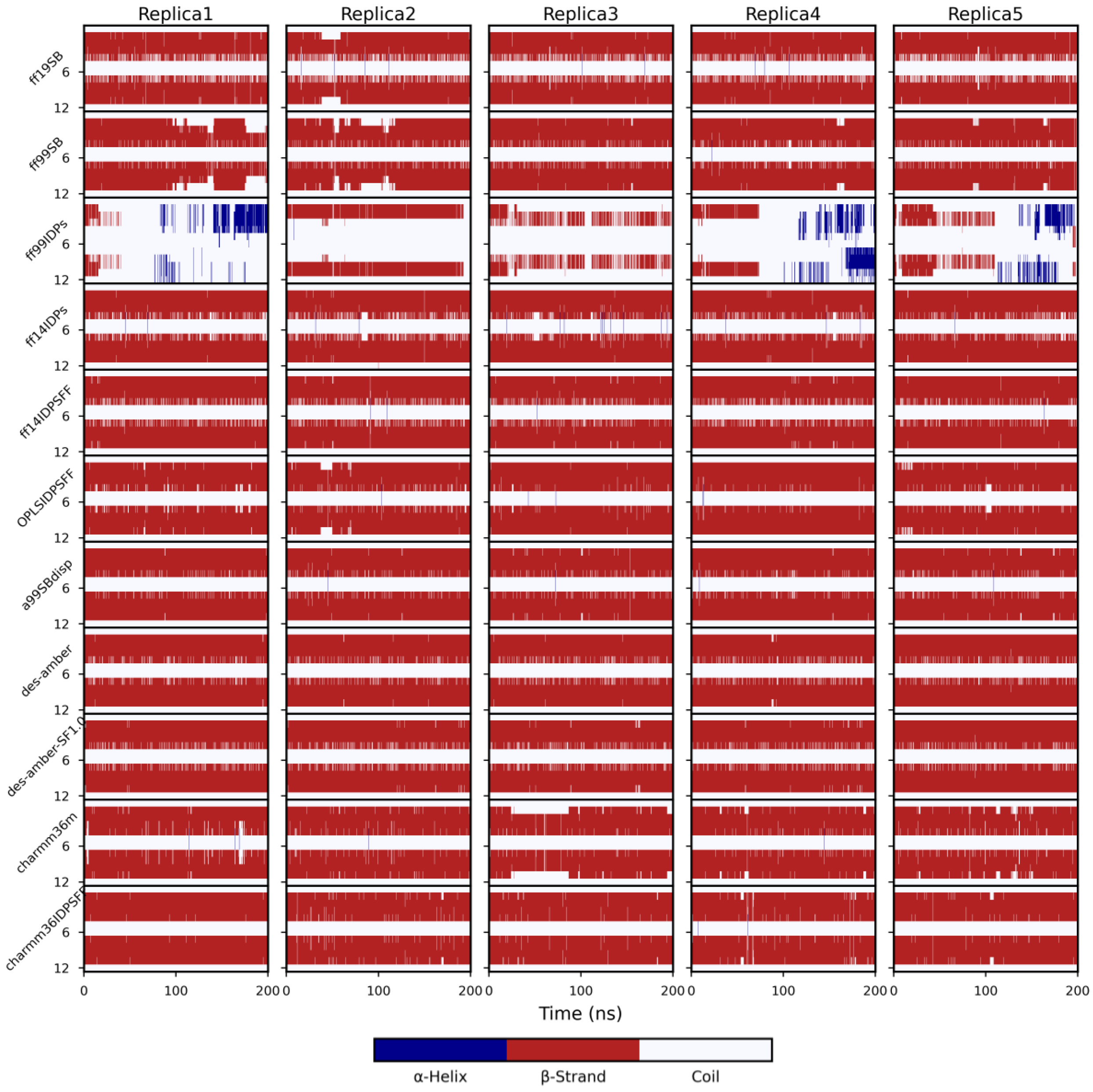


Figure S9: **Per-residue secondary structure time series for Trp-zip (PDB ID: 1LE1)** for five 200-ns simulation replicas initiated from the folded structure. Red denotes $\beta$-strand, and blue denotes $\alpha$-helix or 3-10 helix, and the absence of color corresponds to coil or turn regions.

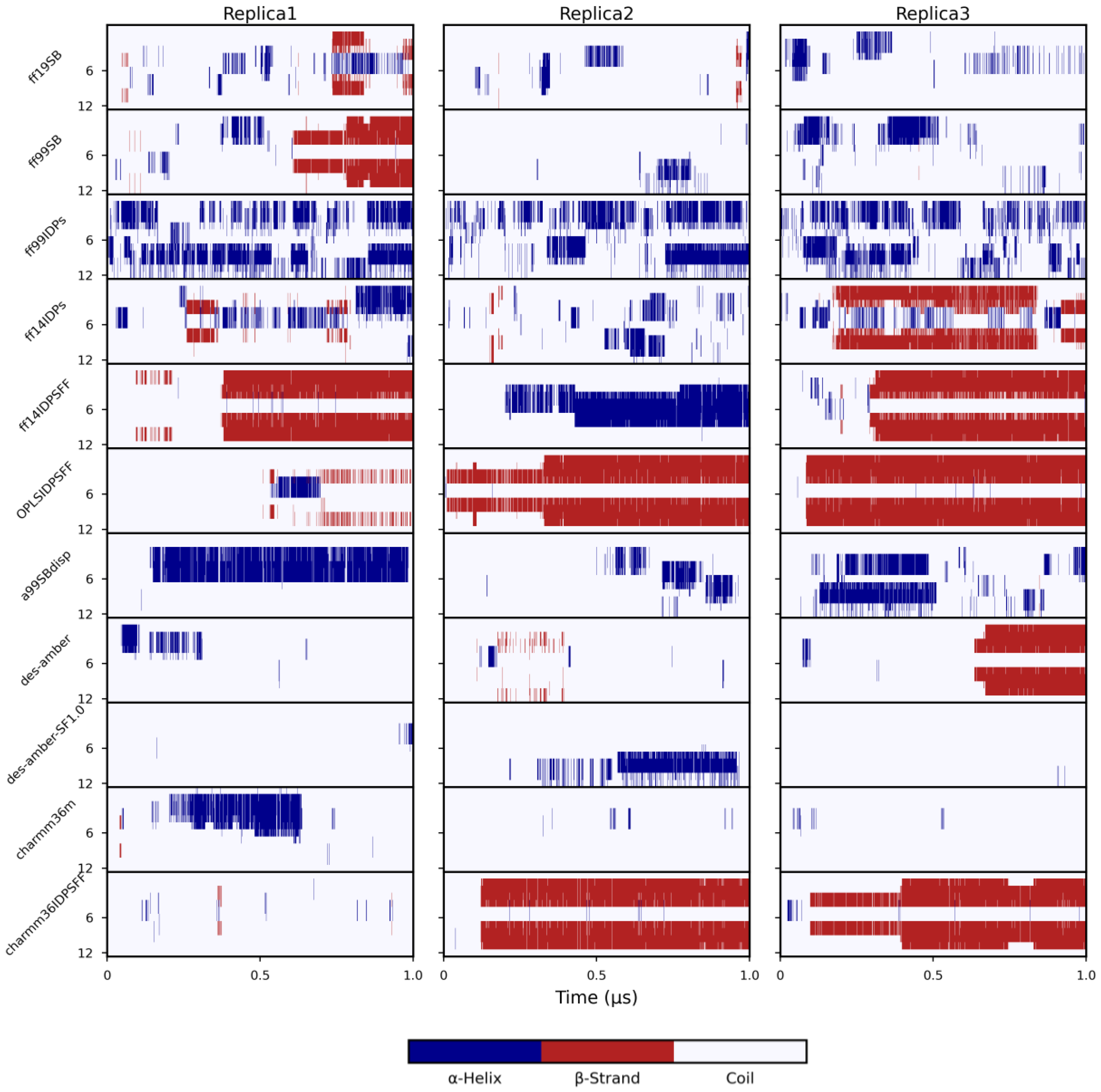


Figure S10: **Per-residue secondary structure time series for Trp-zip (PDB ID: 1LE1)** for three 1 $\mu$s simulation replicas initiated from the folded structure. Red denotes $\beta$-strand, and blue denotes $\alpha$-helix or 3-10 helix, and the absence of color corresponds to coil or turn regions.

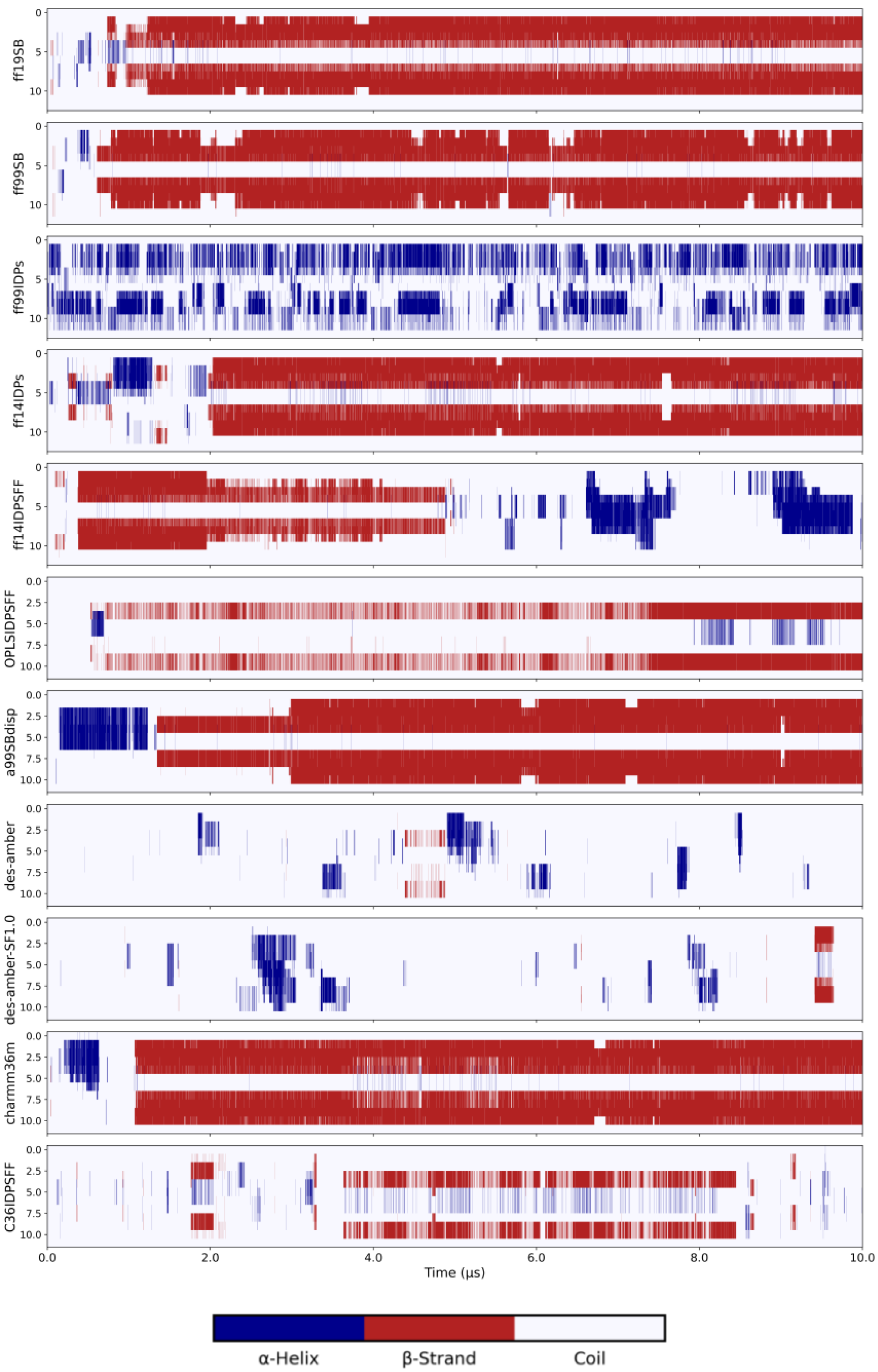


Figure S11: **Per-residue secondary structure time series for Trp-zip (PDB ID: 1LE1)** for the 10 $\mu$s simulation replicas initiated from the folded structure. Red denotes $\beta$-strand, and blue denotes $\alpha$-helix or 3-10 helix, and the absence of color corresponds to coil or turn regions.

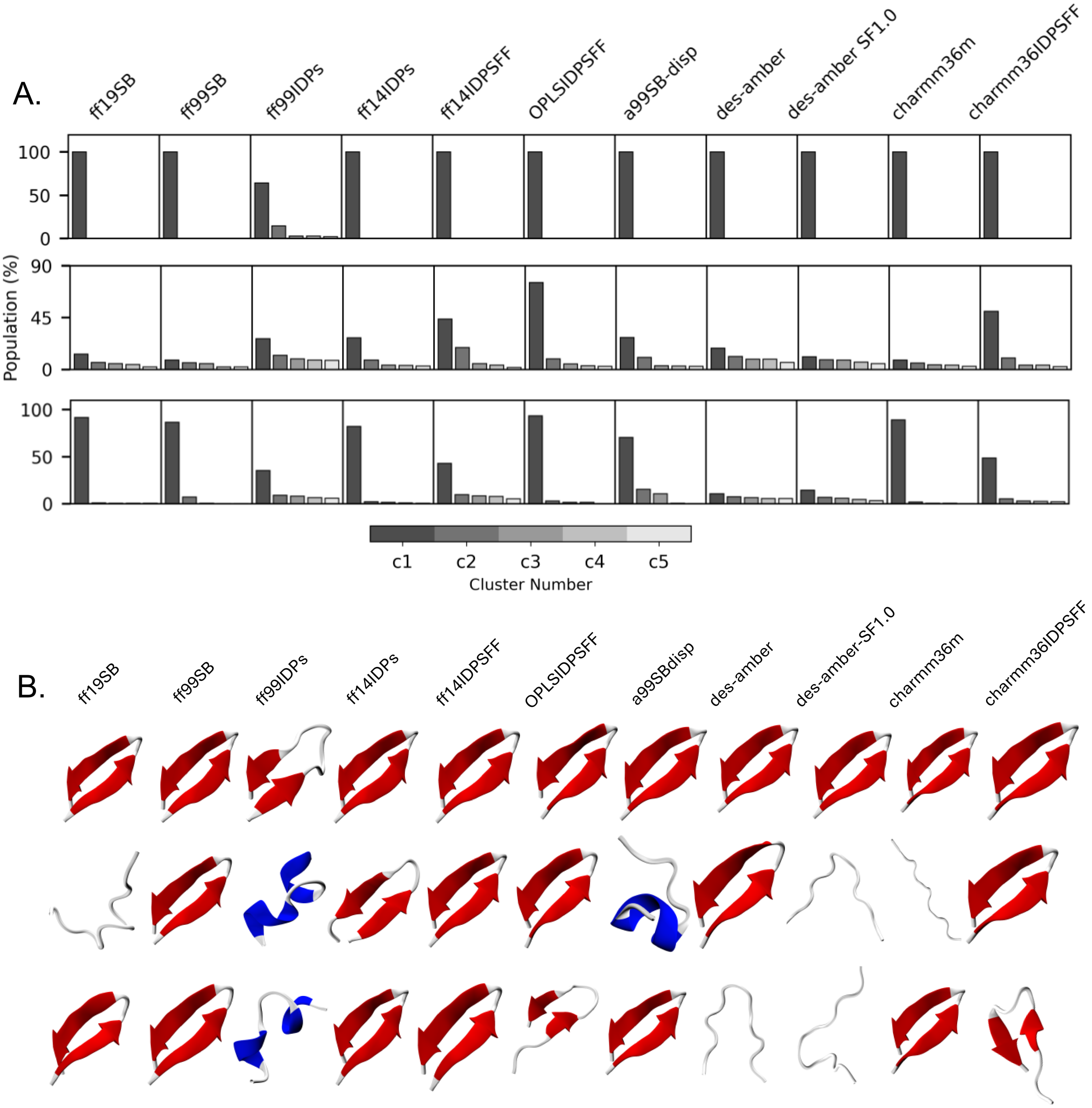


Figure S12: **Clustering results showing (A) cluster populations and (B) representative structures (centroids) of the most populated cluster for Trp-zip (PDB ID: 1LE1).** The top panel shows clustering from five 200-ns stability simulations initiated from the folded structure; the middle panel shows clustering from three 1-µs simulation replicas initiated from the linear peptide sequence; and the bottom panel shows clustering from a 10-µs folding simulation initiated from the extended peptide conformation. Red indicates $\beta$-strand, blue indicates $\alpha$-helix or 3-10 helix, and white indicates coil or turn regions.

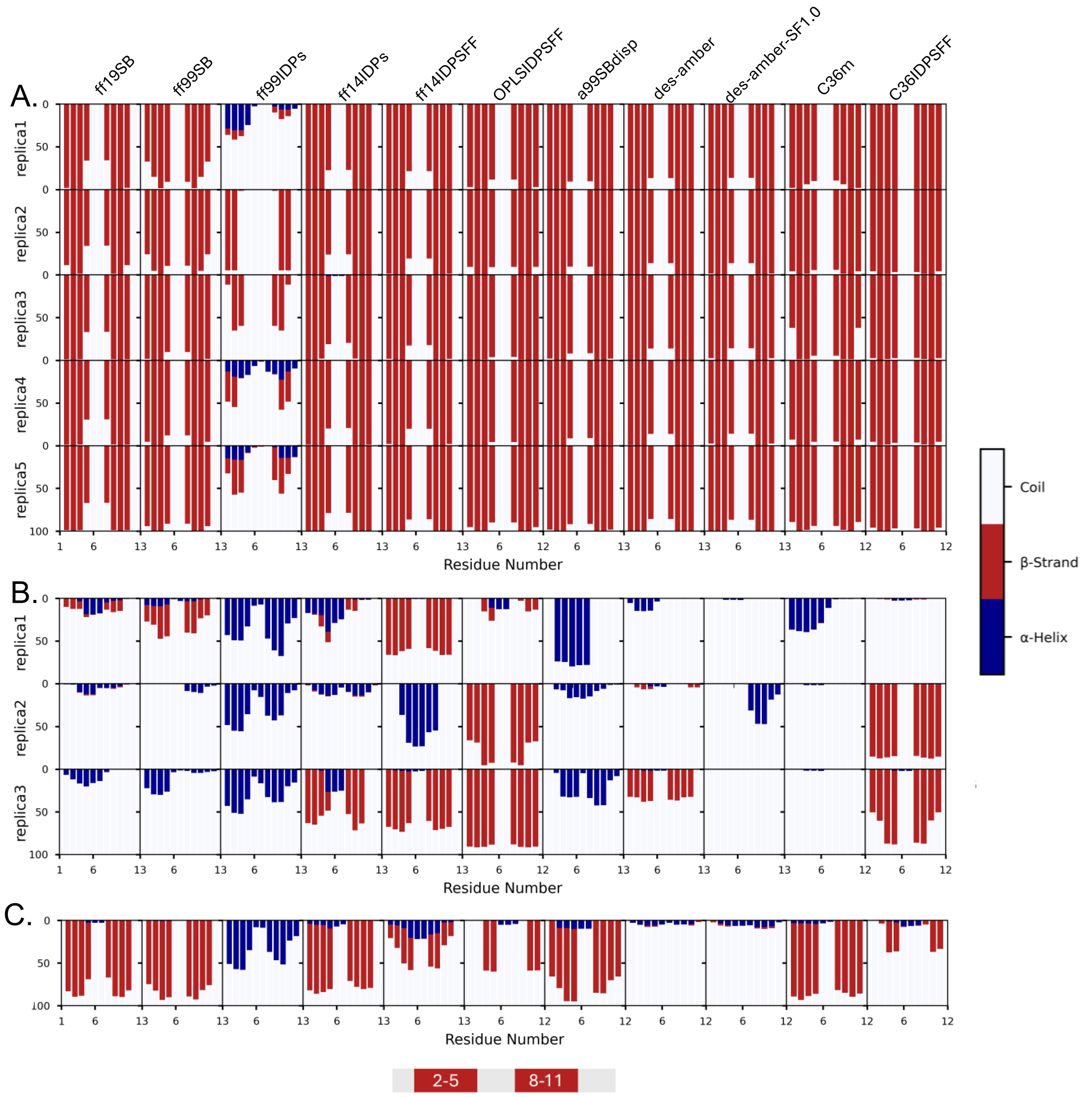


Figure S13: **Per-residue secondary structure preferences for Trp-zip (PDB ID: 1LE1)** across simulations. Bars represent the percentage occurrence of each secondary structure type per residue in (A) five 200-ns replicas, (B) three 1-μs replicas, and (C) a 10-μs simulation. The secondary structure assignment in the experimentally folded structure is shown for reference. Red denotes $\beta$-strand, and blue denotes $\alpha$-helix or 3-10 helix, and the absence of color corresponds to coil or turn regions.

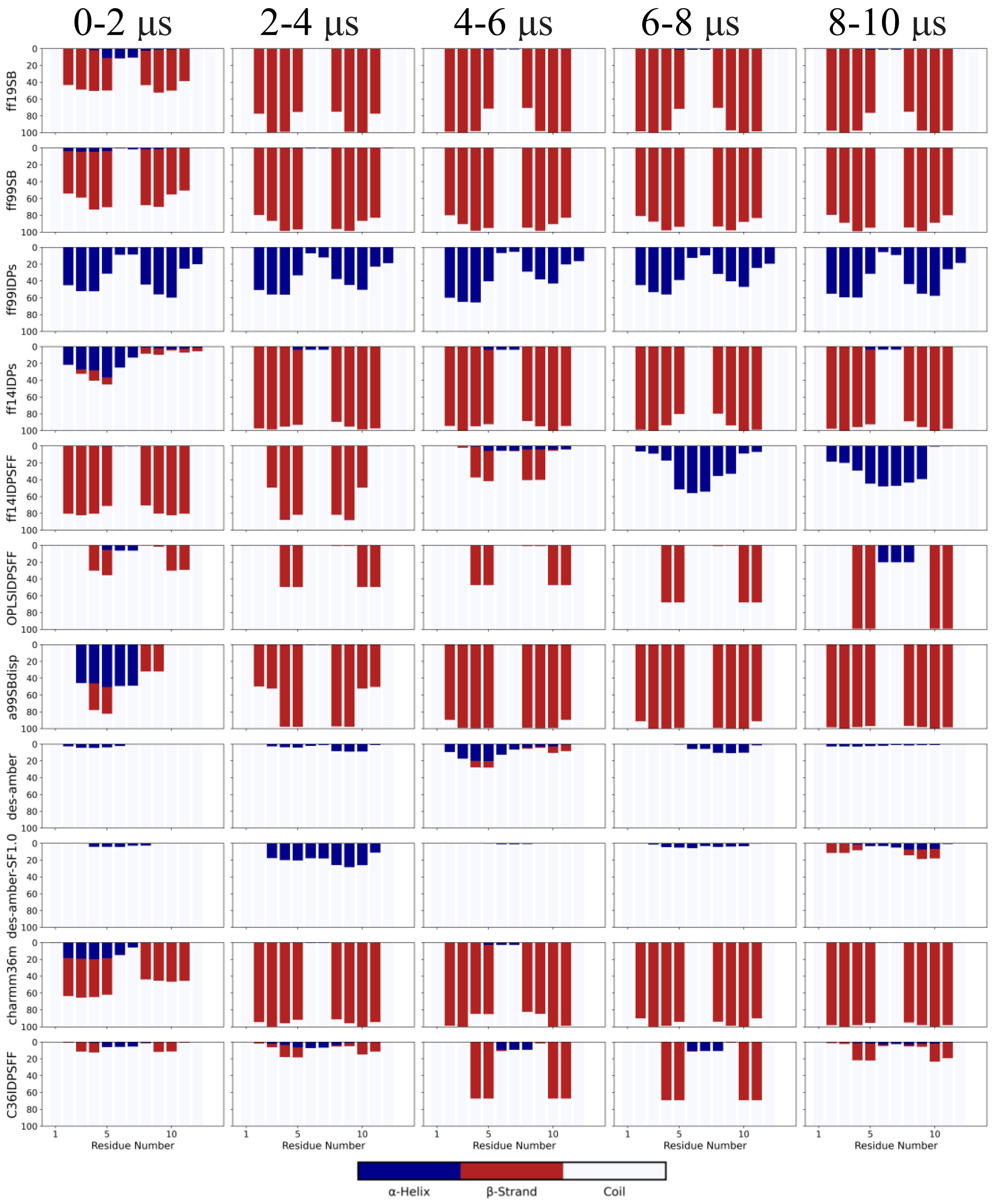


Figure S14: **Per residue secondary structure preferences for Trp-zip (PDB ID: 1LE1)) during the 10 $\mu$s simulation, shown as blocks of 2 $\mu$s intervals.** Bars represent the percentage occurrence of each secondary structure type per residue. Red indicates $\beta$-strand, blue indicates $\alpha$-helix or 3-10 helix, and the absence of color corresponds to coil or turn regions.

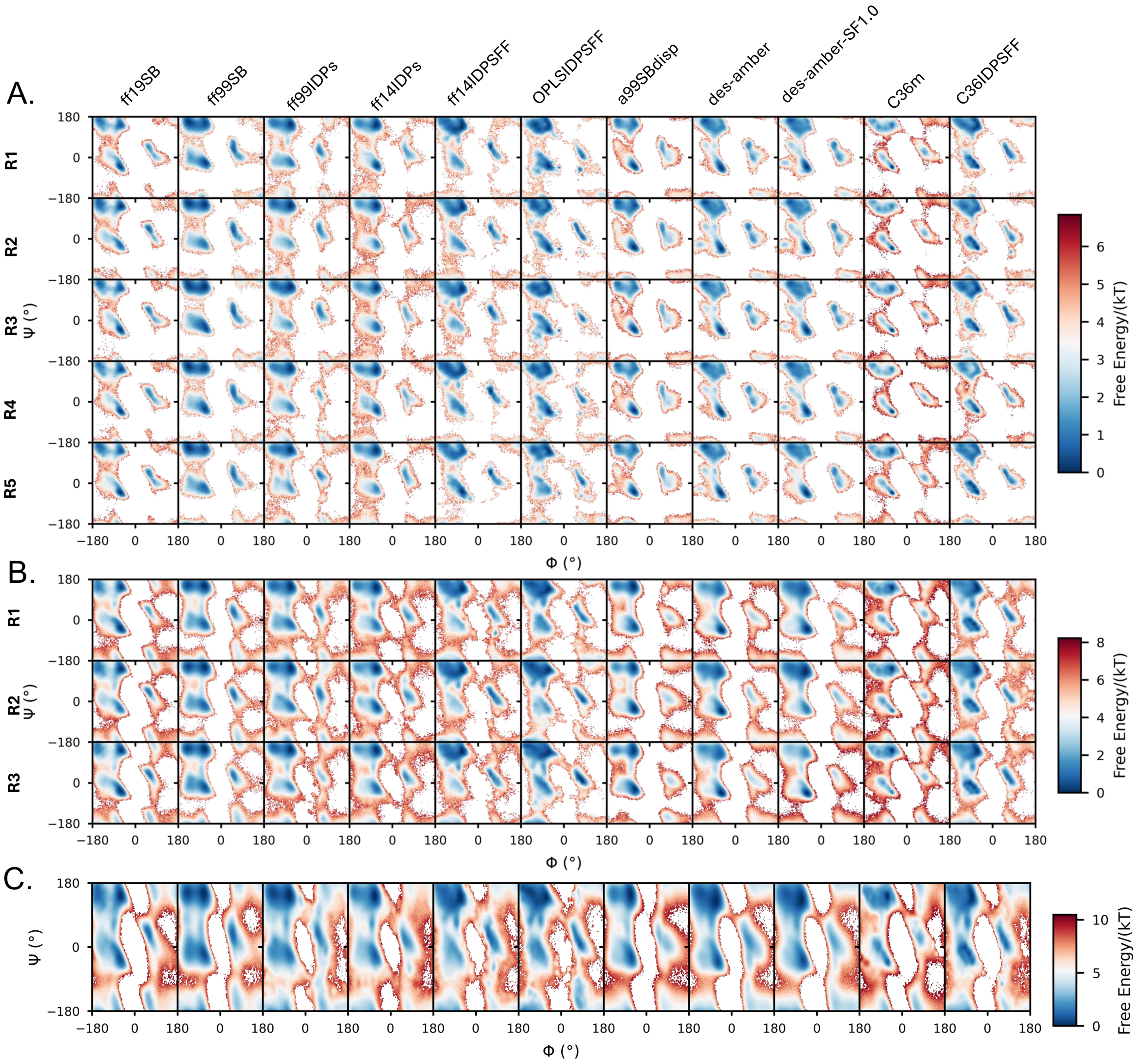


Figure S15: **Ramachandran plots for $\beta$-defensin-1 (PDB ID: 1IJU)**. (A) Five 200-ns simulation replicas initiated from the experimentally folded structure. (B) Three 1-μs simulation replicas initiated from the linear peptide sequence. (C) 10-μs simulation initiated from the linear peptide sequence.

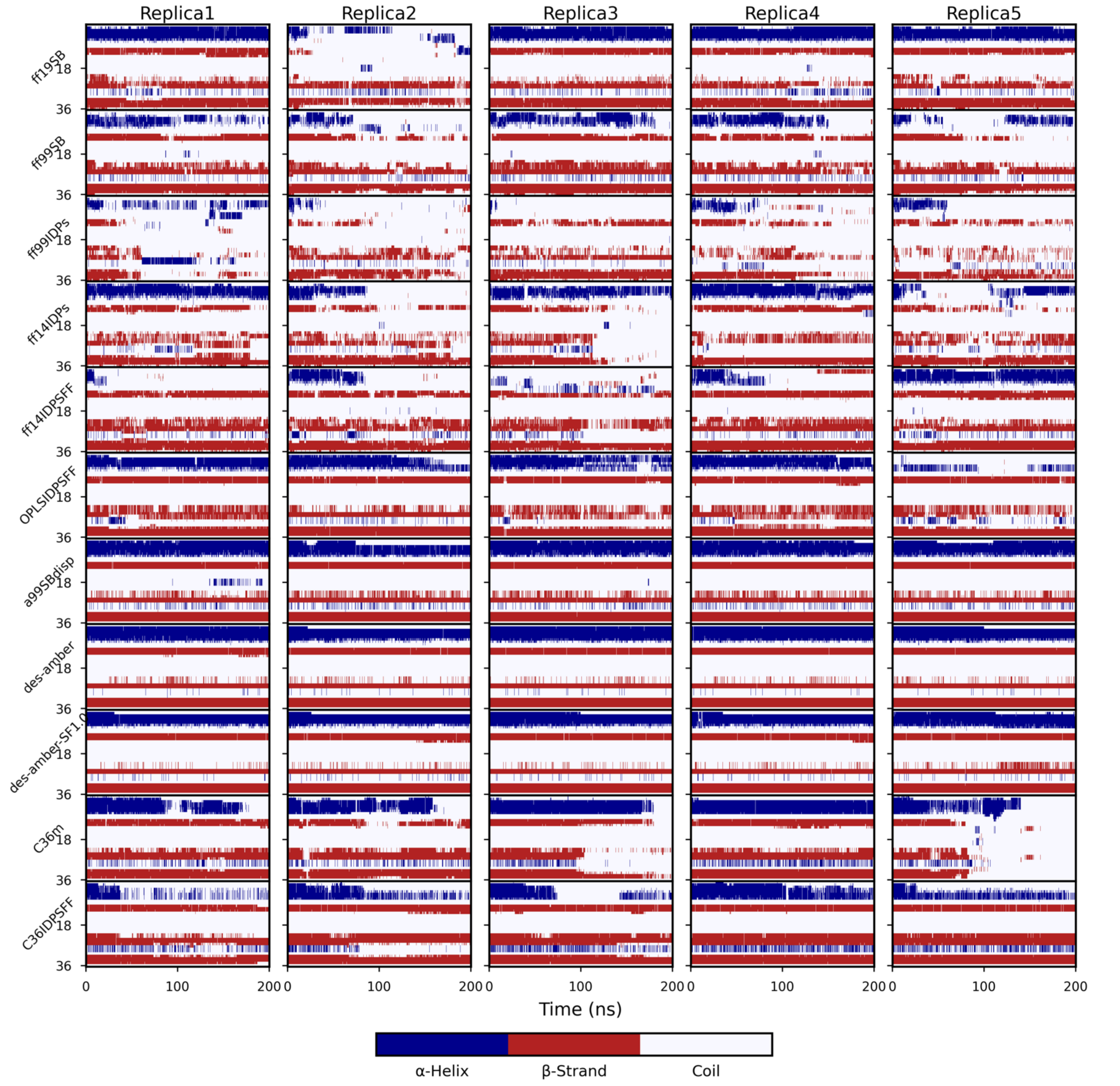


Figure S16: **Per-residue secondary structure time series for $\beta$-defensin-1 (PDB ID: 1IJU)** for five 200-ns simulation replicas initiated from the folded structure. Red denotes $\beta$-strand, and blue denotes $\alpha$-helix or 3-10 helix, and the absence of color corresponds to coil or turn regions.

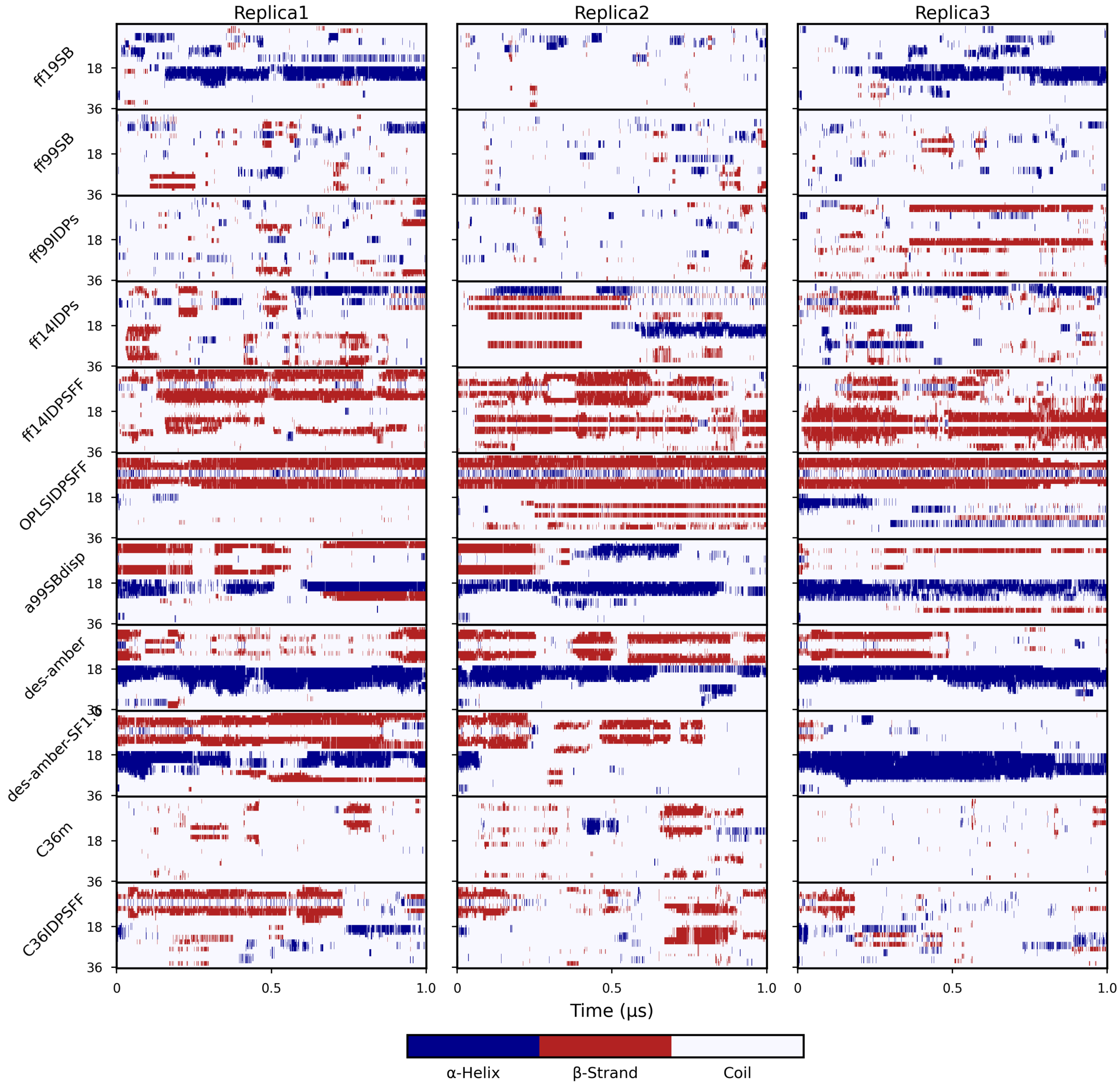


Figure S17: **Per-residue secondary structure time series for $\beta$-defensin-1 (PDB ID: 1IJU)** for three 1 $\mu$s simulation replicas initiated from the folded structure. Red denotes $\beta$-strand, and blue denotes $\alpha$-helix or 3-10 helix, and the absence of color corresponds to coil or turn regions.

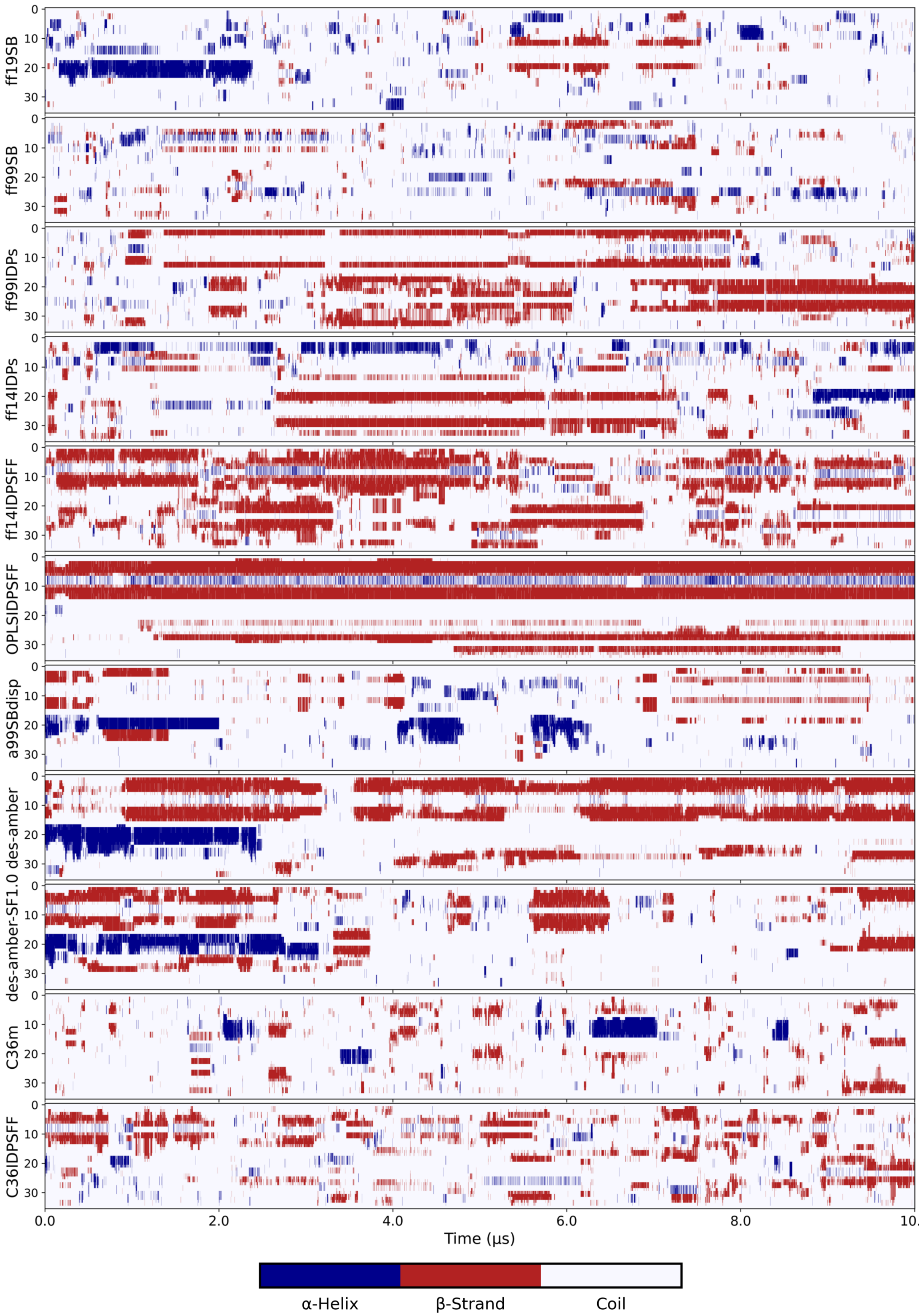


Figure S18: **Per-residue secondary structure time series for $\beta$-defensin-1 (PDB ID: 1IJU)** for the 10 $\mu$s simulation replicas initiated from the folded structure. Red denotes $\beta$-strand, blue denotes $\alpha$-helix or 3–10 helix, and the absence of color corresponds to coil or turn regions.

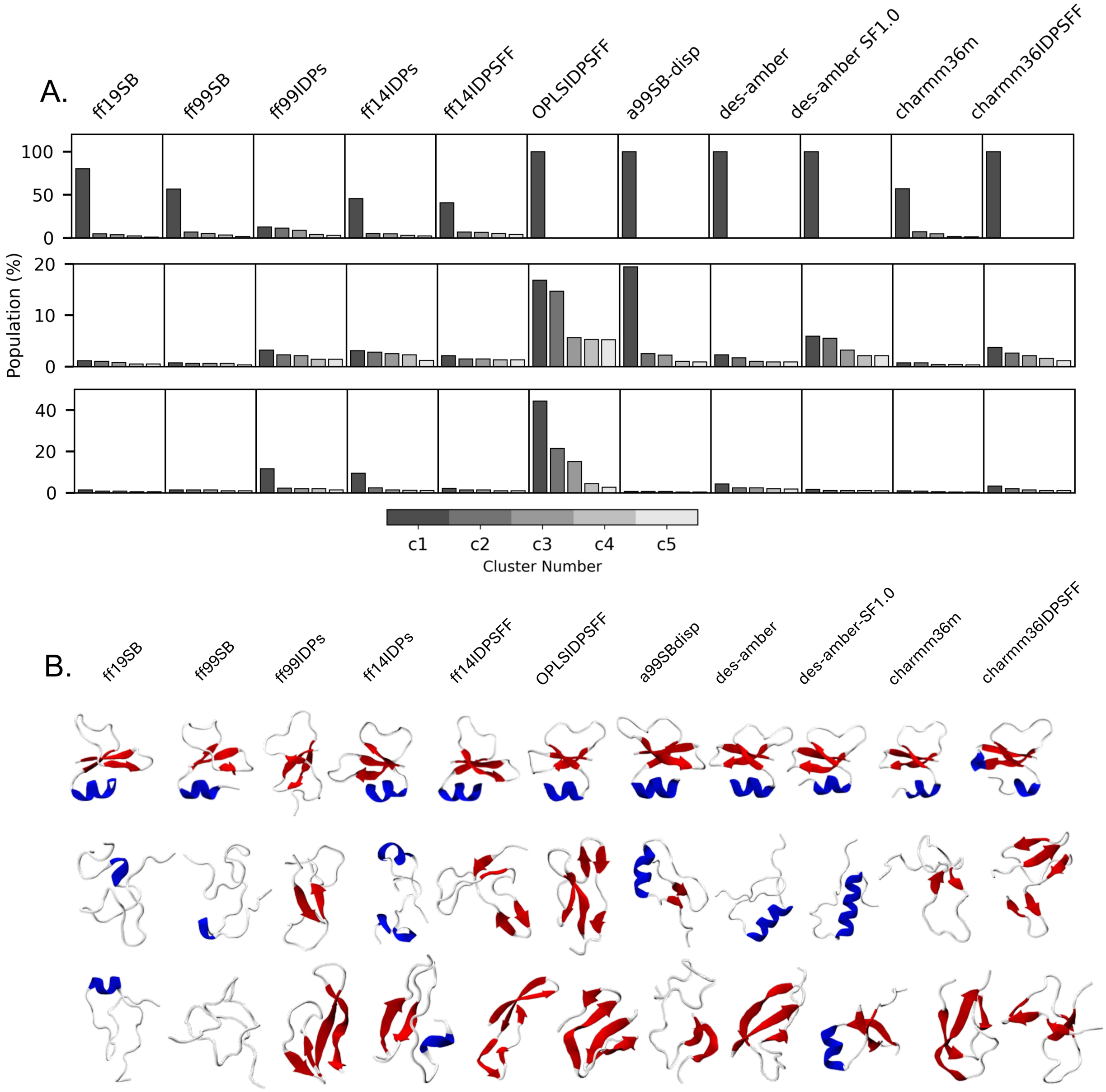


Figure S19: **Clustering results showing (A) cluster populations and (B) representative structures (centroids) of the most populated cluster for $\beta$-defensin-1 (PDB ID: 1IJU).** The top panel shows clustering from five 200-ns stability simulations initiated from the folded structure; the middle panel shows clustering from three 1-µs simulation replicas initiated from the linear peptide sequence; and the bottom panel shows clustering from a 10-µs folding simulation initiated from the extended peptide conformation. Red indicates $\beta$-strand, blue indicates $\alpha$-helix or 3-10 helix, and white indicates coil or turn regions.

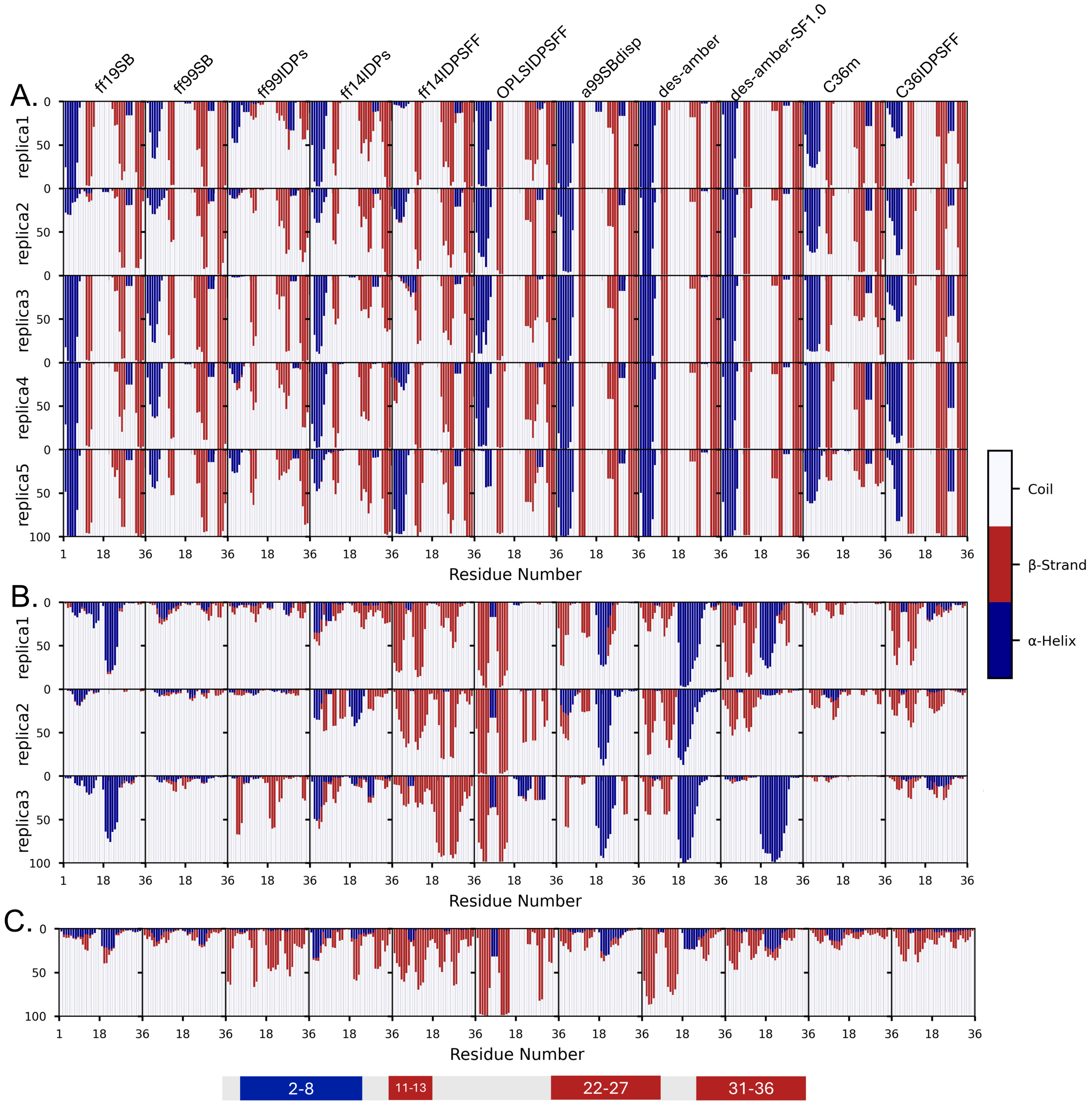


Figure S20: **Per-residue secondary structure preferences for $\beta$-defensin-1 (PDB ID: 1IJU)** across simulations. Bars represent the percentage occurrence of each secondary structure type per residue in (A) five 200-ns replicas, (B) three 1-µs replicas, and (C) a 10-µs simulation. The secondary structure assignment in the experimentally folded structure is shown for reference. Red denotes $\beta$-strand, and blue denotes $\alpha$-helix or 3-10 helix, and the absence of color corresponds to coil or turn regions.

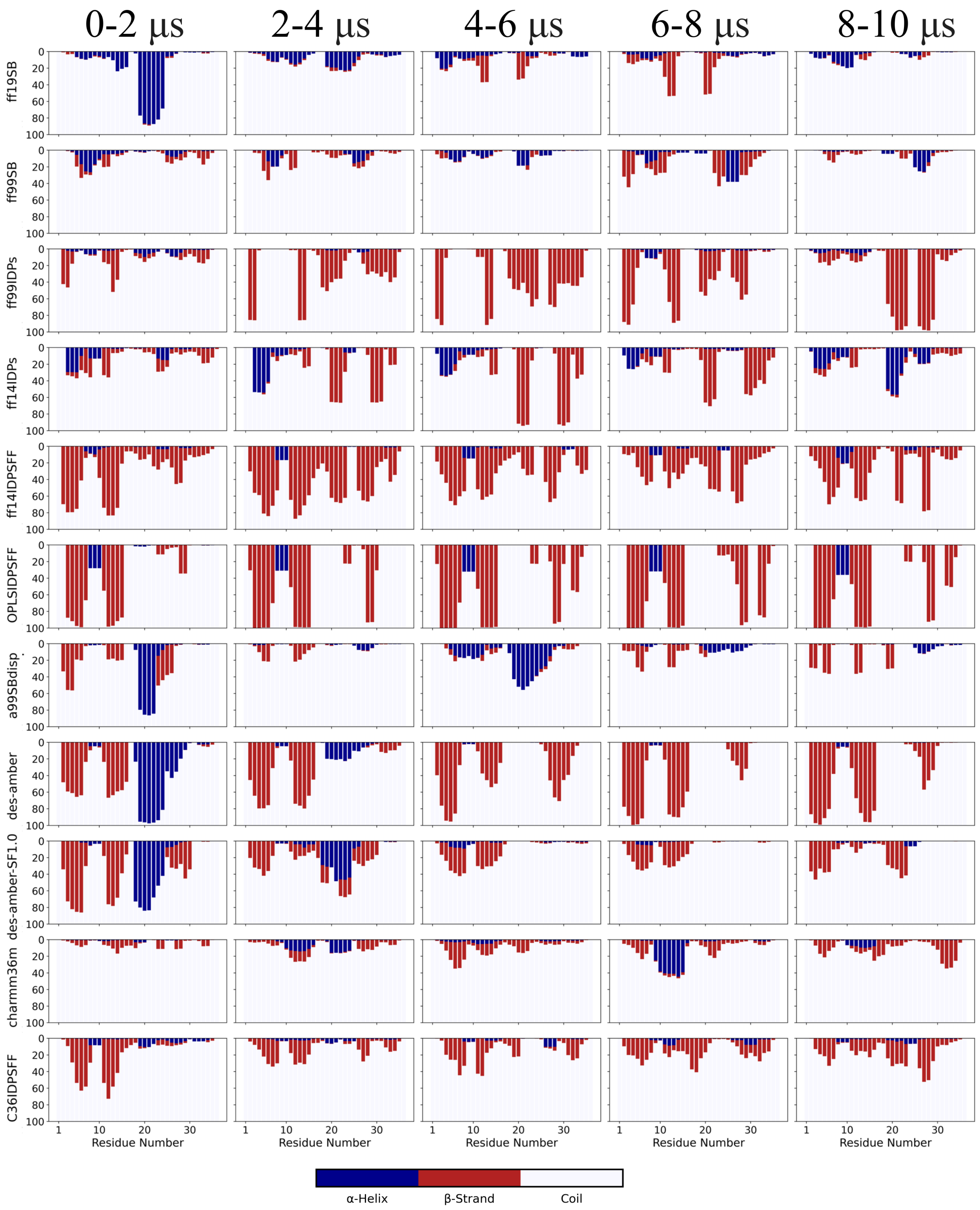


Figure S21: **Per residue secondary structure preferences for $\beta$-defensin-1 (PDB ID 1IJU) during the 10 $\mu$s simulation, shown as blocks of 2 $\mu$s intervals.** Bars represent the percentage occurrence of each secondary structure type per residue. Red indicates $\beta$-strand, blue indicates $\alpha$-helix or 3-10 helix, and the absence of color corresponds to coil or turn regions.

### Partially Structured peptides

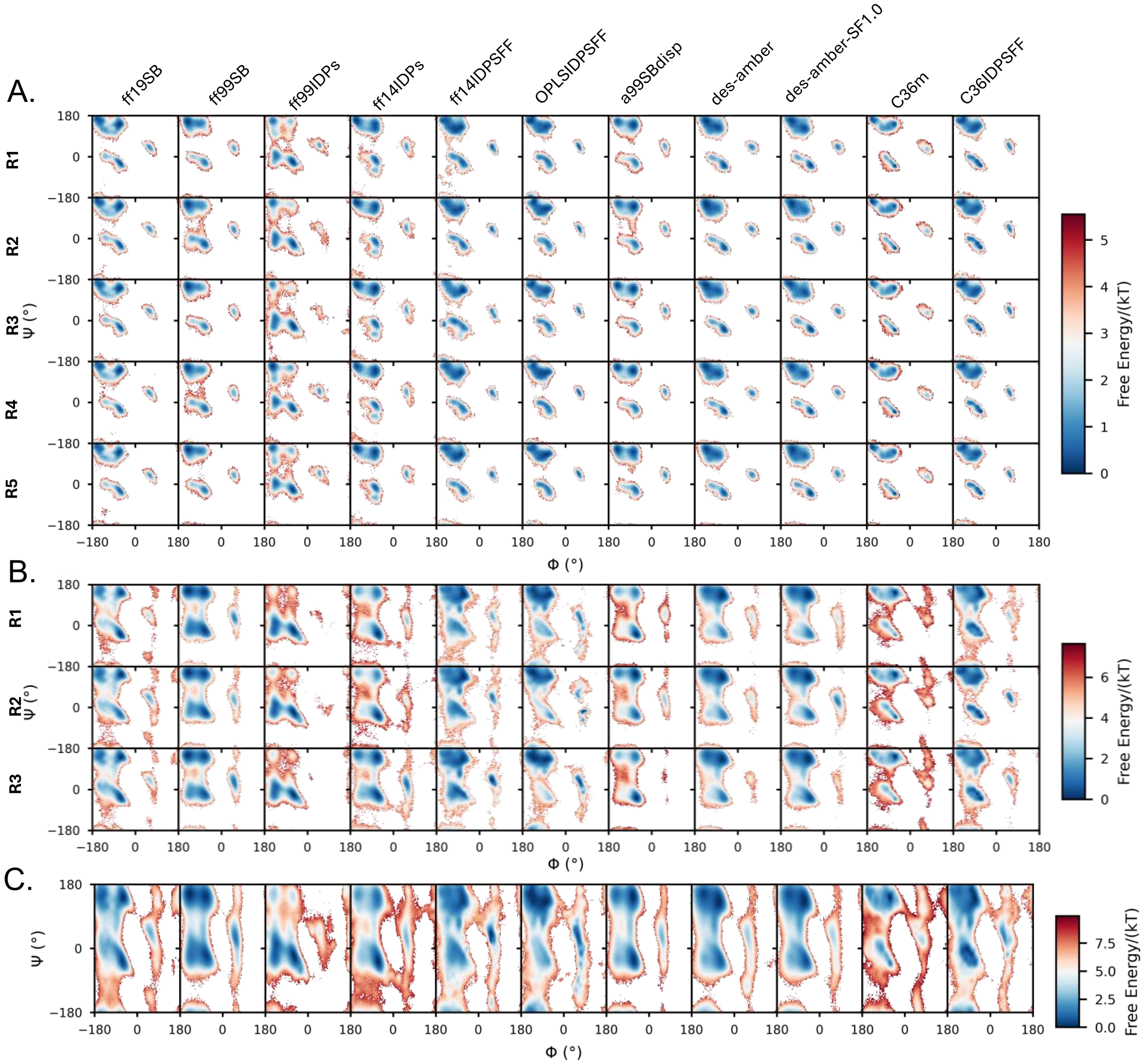


Figure S22: **Ramachandran plots for C-termini from Protein G (PDB ID: 1GB1)**. (A) Five 200-ns simulation replicas initiated from the experimentally folded structure. (B) Three 1-µs simulation replicas initiated from the linear peptide sequence. (C) 10-µs simulation initiated from the linear peptide sequence.

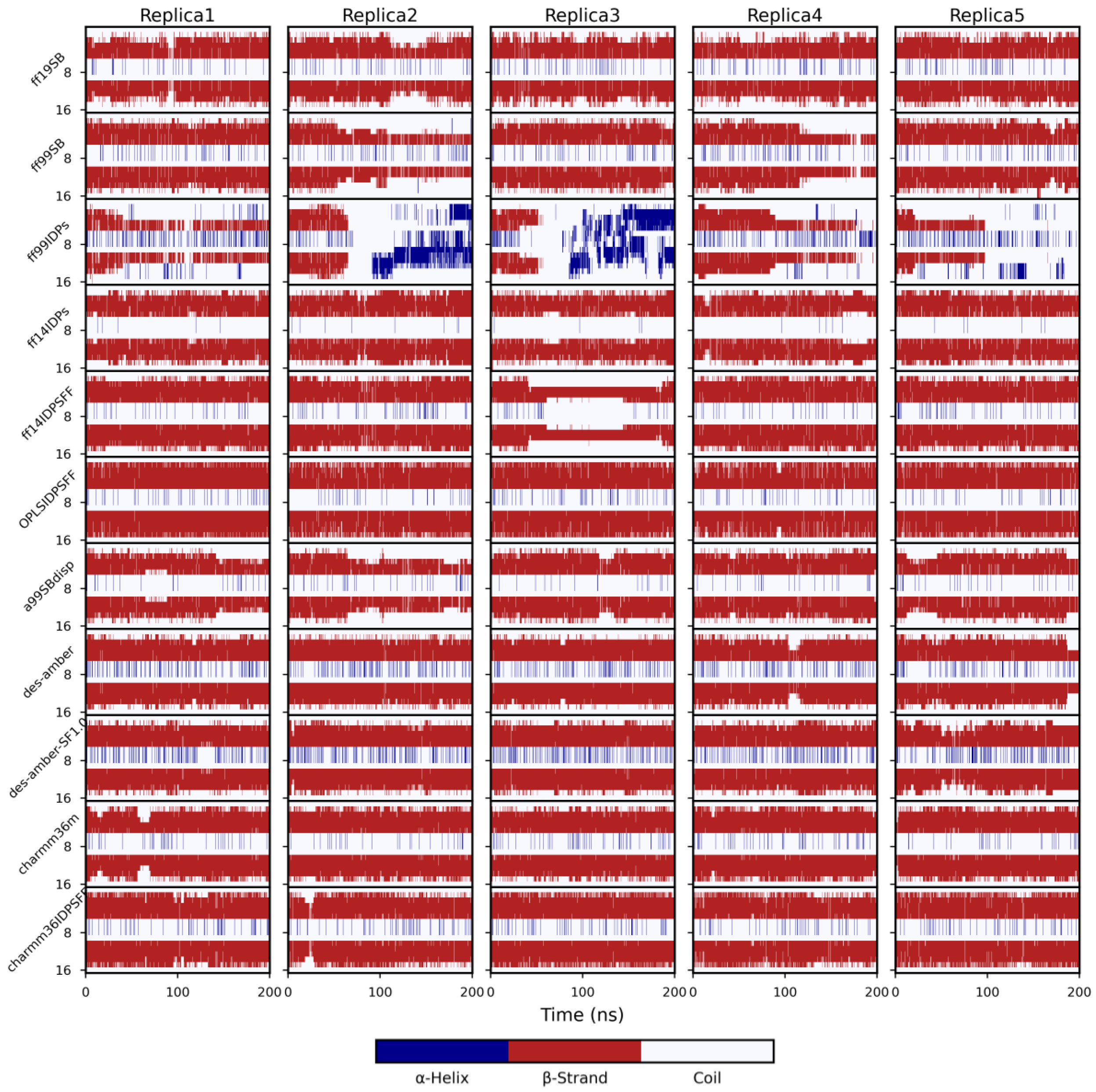


Figure S23: **Per-residue secondary structure time series for C-termini from Protein G (PDB ID: 1GB1)** for five 200-ns simulation replicas initiated from the folded structure. Red denotes $\beta$-strand, and blue denotes $\alpha$-helix or 3-10 helix, and the absence of color corresponds to coil or turn regions.

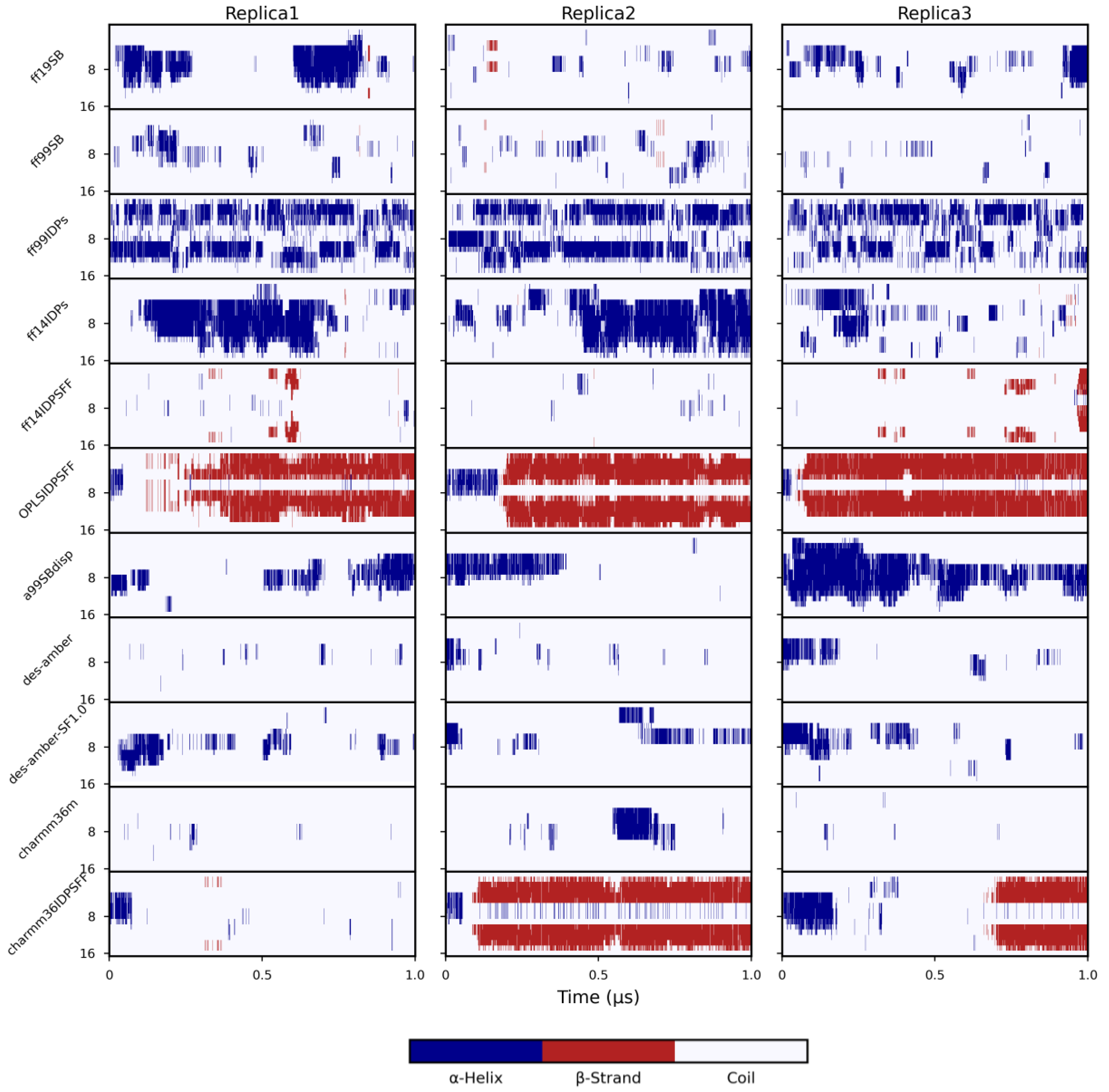


Figure S24: **Per-residue secondary structure time series for C-termini from Protein G (PDB ID: 1GB1)** for three 1 $\mu$s simulation replicas initiated from the folded structure. Red denotes $\beta$-strand, and blue denotes $\alpha$-helix or 3-10 helix, and the absence of color corresponds to coil or turn regions.

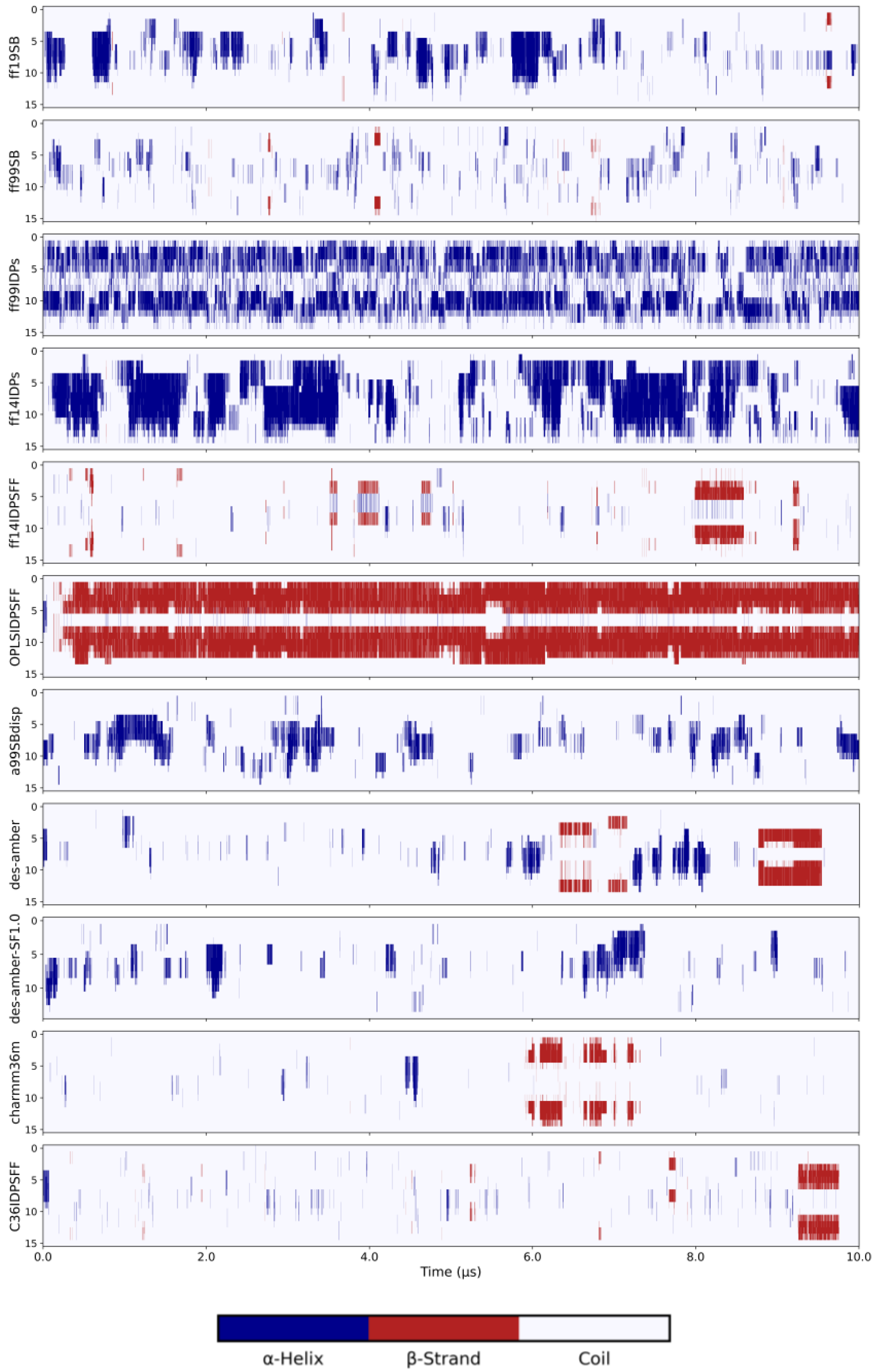


Figure S25: **Per-residue secondary structure time series for C-termini from Protein G (PDB ID: 1GB1)** for the 10 $\mu$s simulation replicas initiated from the folded structure. Red denotes $\beta$-strand, and blue denotes $\alpha$-helix or 3-10 helix, and the absence of color corresponds to coil or turn regions.

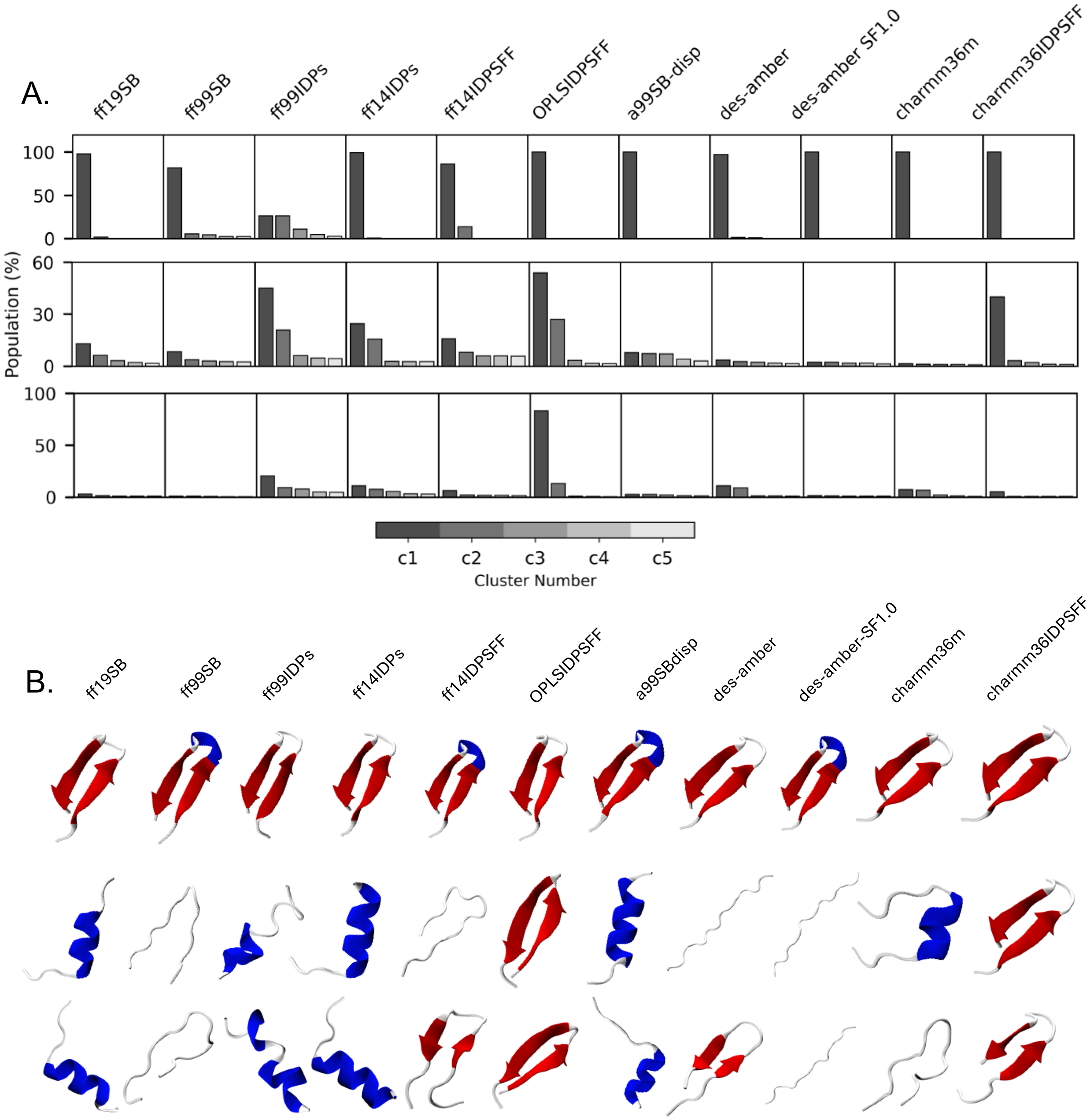


Figure S26: **Clustering results showing (A) cluster populations and (B) representative structures (centroids) of the most populated cluster for C-termini from Protein G (PDB ID: 1GB1).** The top panel shows clustering from five 200-ns stability simulations initiated from the folded structure; the middle panel shows clustering from three 1-μs simulation replicas initiated from the linear peptide sequence; and the bottom panel shows clustering from a 10-μs folding simulation initiated from the extended peptide conformation. Red indicates $\beta$-strand, blue indicates $\alpha$-helix or 3-10 helix, and white indicates coil or turn regions.

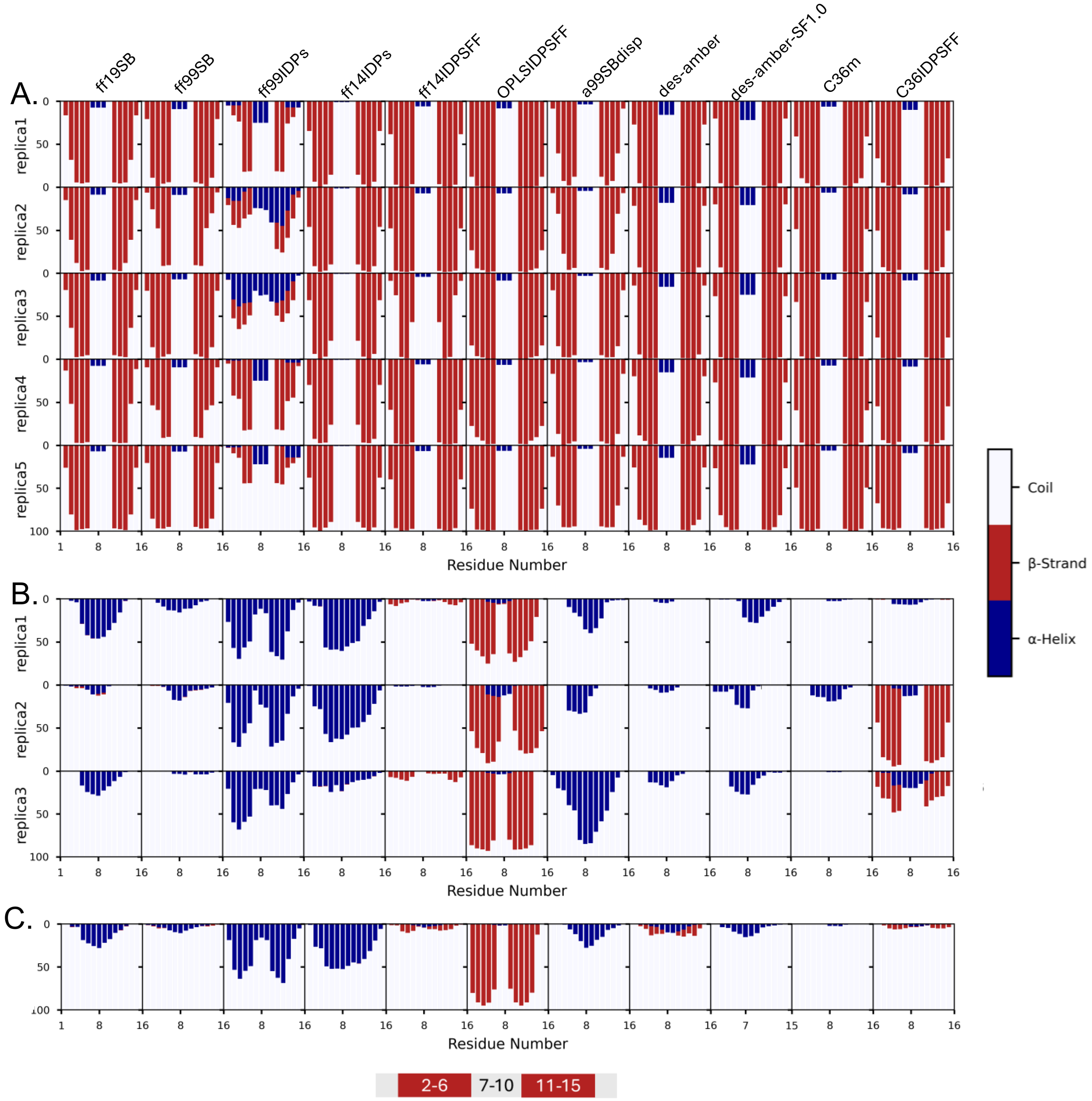


Figure S27: **Per-residue secondary structure preferences for C-termini from Protein G (PDB ID: 1GB1)** across simulations. Bars represent the percentage occurrence of each secondary structure type per residue in (A) five 200-ns replicas, (B) three 1-μs replicas, and (C) a 10-μs simulation. The secondary structure assignment in the experimentally folded structure is shown for reference. Red denotes $\beta$-strand, and blue denotes $\alpha$-helix or 3-10 helix, and the absence of color corresponds to coil or turn regions.

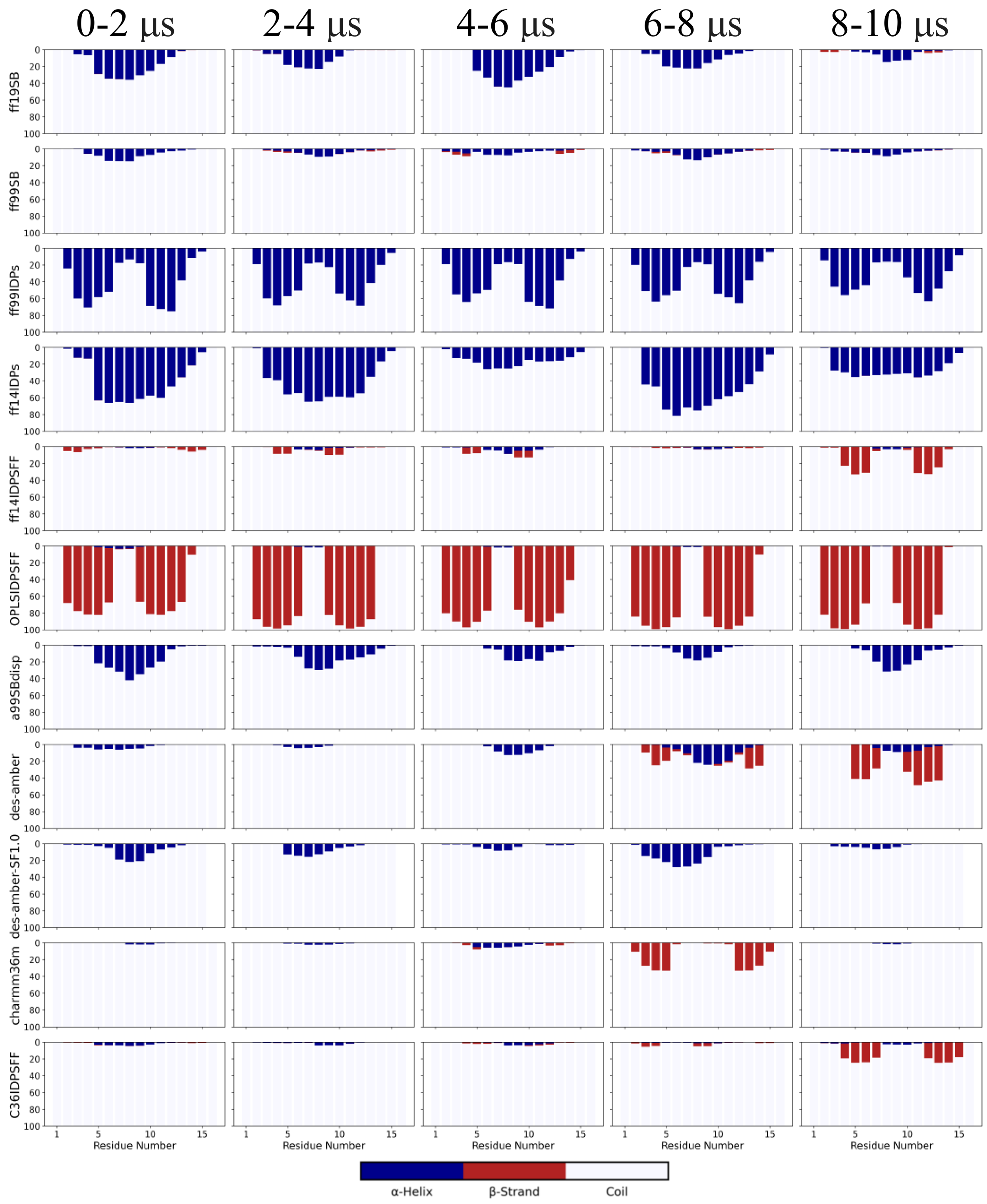


Figure S28: **Per residue secondary structure preferences for C-termini from Protein G (PDB ID: 1GB1) during the 10 $\mu$s simulation, shown as blocks of 2 $\mu$s intervals.** Bars represent the percentage occurrence of each secondary structure type per residue. Red indicates $\beta$-strand, blue indicates $\alpha$-helix or 3-10 helix, and the absence of color corresponds to coil or turn regions.

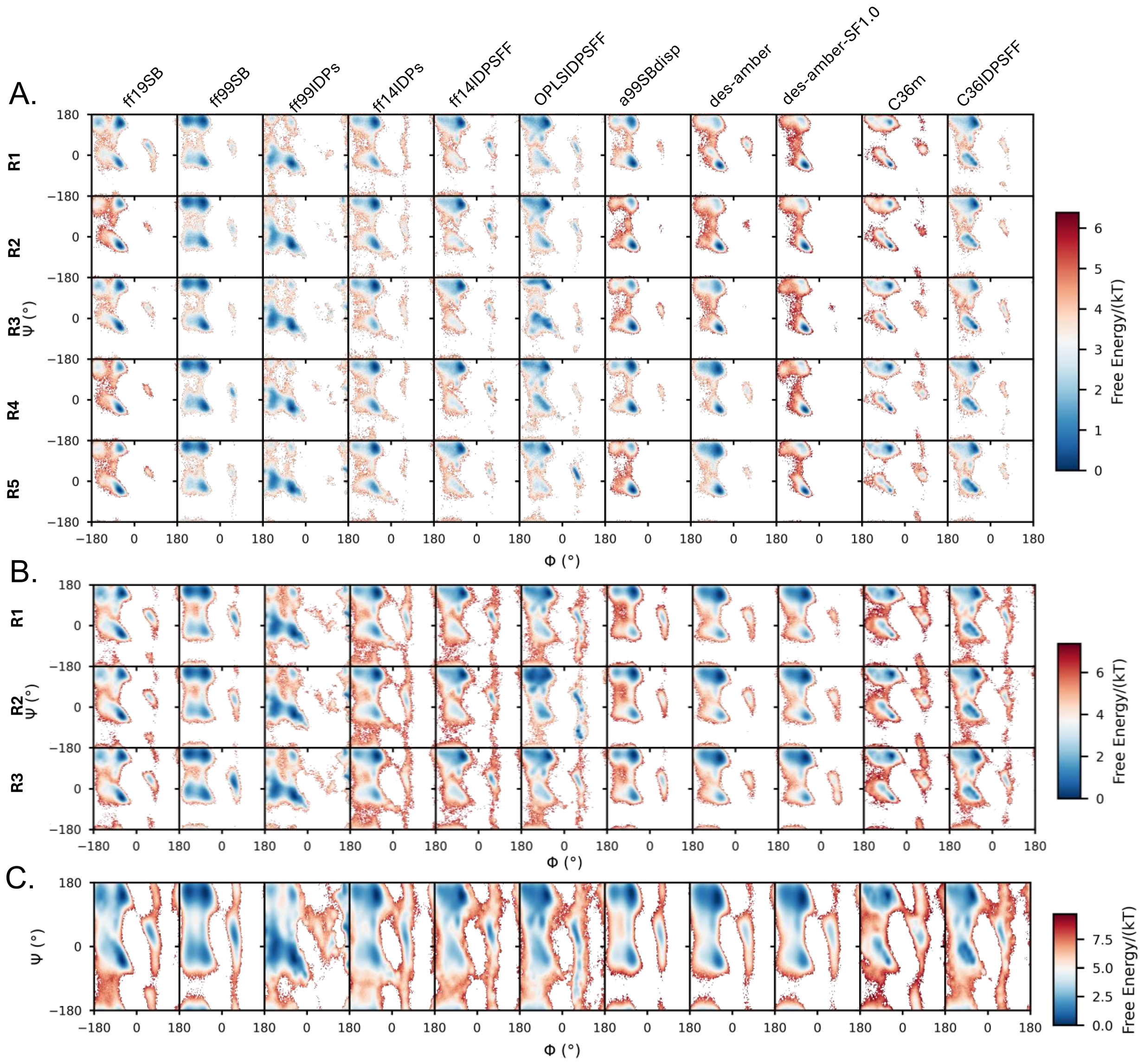


Figure S29: **Ramachandran plots for C-peptide (PDB ID: 5RSA)**. (A) Five 200-ns simulation replicas initiated from the experimentally folded structure. (B) Three 1-µs simulation replicas initiated from the linear peptide sequence. (C) 10-µs simulation initiated from the linear peptide sequence.

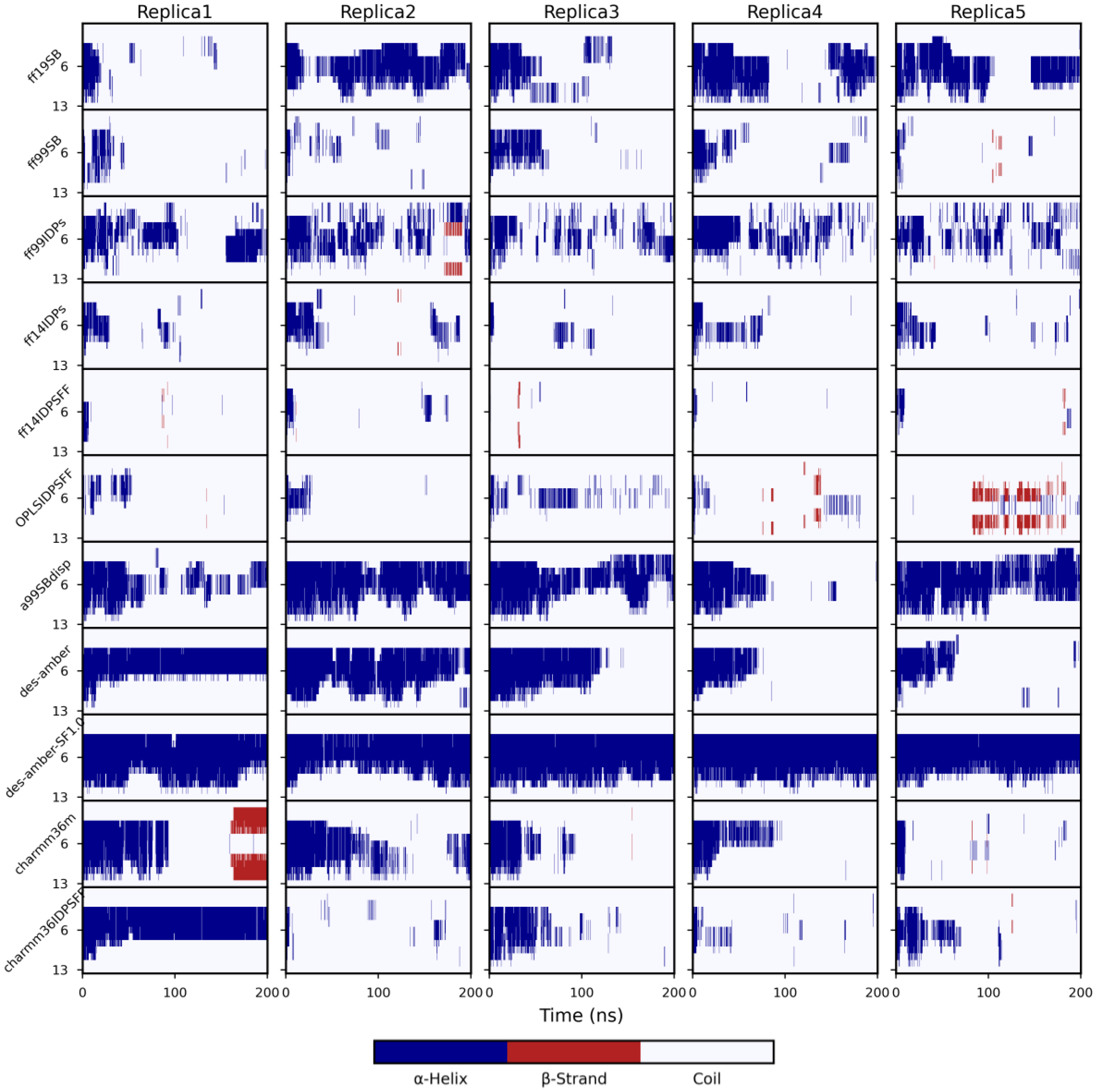


Figure S30: **Per-residue secondary structure time series for C-peptide (PDB ID: 5RSA)** for five 200-ns simulation replicas initiated from the folded structure. Red denotes $\beta$-strand, and blue denotes $\alpha$-helix or 3-10 helix, and the absence of color corresponds to coil or turn regions.

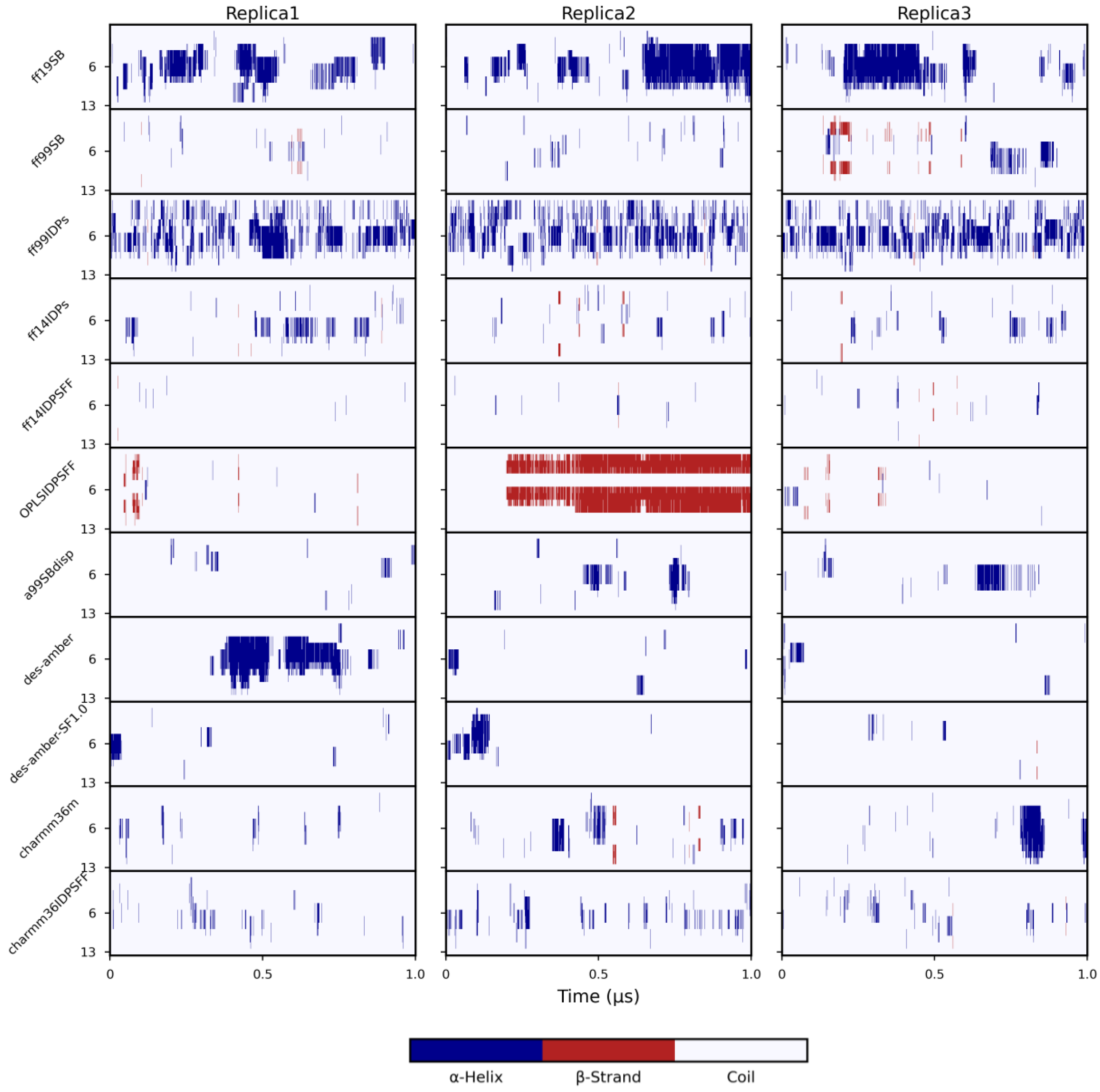


Figure S31: **Per-residue secondary structure time series for C-peptide (PDB ID: 5RSA)** for three 1 $\mu$s simulation replicas initiated from the folded structure. Red denotes $\beta$-strand, and blue denotes $\alpha$-helix or 3-10 helix, and the absence of color corresponds to coil or turn regions.

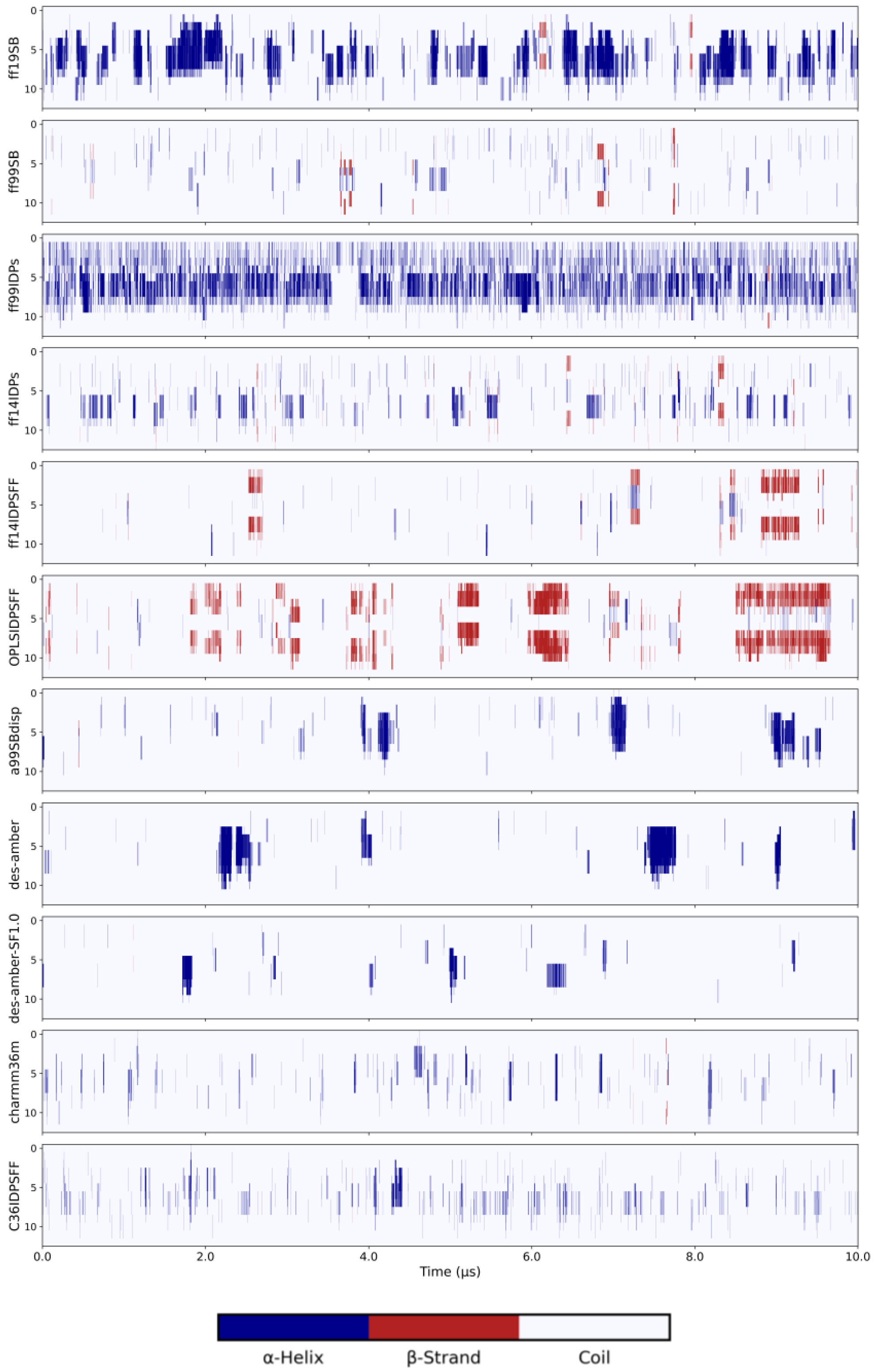


Figure S32: **Per-residue secondary structure time series for C-peptide (PDB ID: 5RSA)** for the 10 $\mu$s simulation replicas initiated from the folded structure. Red denotes $\beta$-strand, and blue denotes $\alpha$-helix or 3-10 helix, and the absence of color corresponds to coil or turn regions.

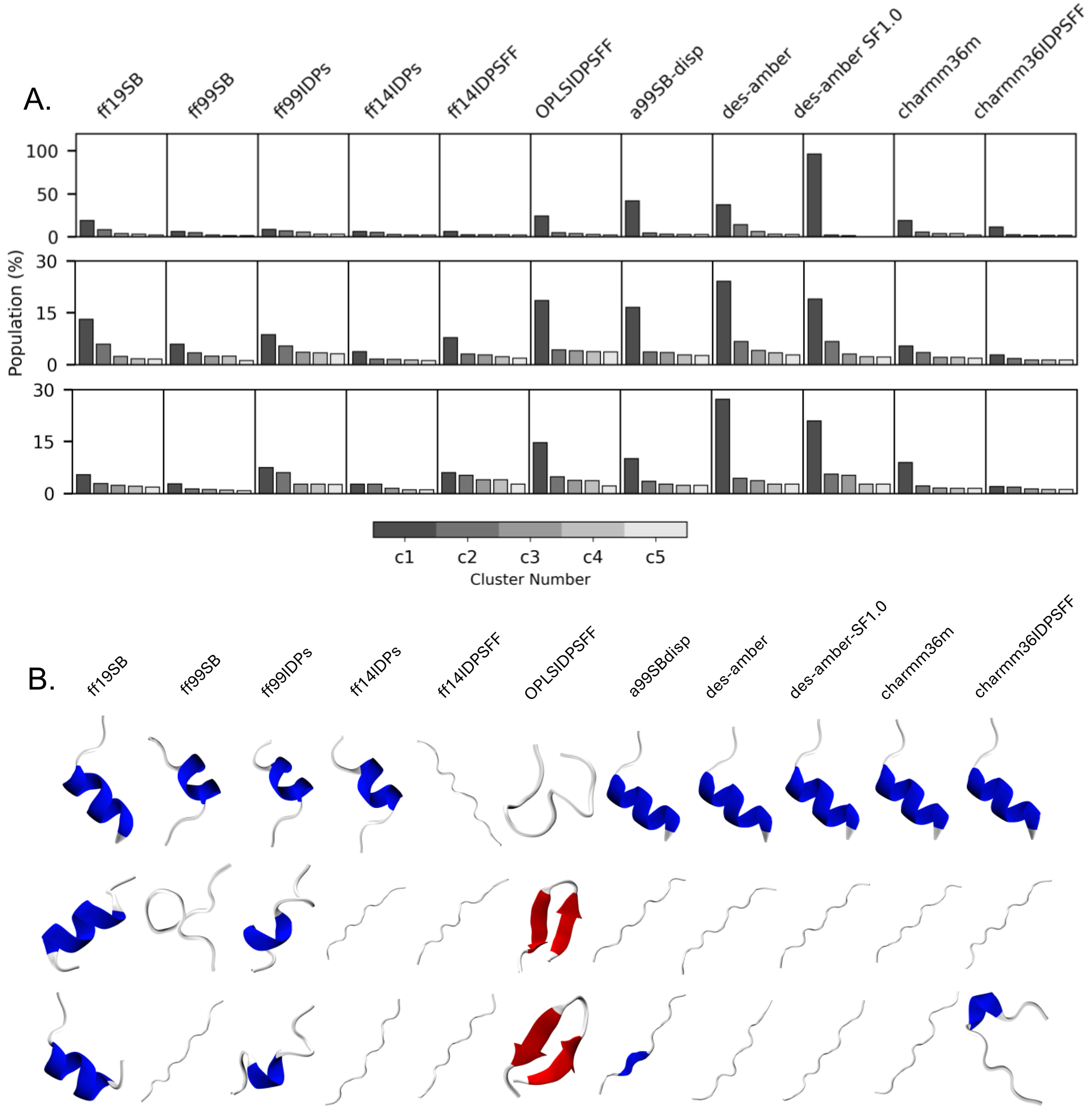


Figure S33: **Clustering results showing (A) cluster populations and (B) representative structures (centroids) of the most populated cluster for C-peptide (PDB ID: 5RSA).** The top panel shows clustering from five 200-ns stability simulations initiated from the folded structure; the middle panel shows clustering from three 1-µs simulation replicas initiated from the linear peptide sequence; and the bottom panel shows clustering from a 10-µs folding simulation initiated from the extended peptide conformation. Red indicates $\beta$-strand, blue indicates $\alpha$-helix or 3-10 helix, and white indicates coil or turn regions.

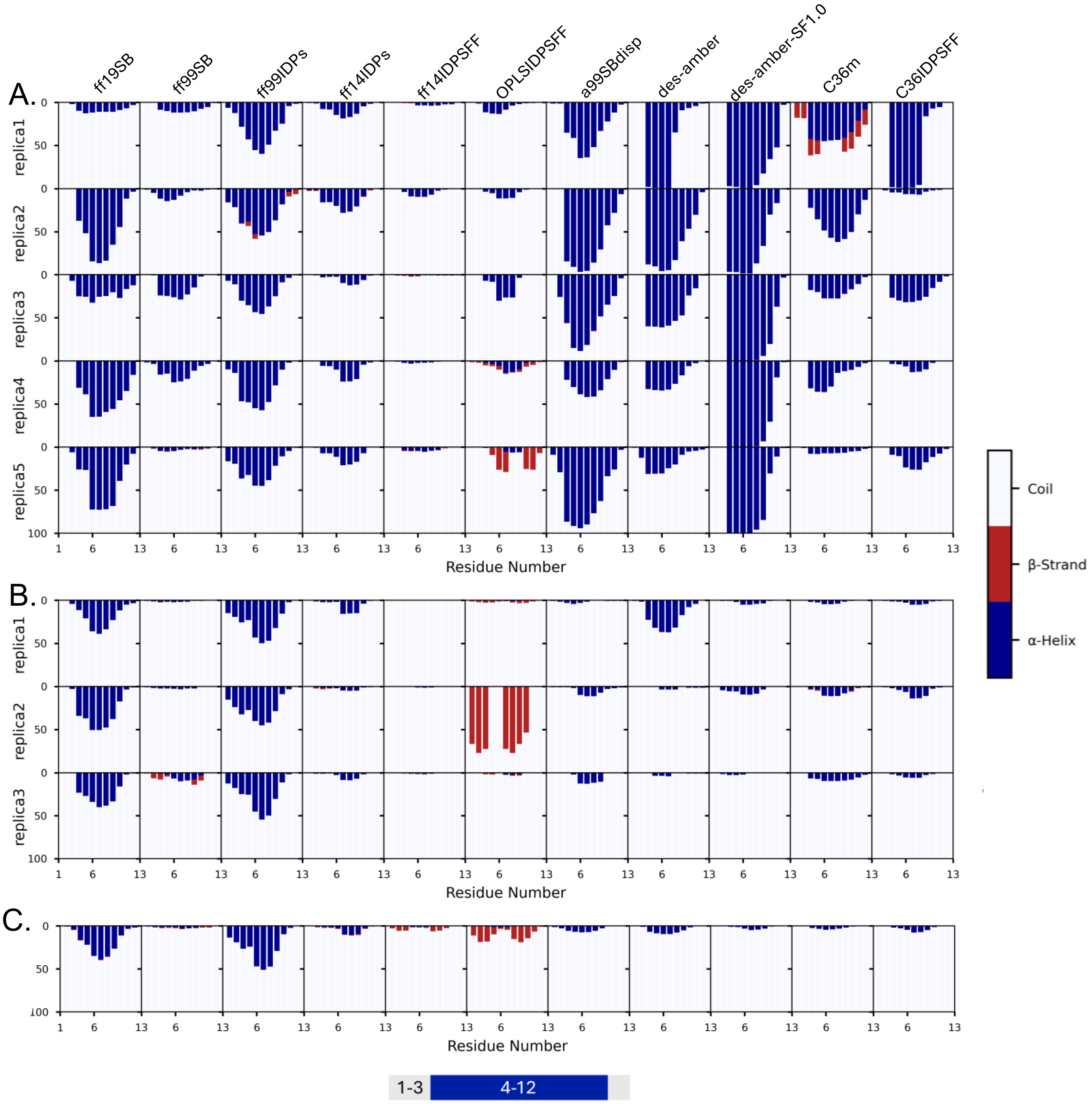


Figure S34: **Per-residue secondary structure preferences for C-peptide (PDB ID: 5RSA)** across simulations. Bars represent the percentage occurrence of each secondary structure type per residue in (A) five 200-ns replicas, (B) three 1-µs replicas, and (C) a 10-µs simulation. The secondary structure assignment in the experimentally folded structure is shown for reference. Red denotes $\beta$-strand, and blue denotes $\alpha$-helix or 3-10 helix, and the absence of color corresponds to coil or turn regions.

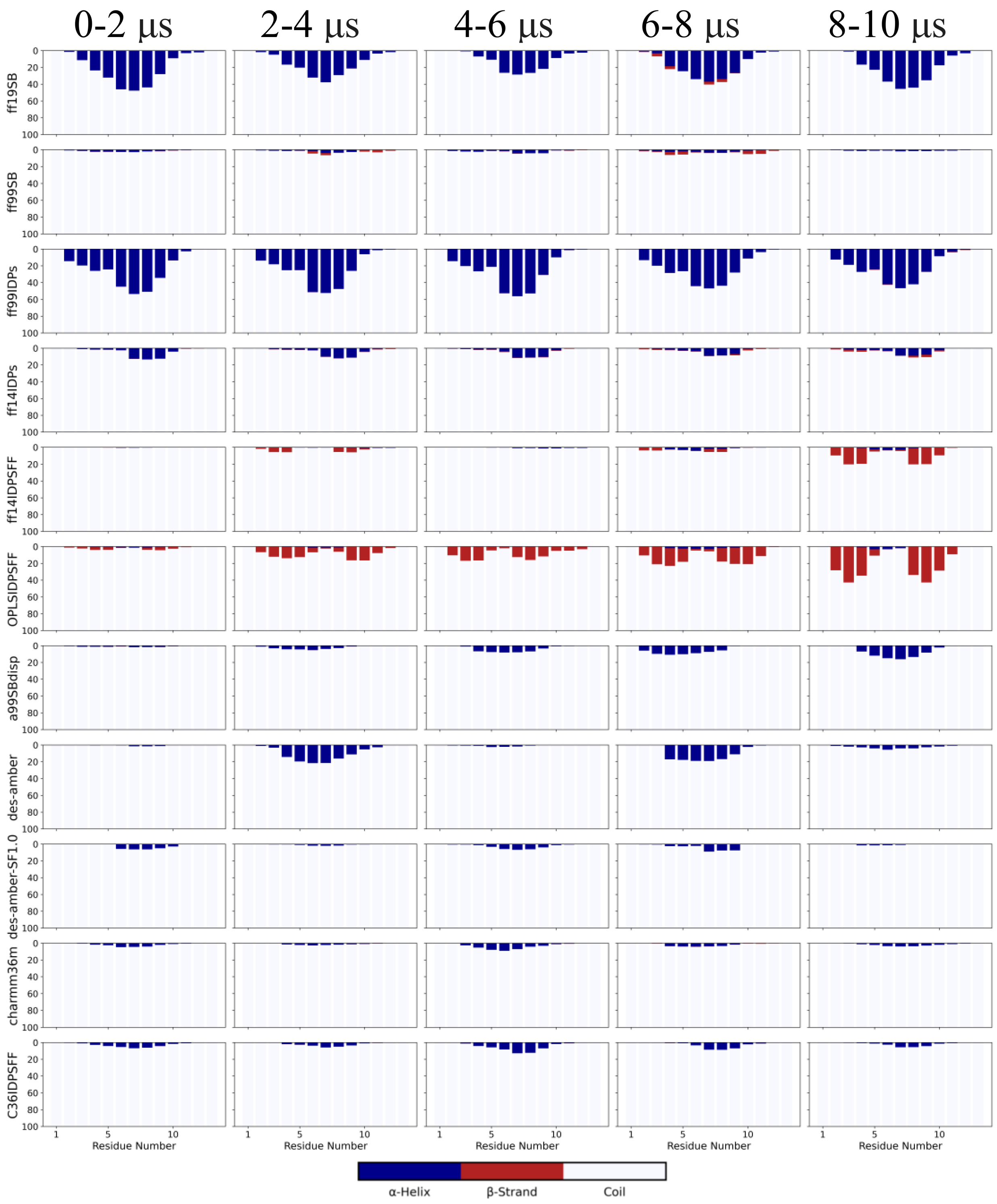


Figure S35: **Per residue secondary structure preferences for C-peptide (PDB ID: 5RSA) during the 10 $\mu$s simulation, shown as blocks of 2 $\mu$s intervals.** Bars represent the percentage occurrence of each secondary structure type per residue. Red indicates $\beta$-strand, blue indicates $\alpha$-helix or 3-10 helix, and the absence of color corresponds to coil or turn regions.

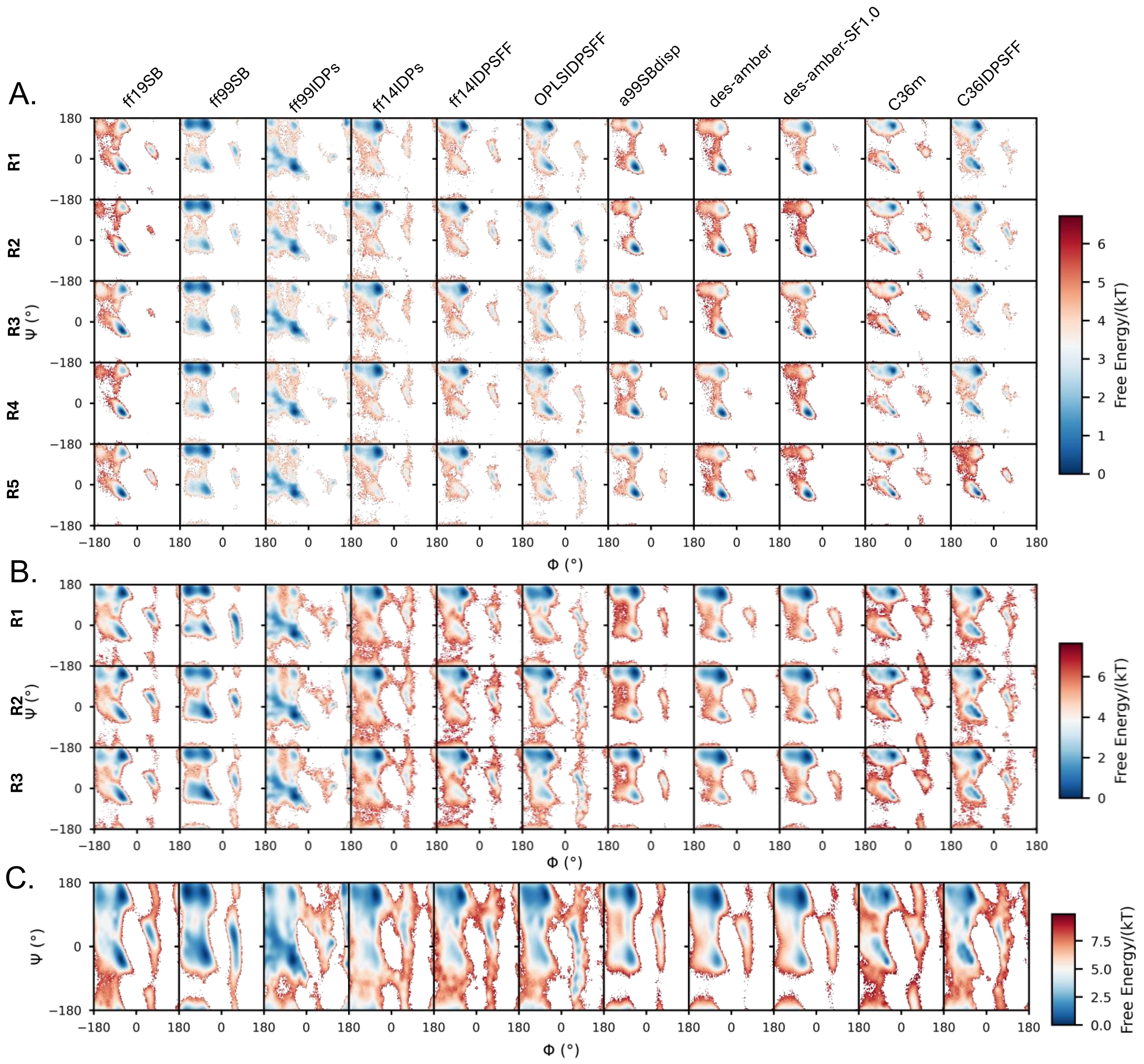


Figure S36: **Ramachandran plots for The EK peptide**. (A) Five 200-ns simulation replicas initiated from the experimentally folded structure. (B) Three 1-μs simulation replicas initiated from the linear peptide sequence. (C) 10-μs simulation initiated from the linear peptide sequence.

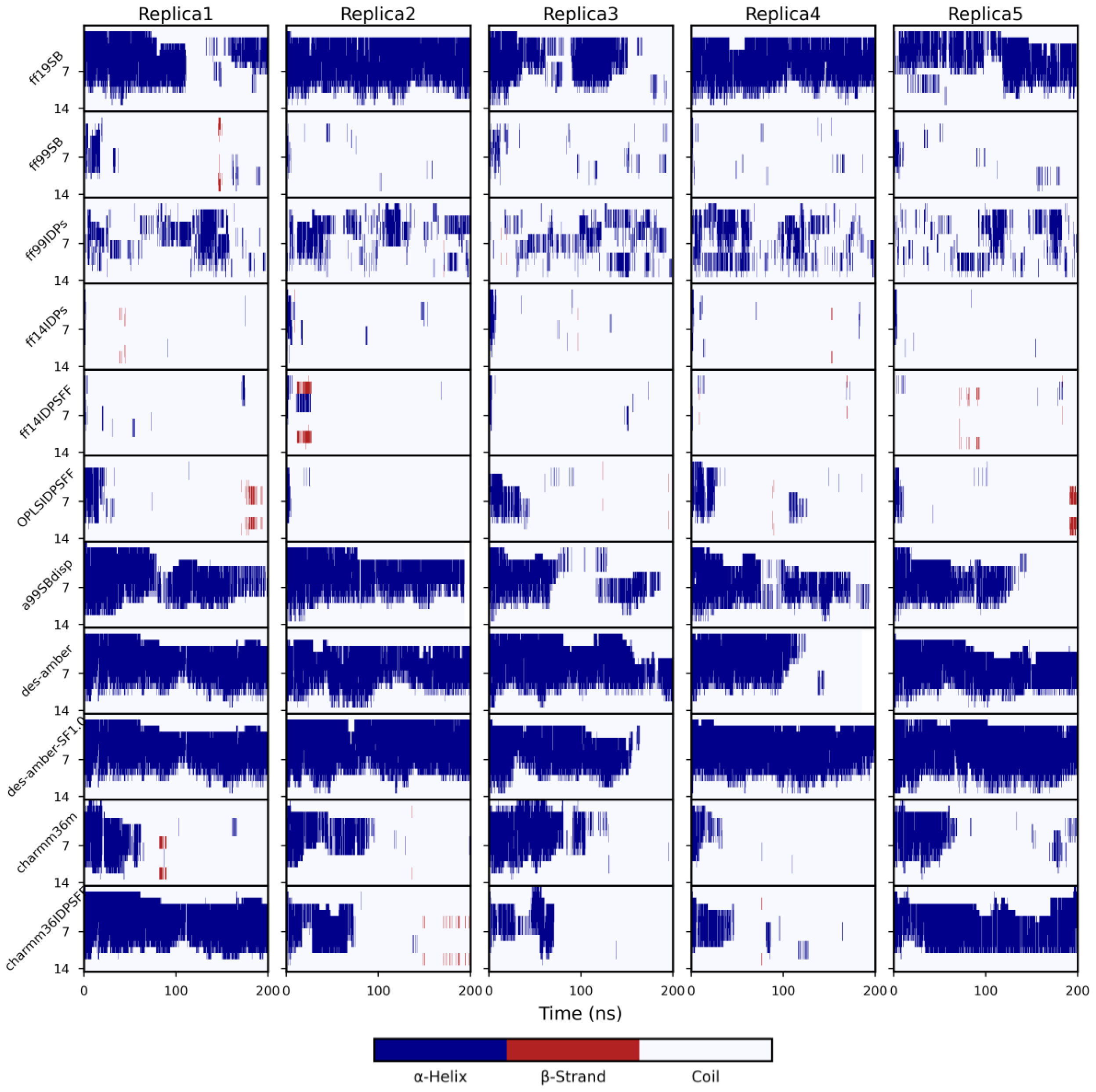


Figure S37: **Per-residue secondary structure time series for The EK peptide** for five 200-ns simulation replicas initiated from the folded structure. Red denotes $\beta$-strand, and blue denotes $\alpha$-helix or 3-10 helix, and the absence of color corresponds to coil or turn regions.

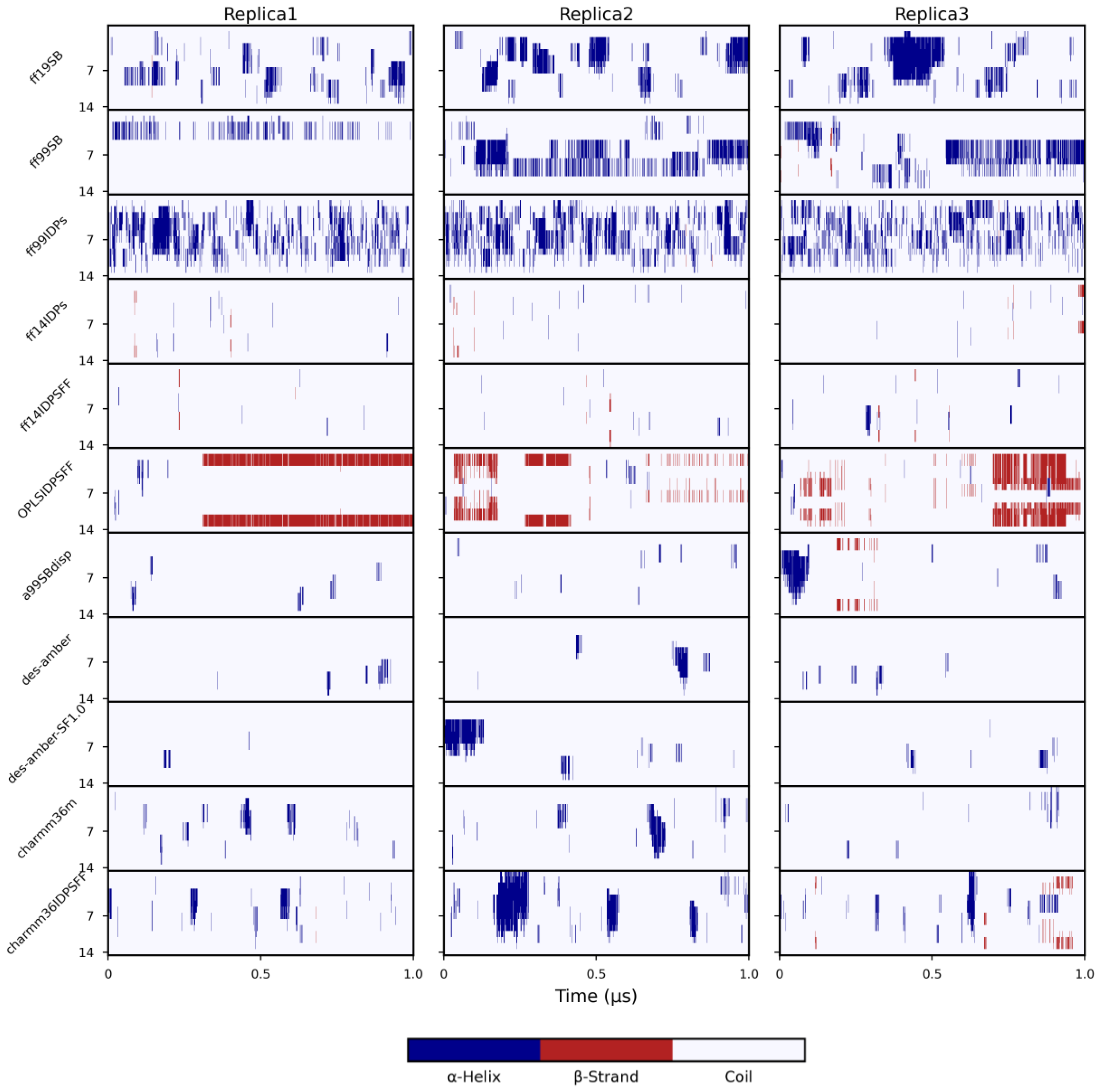


Figure S38: **Per-residue secondary structure time series for The EK peptide** for three 1 $\mu$s simulation replicas initiated from the folded structure. Red denotes $\beta$-strand, and blue denotes $\alpha$-helix or 3-10 helix, and the absence of color corresponds to coil or turn regions.

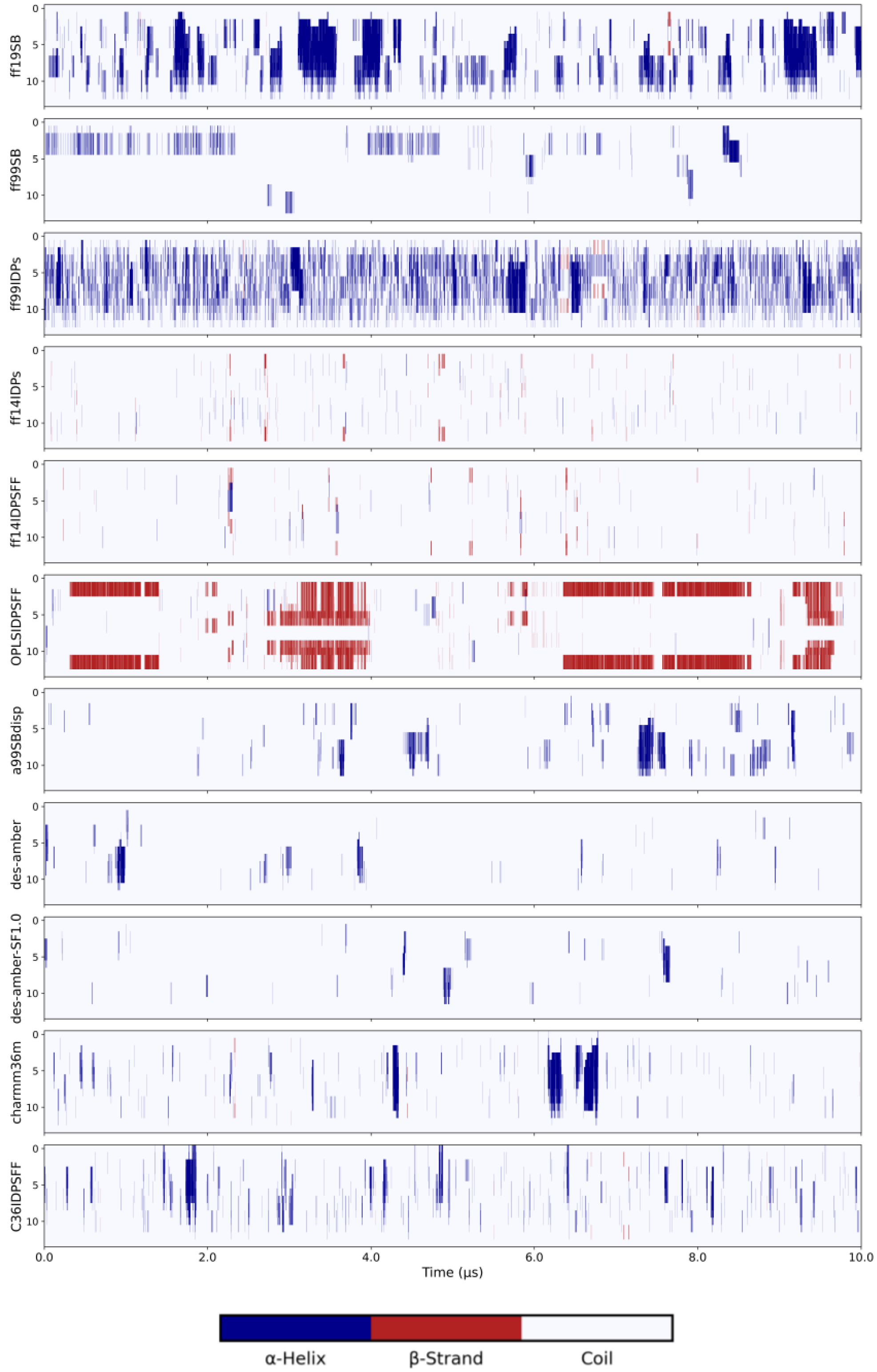


Figure S39: **Per-residue secondary structure time series for The EK peptide** for the 10 $\mu$s simulation replicas initiated from the folded structure. Red denotes $\beta$-strand, and blue denotes $\alpha$-helix or 3-10 helix, and the absence of color corresponds to coil or turn regions.

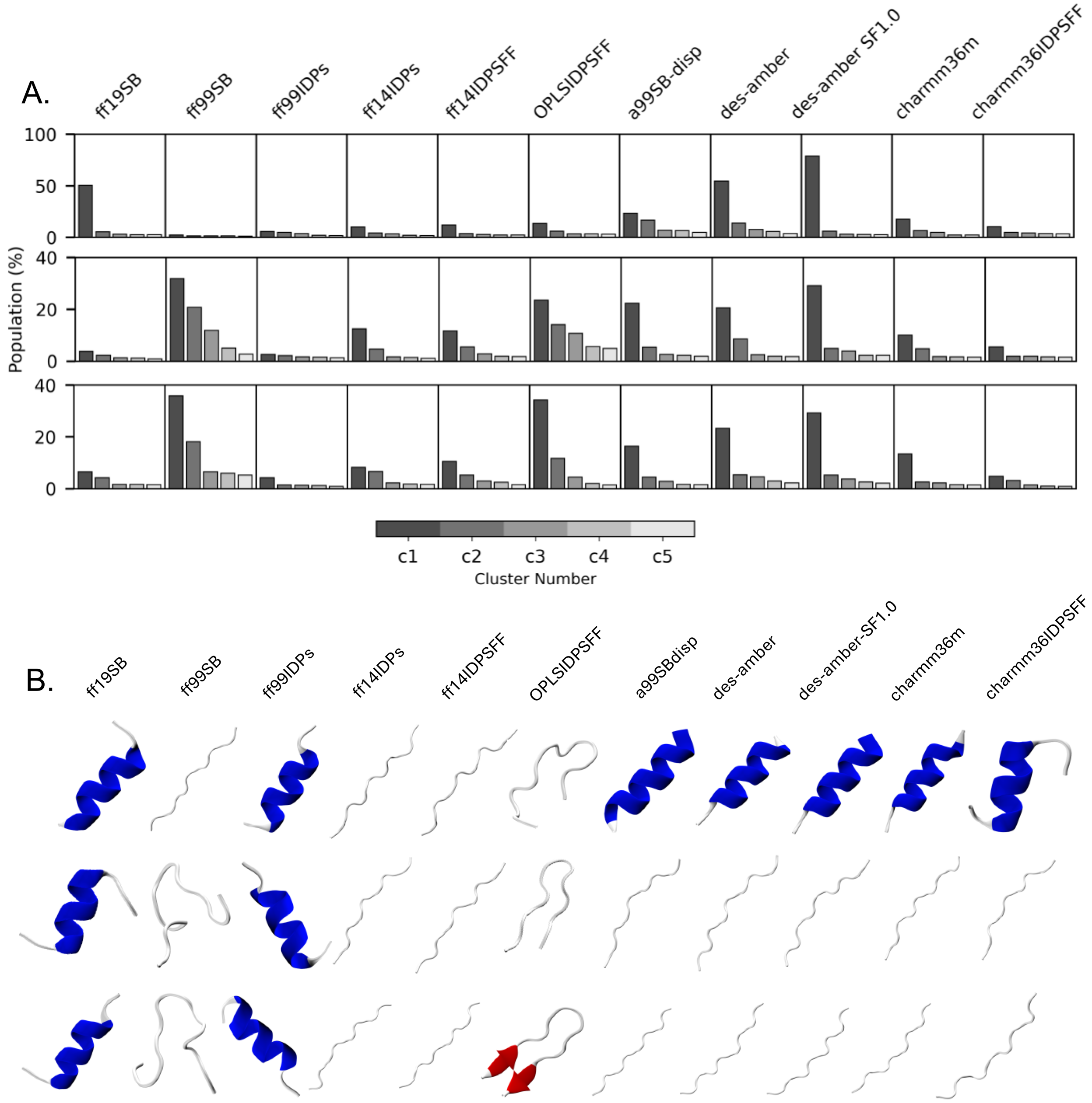


Figure S40: **Clustering results showing (A) cluster populations and (B) representative structures (centroids) of the most populated cluster for The EK peptide.** The top panel shows clustering from five 200-ns stability simulations initiated from the folded structure; the middle panel shows clustering from three 1-µs simulation replicas initiated from the linear peptide sequence; and the bottom panel shows clustering from a 10-µs folding simulation initiated from the extended peptide conformation. Red indicates $\beta$-strand, blue indicates $\alpha$-helix or 3-10 helix, and white indicates coil or turn regions.

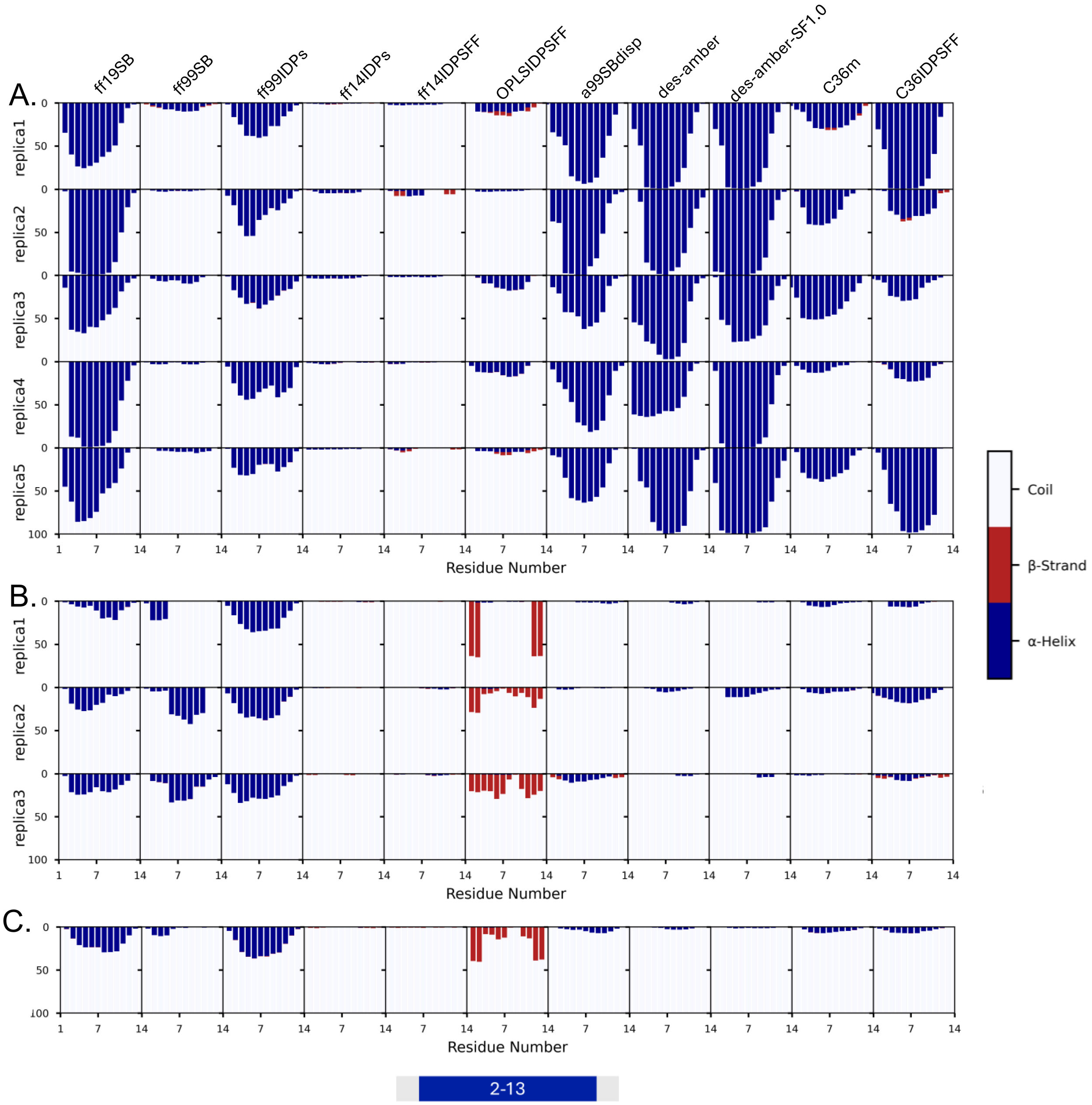


Figure S41: **Per-residue secondary structure preferences for The EK peptide** across simulations. Bars represent the percentage occurrence of each secondary structure type per residue in (A) five 200-ns replicas, (B) three 1-μs replicas, and (C) a 10-μs simulation. The secondary structure assignment in the experimentally folded structure is shown for reference. Red denotes $\beta$-strand, and blue denotes $\alpha$-helix or 3-10 helix, and the absence of color corresponds to coil or turn regions.

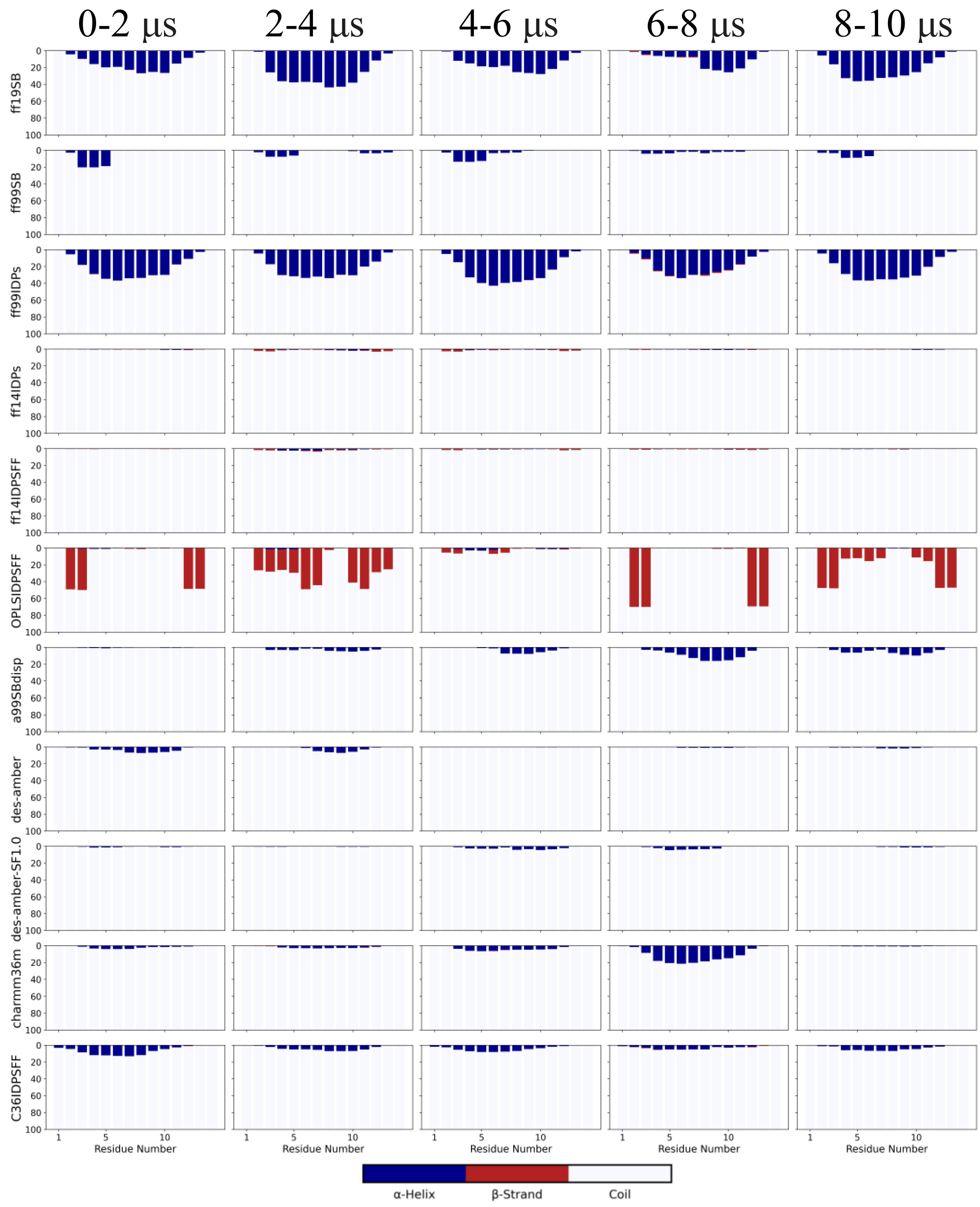


Figure S42: **Per residue secondary structure preferences for The EK peptide during the 10 $\mu$s simulation, shown as blocks of 2 $\mu$s intervals.** Bars represent the percentage occurrence of each secondary structure type per residue. Red indicates $\beta$-strand, blue indicates $\alpha$-helix or 3-10 helix, and the absence of color corresponds to coil or turn regions.

### Structured upon binding

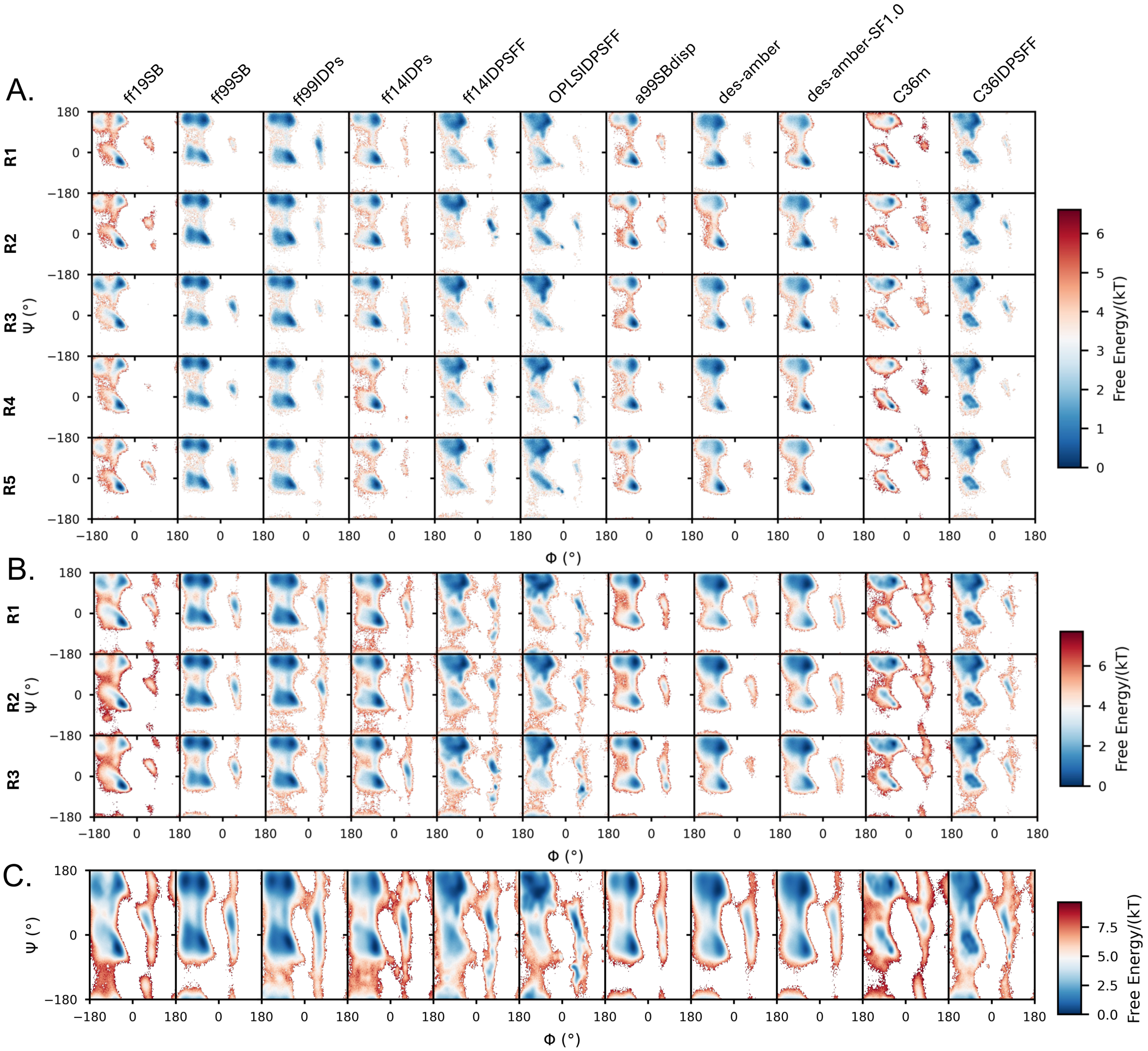


Figure S43: **Ramachandran plots for mat$\alpha$2 (PDB ID 1MNM)**. (A) Five 200-ns simulation replicas initiated from the experimentally folded structure. (B) Three 1-µs simulation replicas initiated from the linear peptide sequence. (C) 10-µs simulation initiated from the linear peptide sequence.

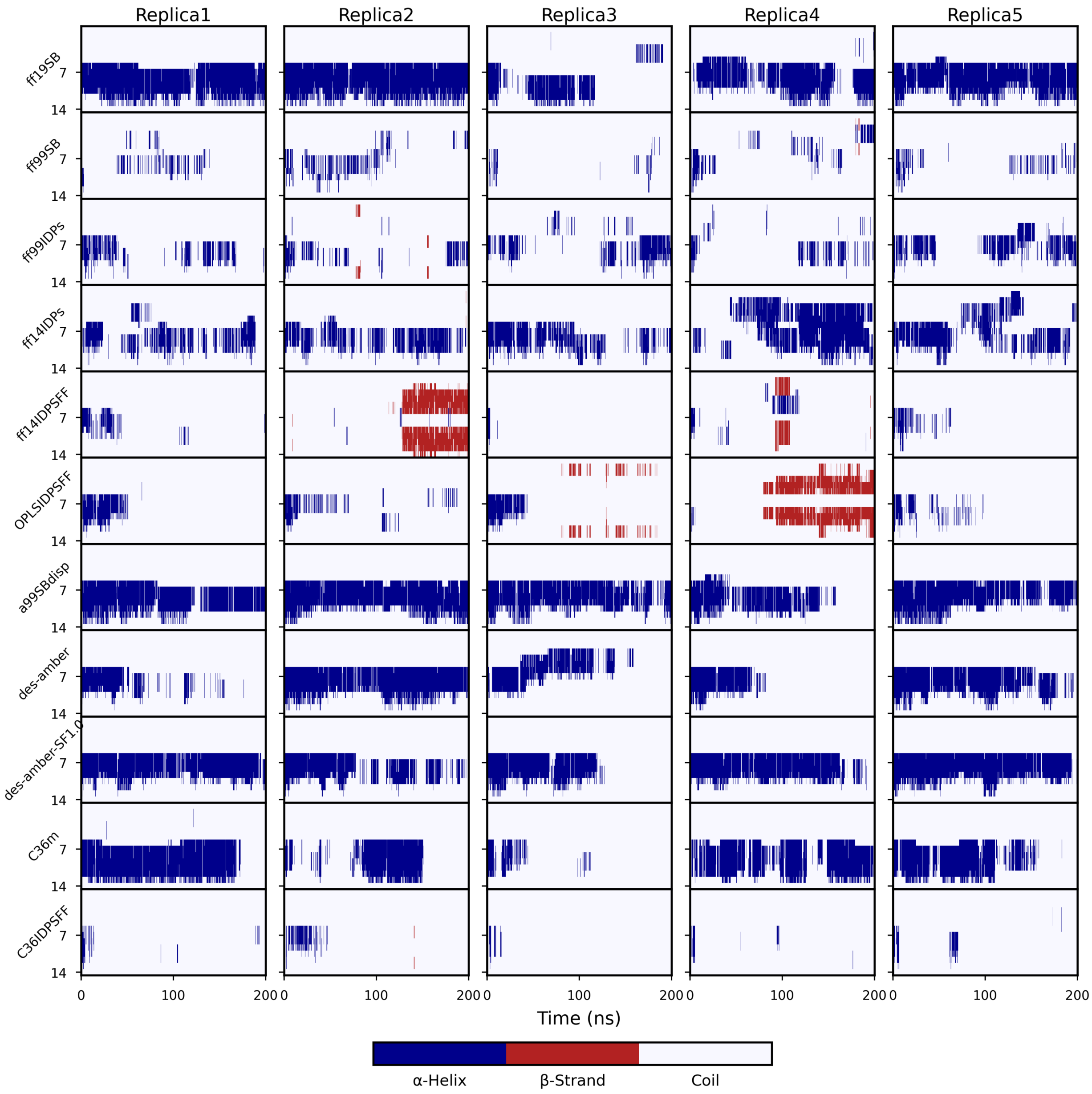


Figure S44: **Per-residue secondary structure time series for mat$\alpha$2 (PDB ID 1MNM)** for five 200-ns simulation replicas initiated from the folded structure. Red denotes $\beta$-strand, and blue denotes $\alpha$-helix or 3-10 helix, and the absence of color corresponds to coil or turn regions.

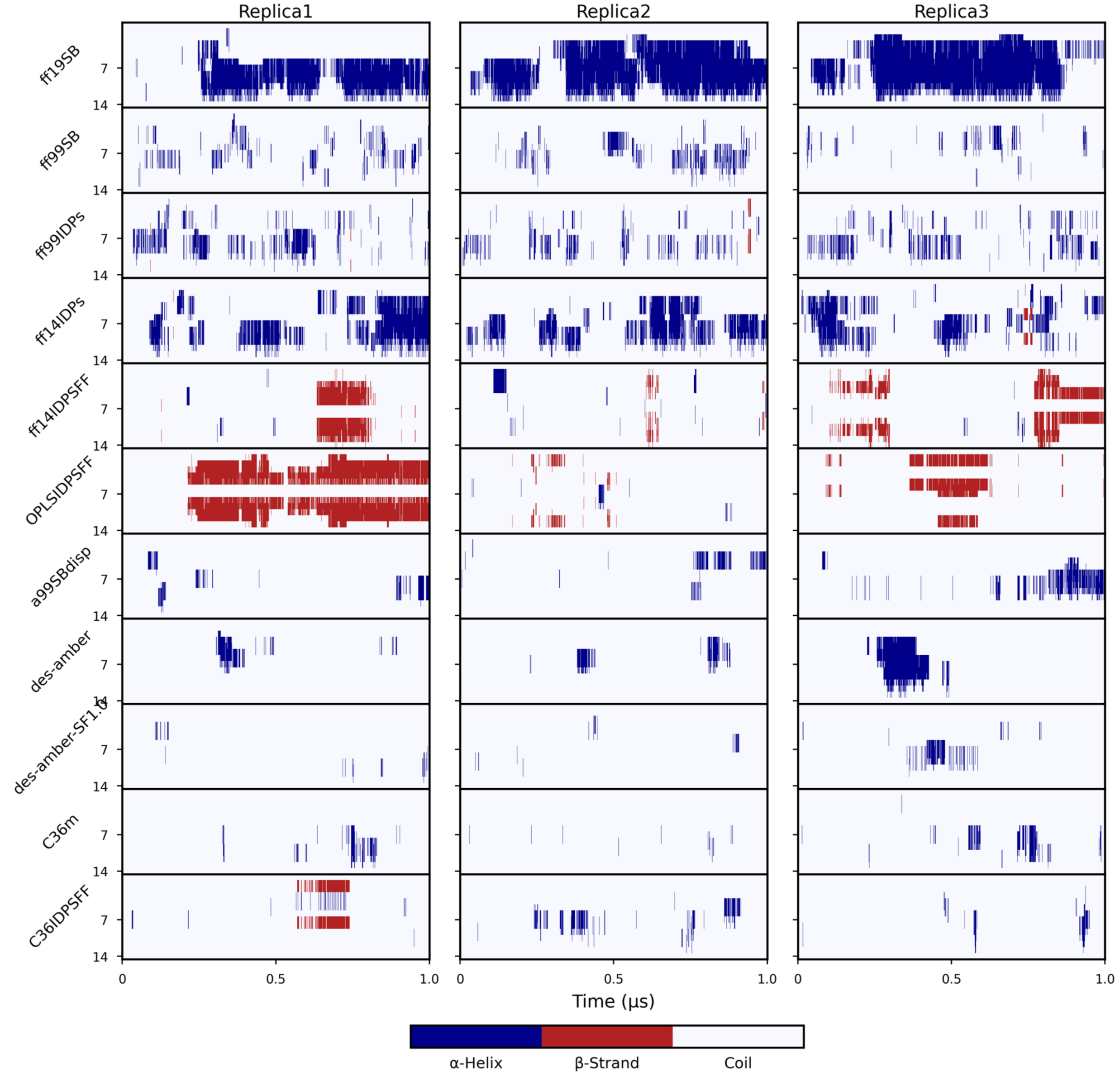


Figure S45: **Per-residue secondary structure time series for mat$\alpha$2 (PDB ID 1MNM)** for three 1 $\mu$s simulation replicas initiated from the folded structure. Red denotes $\beta$-strand, and blue denotes $\alpha$-helix or 3-10 helix, and the absence of color corresponds to coil or turn regions.

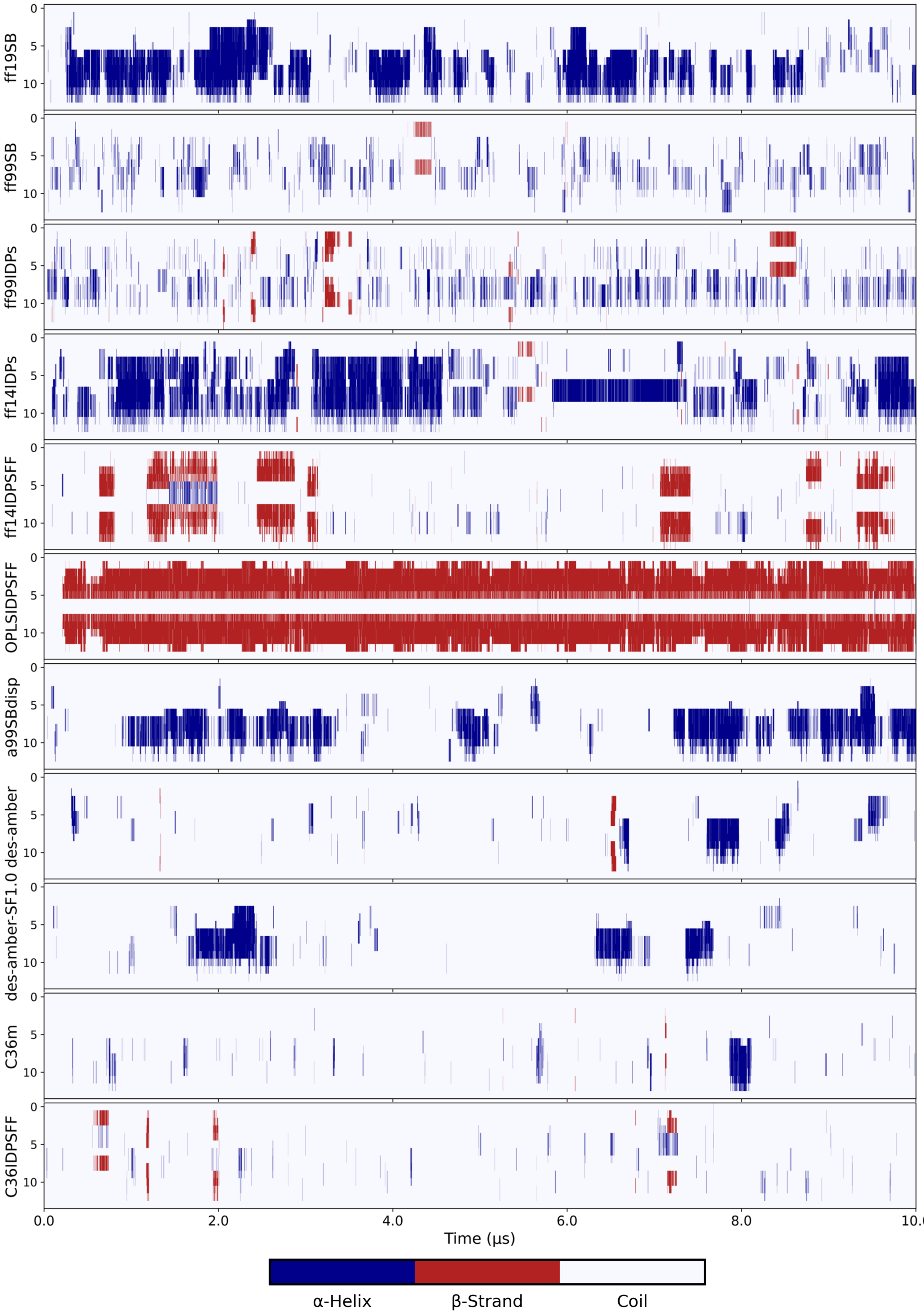


Figure S46: **Per-residue secondary structure time series for mat$\alpha$2 (PDB ID 1MNM)** for the 10 $\mu$s simulation replicas initiated from the folded structure. Red denotes $\beta$-strand, and blue denotes $\alpha$-helix or 3-10 helix, and the absence of color corresponds to coil or turn regions.

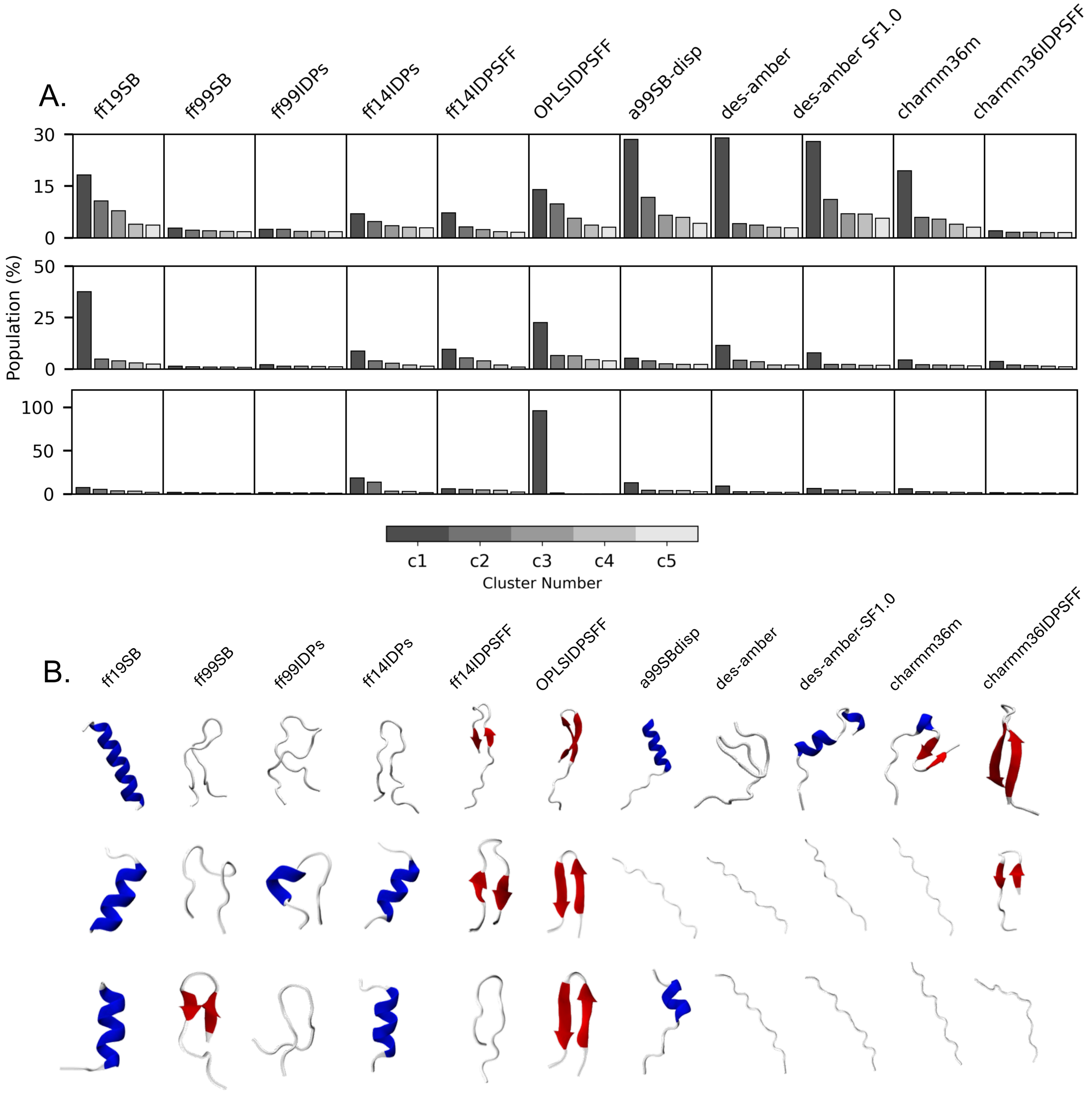


Figure S47: **Clustering results showing (A) cluster populations and (B) representative structures (centroids) of the most populated cluster for mat$\alpha$2 (PDB ID 1MNM).** The top panel shows clustering from five 200-ns stability simulations initiated from the folded structure; the middle panel shows clustering from three 1-µs simulation replicas initiated from the linear peptide sequence; and the bottom panel shows clustering from a 10-µs folding simulation initiated from the extended peptide conformation. Red indicates $\beta$-strand, blue indicates $\alpha$-helix or 3-10 helix, and white indicates coil or turn regions.

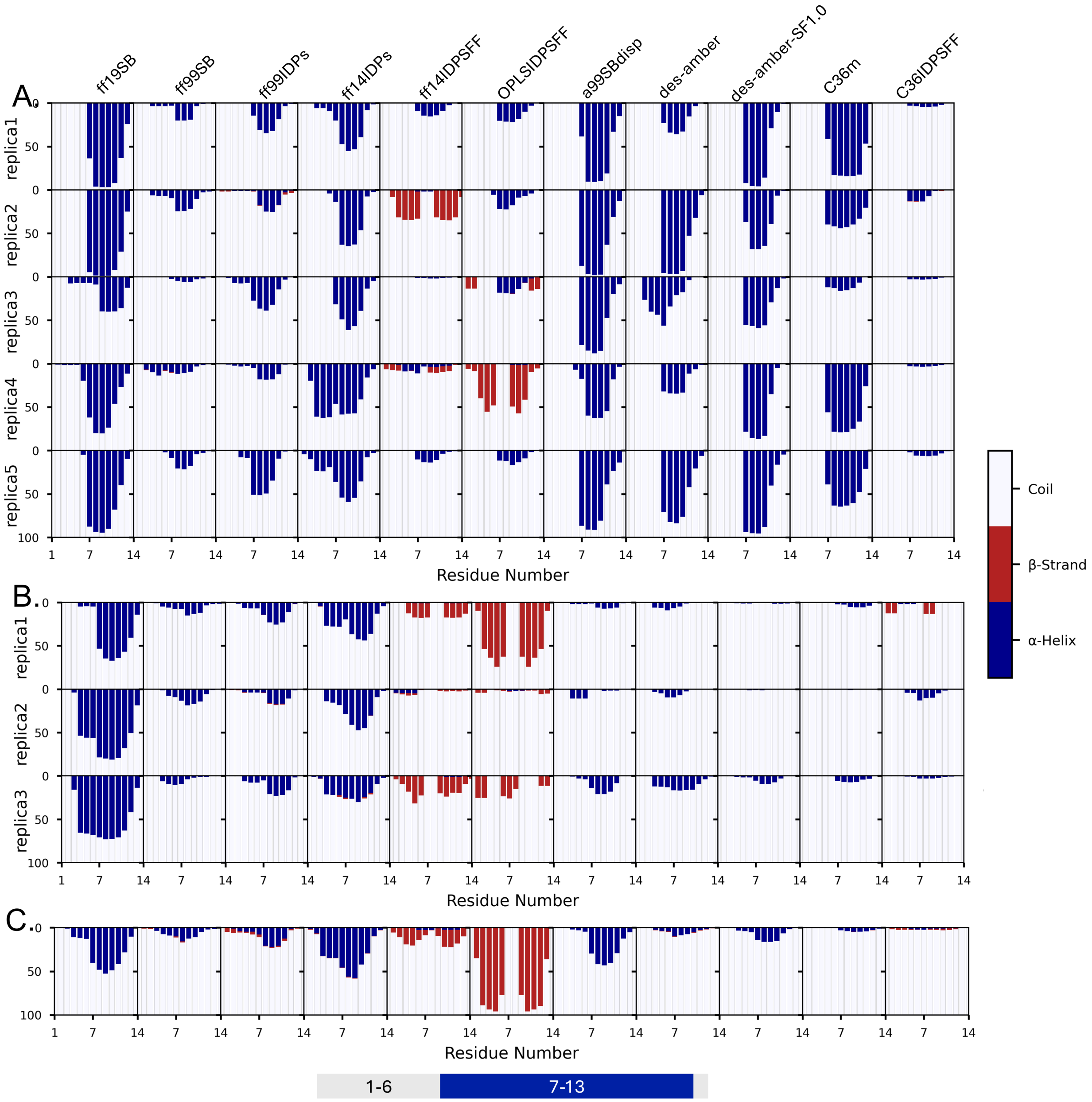


Figure S48: **Per-residue secondary structure preferences for mat$\alpha$2 (PDB ID 1MNM)** across simulations. Bars represent the percentage occurrence of each secondary structure type per residue in (A) five 200-ns replicas, (B) three 1-µs replicas, and (C) a 10-µs simulation. The secondary structure assignment in the experimentally folded structure is shown for reference. Red denotes $\beta$-strand, and blue denotes $\alpha$-helix or 3-10 helix, and the absence of color corresponds to coil or turn regions.

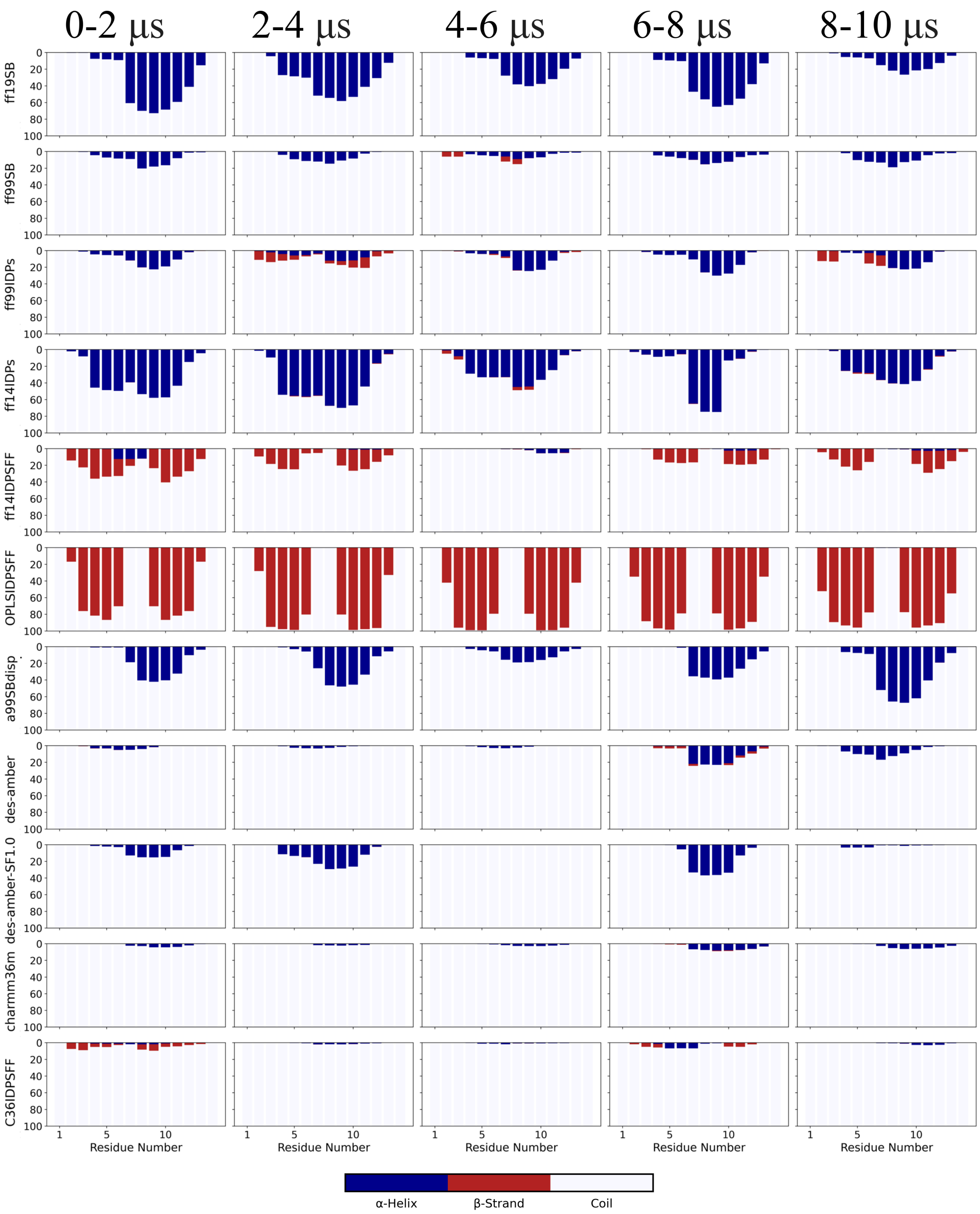


Figure S49: **Per residue secondary structure preferences for mat$\alpha$2 (PDB ID 1MNM) during the 10 $\mu$s simulation, shown as blocks of 2 $\mu$s intervals.** Bars represent the percentage occurrence of each secondary structure type per residue. Red indicates $\beta$-strand, blue indicates $\alpha$-helix or 3-10 helix, and the absence of color corresponds to coil or turn regions.

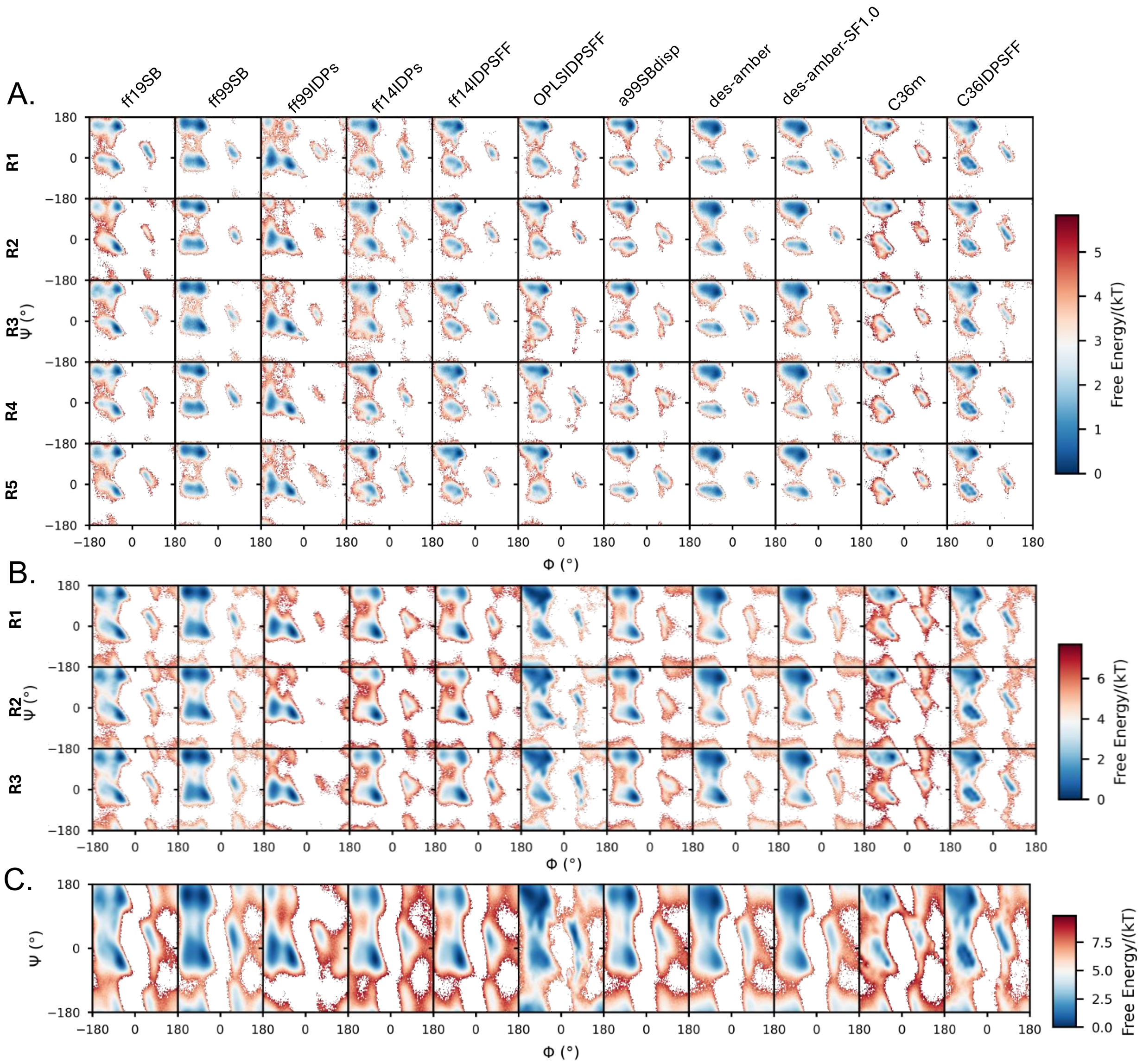


Figure S50: **Ramachandran plots for NRF2 peptide (PDB ID: 2FLU)**. (A) Five 200-ns simulation replicas initiated from the experimentally folded structure. (B) Three 1-μs simulation replicas initiated from the linear peptide sequence. (C) 10-μs simulation initiated from the linear peptide sequence.

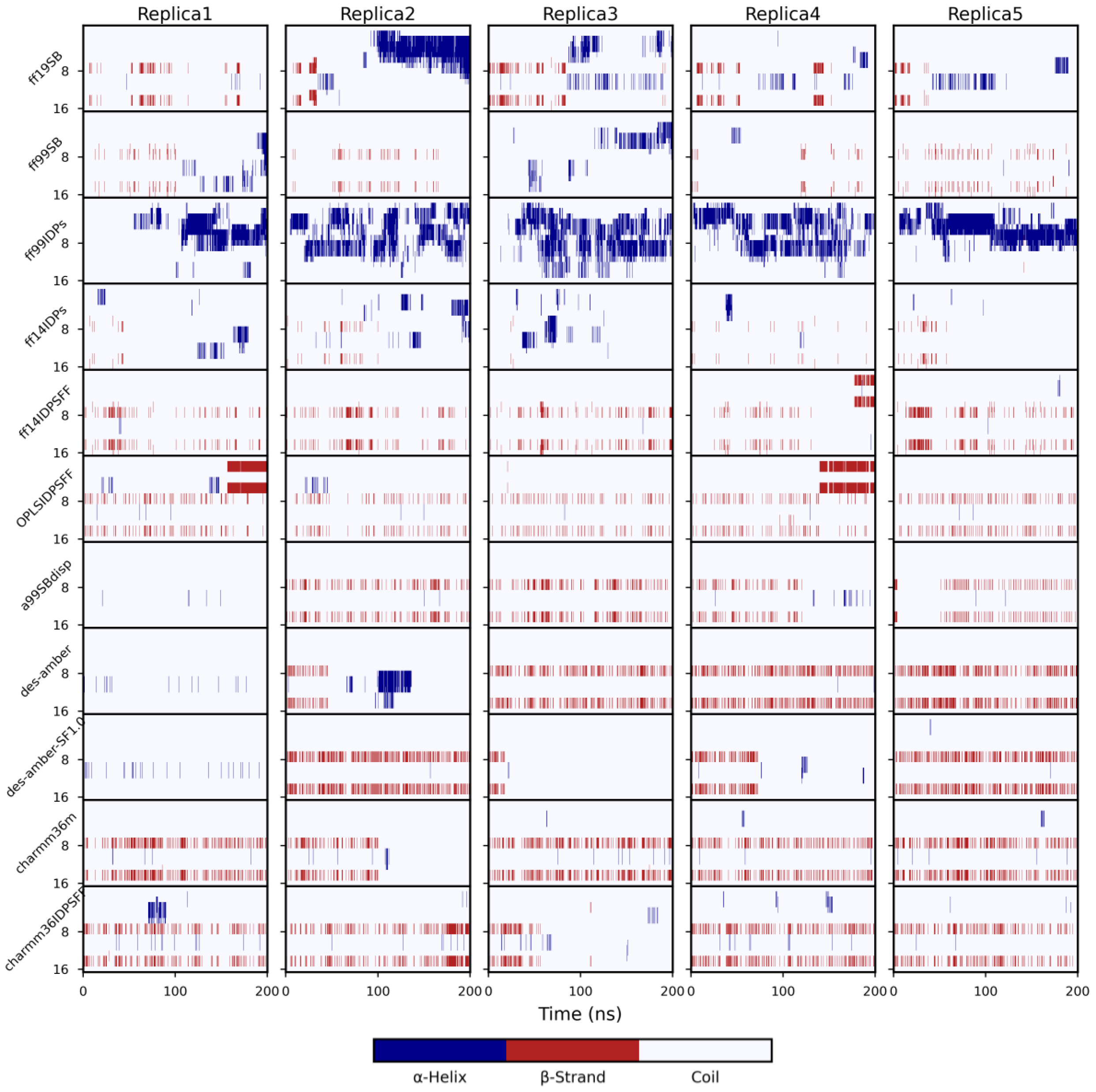


Figure S51: **Per-residue secondary structure time series for NRF2 peptide (PDB ID: 2FLU)** for five 200-ns simulation replicas initiated from the folded structure. Red denotes $\beta$-strand, and blue denotes $\alpha$-helix or 3-10 helix, and the absence of color corresponds to coil or turn regions.

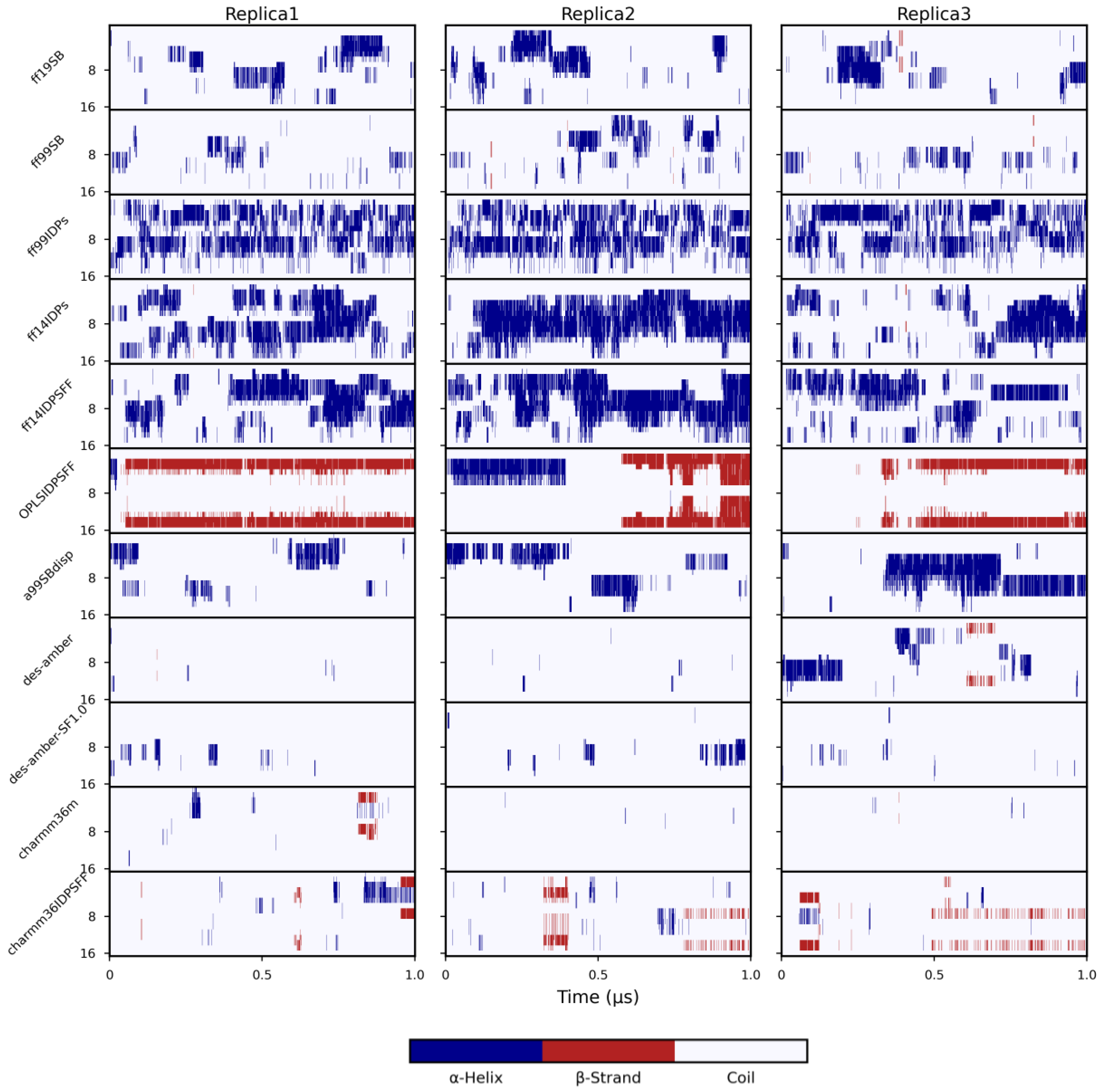


Figure S52: **Per-residue secondary structure time series for NRF2 peptide (PDB ID: 2FLU)** for three 1 $\mu$s simulation replicas initiated from the folded structure. Red denotes $\beta$-strand, and blue denotes $\alpha$-helix or 3-10 helix, and the absence of color corresponds to coil or turn regions.

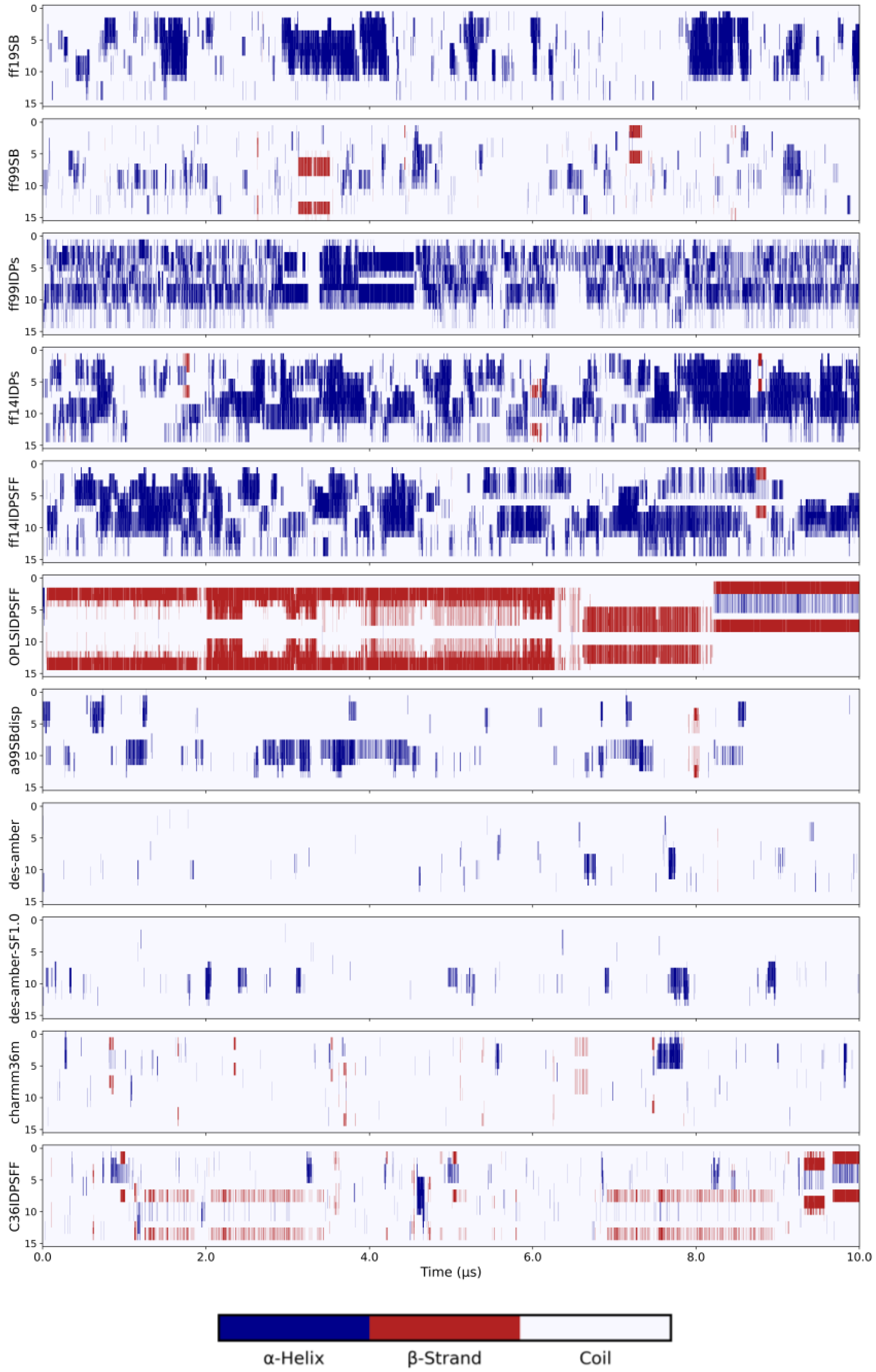


Figure S53: **Per-residue secondary structure time series for NRF2 peptide (PDB ID: 2FLU)** for the 10 $\mu$s simulation replicas initiated from the folded structure. Red denotes $\beta$-strand, and blue denotes $\alpha$-helix or 3-10 helix, and the absence of color corresponds to coil or turn regions.

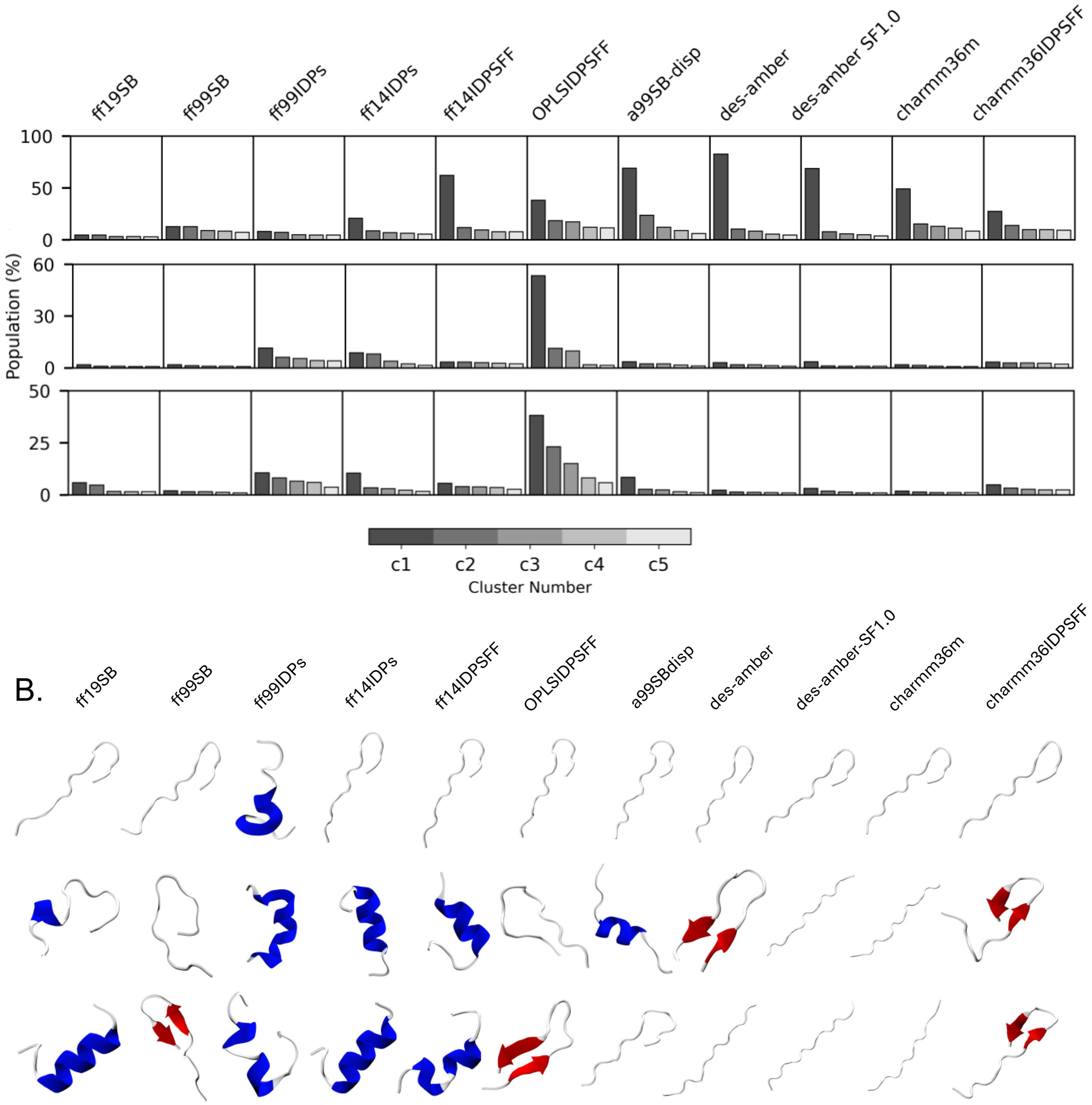


Figure S54: **Clustering results showing (A) cluster populations and (B) representative structures (centroids) of the most populated cluster for NRF2 peptide (PDB ID: 2FLU).** The top panel shows clustering from five 200-ns stability simulations initiated from the folded structure; the middle panel shows clustering from three 1-µs simulation replicas initiated from the linear peptide sequence; and the bottom panel shows clustering from a 10-µs folding simulation initiated from the extended peptide conformation. Red indicates $\beta$-strand, blue indicates $\alpha$-helix or 3-10 helix, and white indicates coil or turn regions.

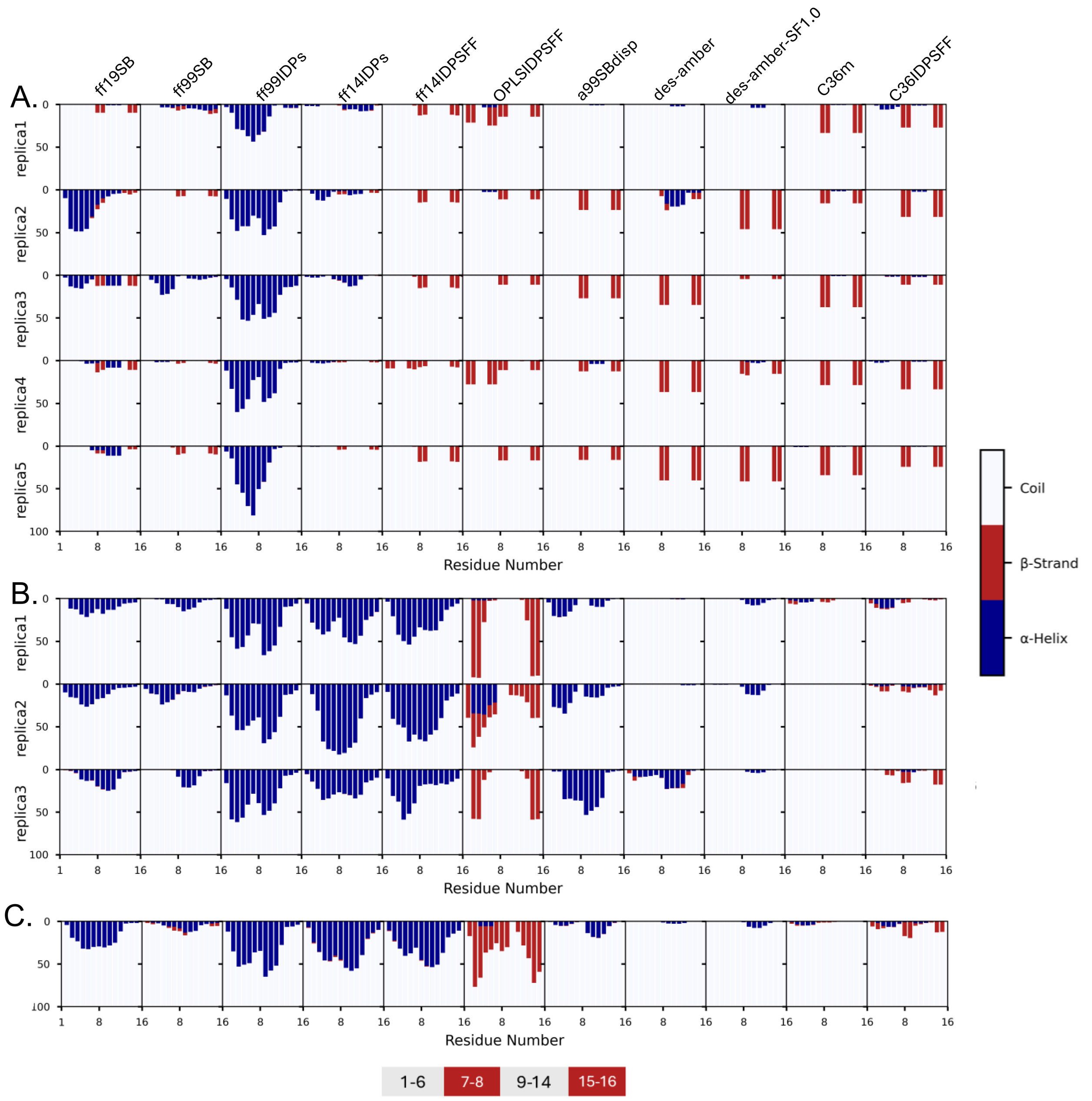


Figure S55: **Per-residue secondary structure preferences for NRF2 peptide (PDB ID: 2FLU)** across simulations. Bars represent the percentage occurrence of each secondary structure type per residue in (A) five 200-ns replicas, (B) three 1-µs replicas, and (C) a 10-µs simulation. The secondary structure assignment in the experimentally folded structure is shown for reference. Red denotes $\beta$-strand, and blue denotes $\alpha$-helix or 3-10 helix, and the absence of color corresponds to coil or turn regions.

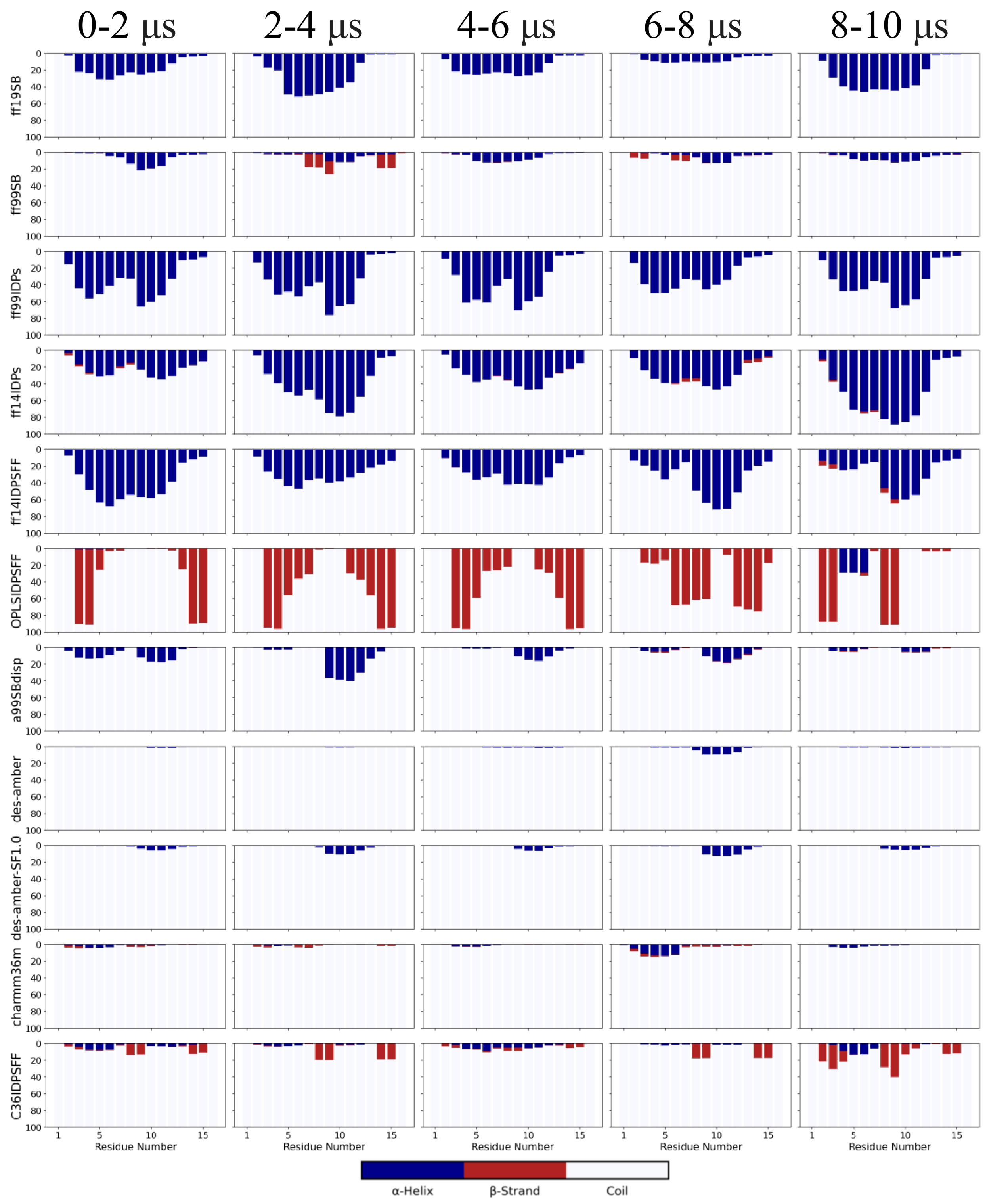


Figure S56: **Per residue secondary structure preferences for NRF2 peptide (PDB ID: 2FLU) during the 10 $\mu$s simulation, shown as blocks of 2 $\mu$s intervals.** Bars represent the percentage occurrence of each secondary structure type per residue. Red indicates $\beta$-strand, blue indicates $\alpha$-helix or 3-10 helix, and the absence of color corresponds to coil or turn regions.

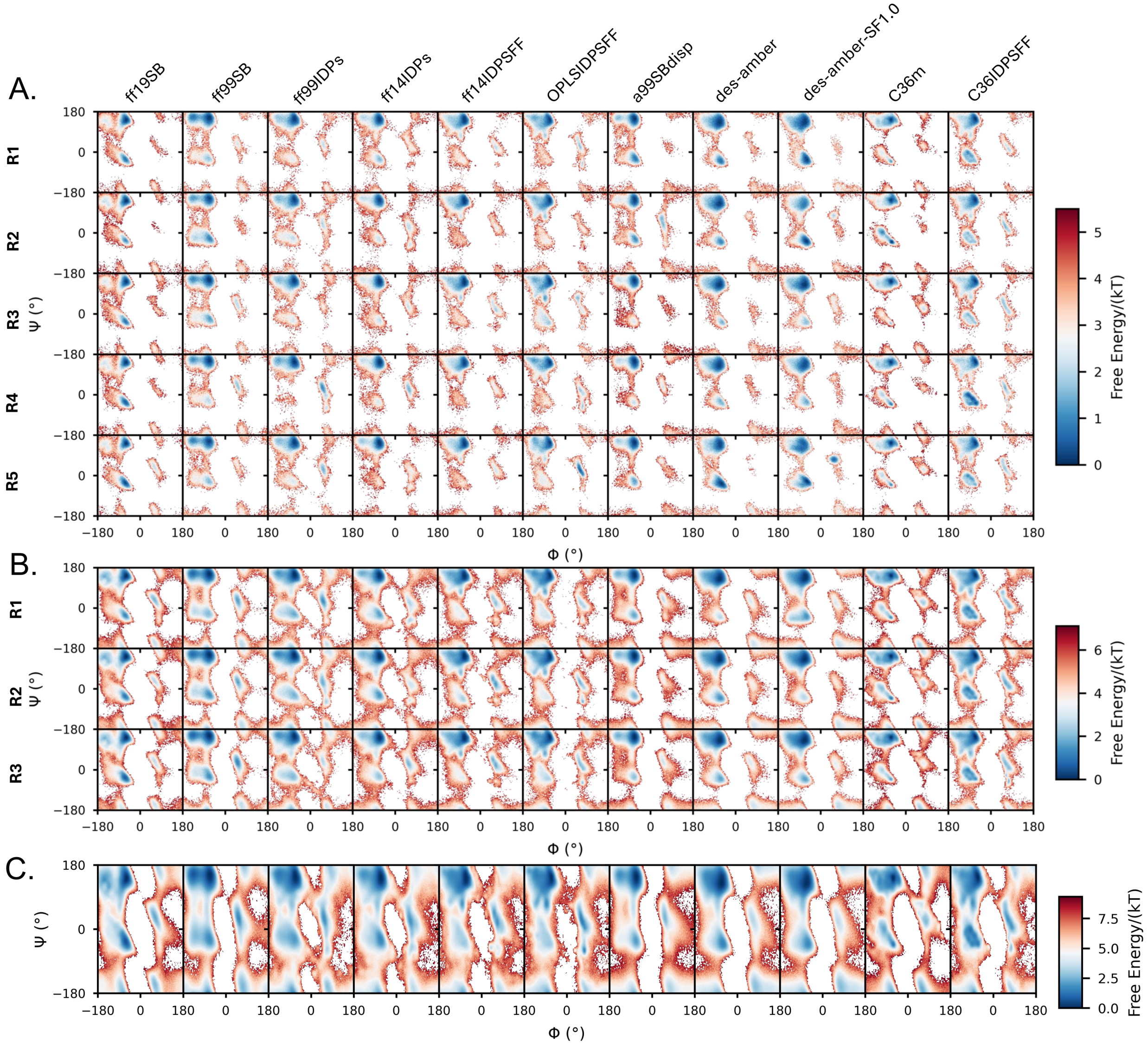


Figure S57: **Ramachandran plots for dynorphin A (PDB ID 2N2F)**. (A) Five 200-ns simulation replicas initiated from the experimentally folded structure. (B) Three 1-μs simulation replicas initiated from the linear peptide sequence. (C) 10-μs simulation initiated from the linear peptide sequence.

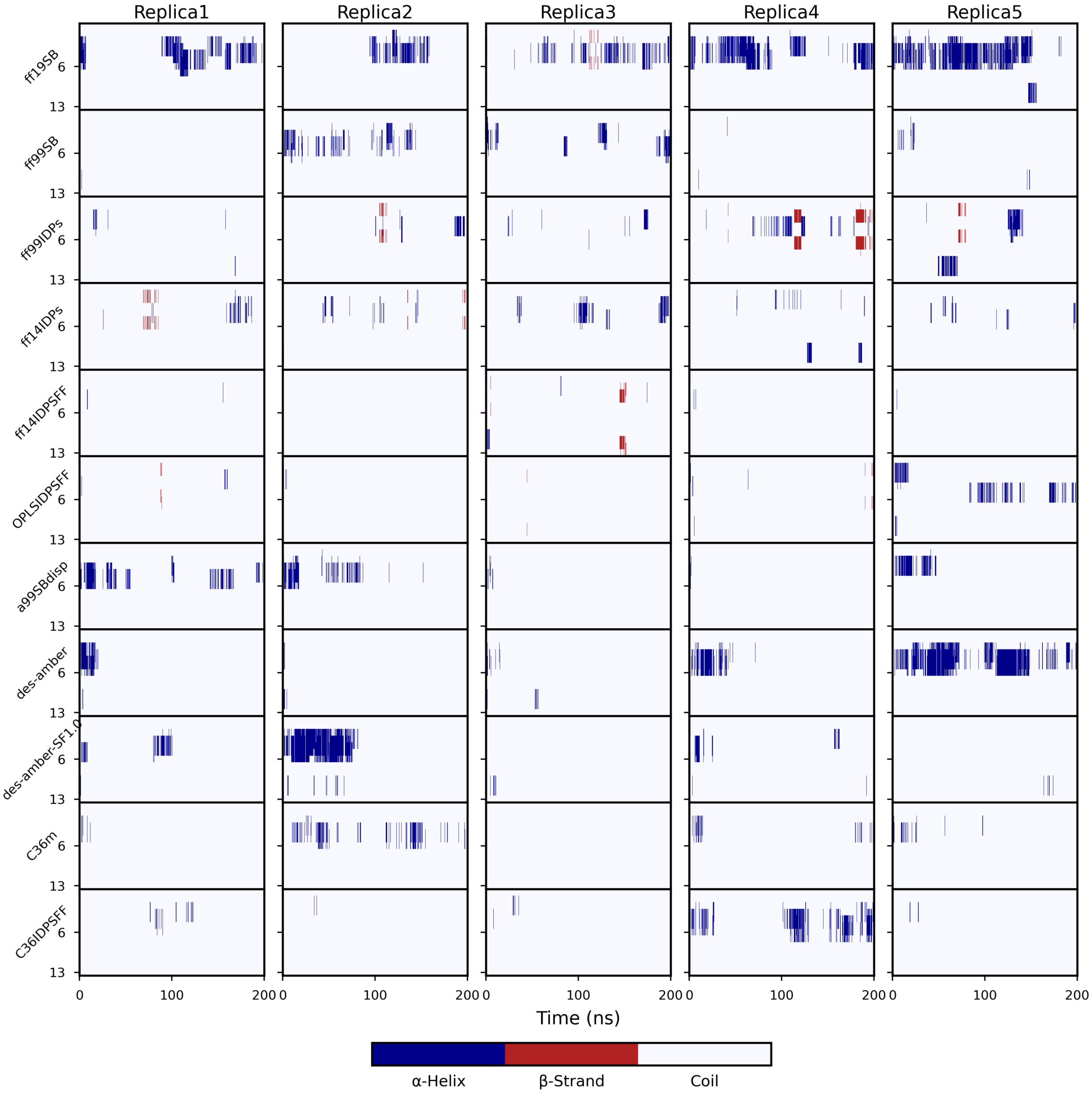


Figure S58: **Per-residue secondary structure time series for dynorphin A (PDB ID 2N2F)** for five 200-ns simulation replicas initiated from the folded structure. Red denotes $\beta$-strand, and blue denotes $\alpha$-helix or 3-10 helix, and the absence of color corresponds to coil or turn regions.

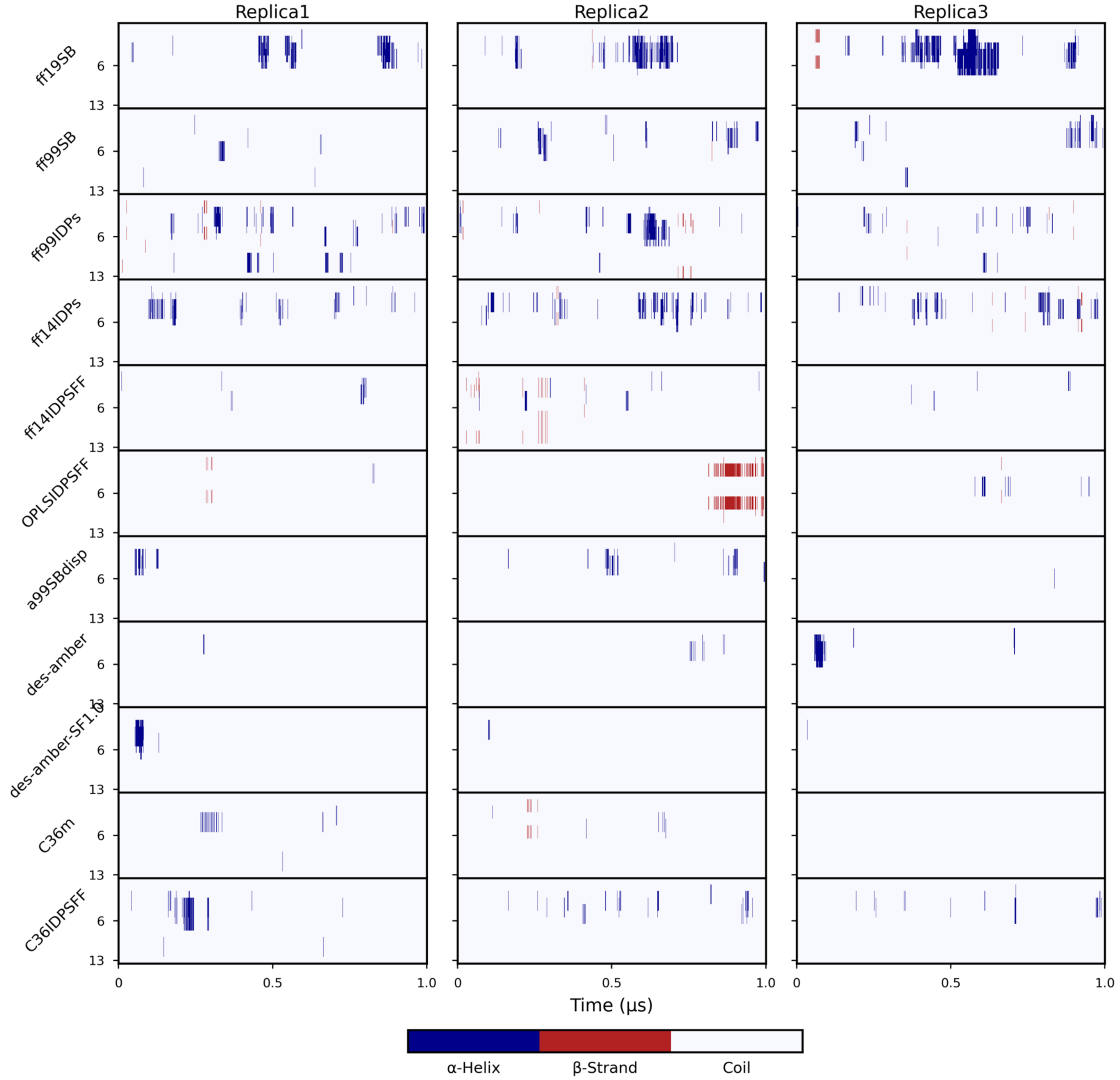


Figure S59: **Per-residue secondary structure time series for dynorphin A (PDB ID 2N2F)** for three 1 $\mu$s simulation replicas initiated from the folded structure. Red denotes $\beta$-strand, and blue denotes $\alpha$-helix or 3-10 helix, and the absence of color corresponds to coil or turn regions.

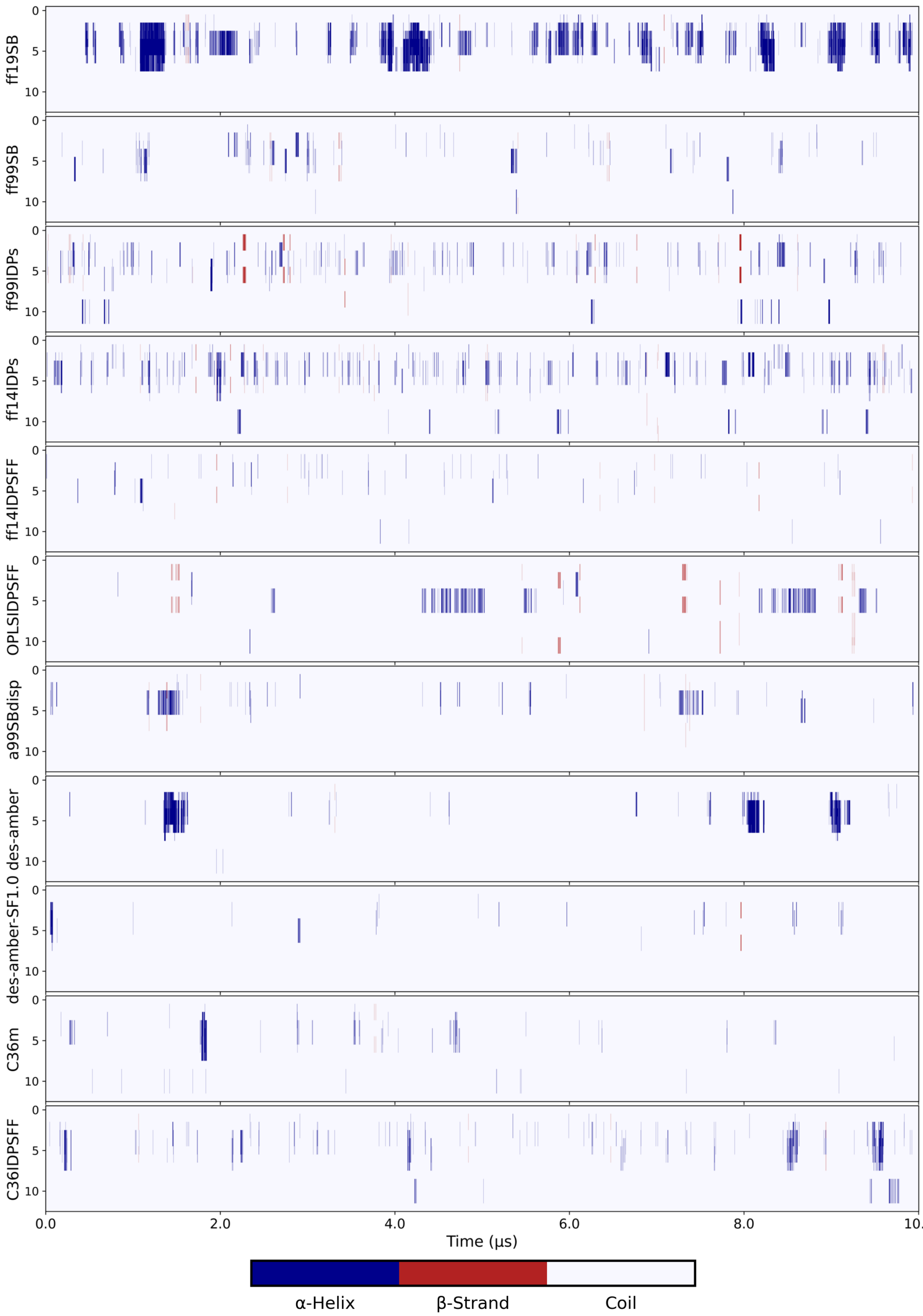


Figure S60: **Per-residue secondary structure time series for dynorphin A (PDB ID 2N2F)** for the 10 $\mu$s simulation replicas initiated from the folded structure. Red denotes $\beta$-strand, and blue denotes $\alpha$-helix or 3-10 helix, and the absence of color corresponds to coil or turn regions.

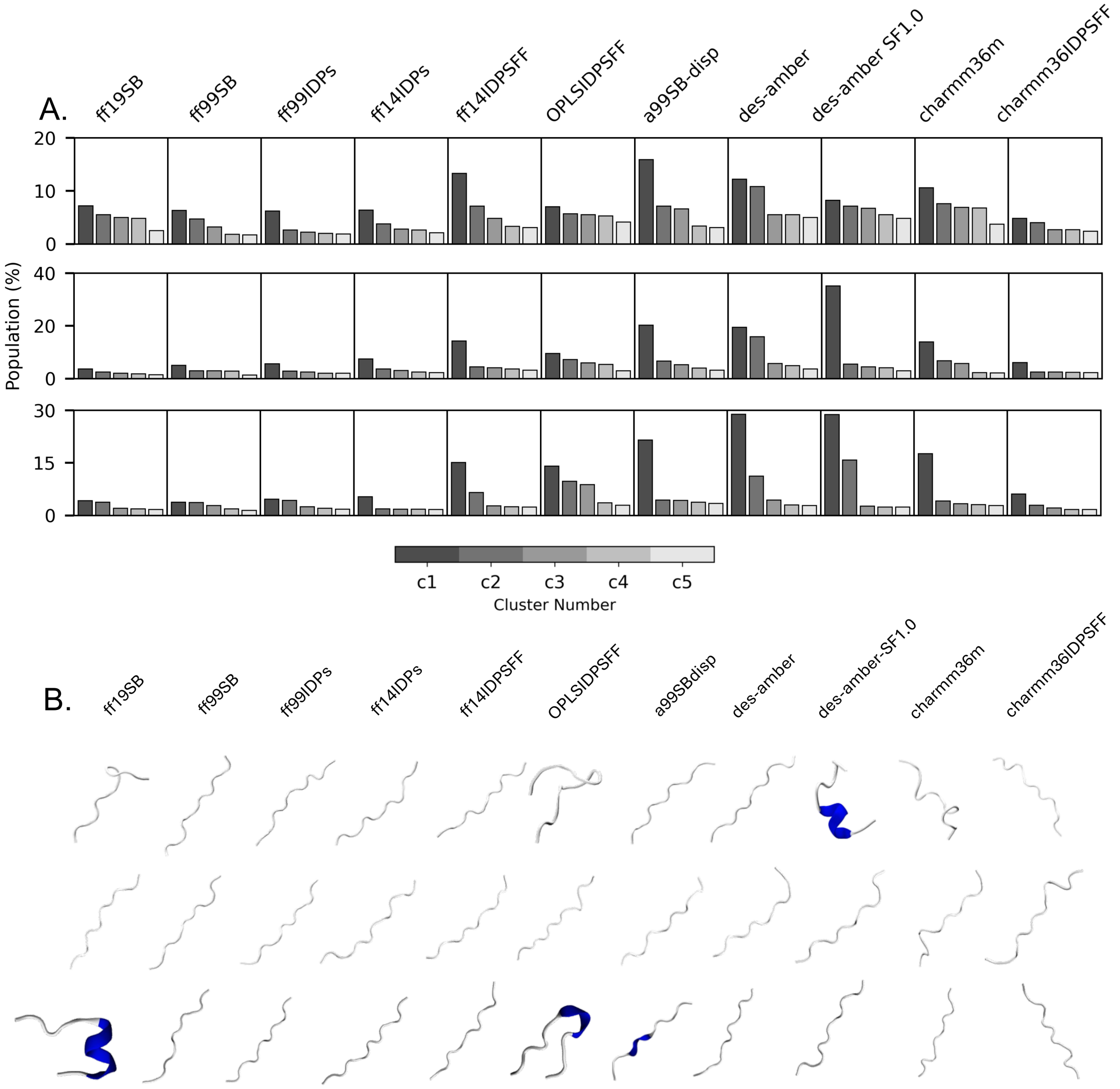


Figure S61: **Clustering results showing (A) cluster populations and (B) representative structures (centroids) of the most populated cluster for dynorphin A (PDB ID 2N2F).** The top panel shows clustering from five 200-ns stability simulations initiated from the folded structure; the middle panel shows clustering from three 1-µs simulation replicas initiated from the linear peptide sequence; and the bottom panel shows clustering from a 10-µs folding simulation initiated from the extended peptide conformation. Red indicates $\beta$-strand, blue indicates $\alpha$-helix or 3-10 helix, and white indicates coil or turn regions.

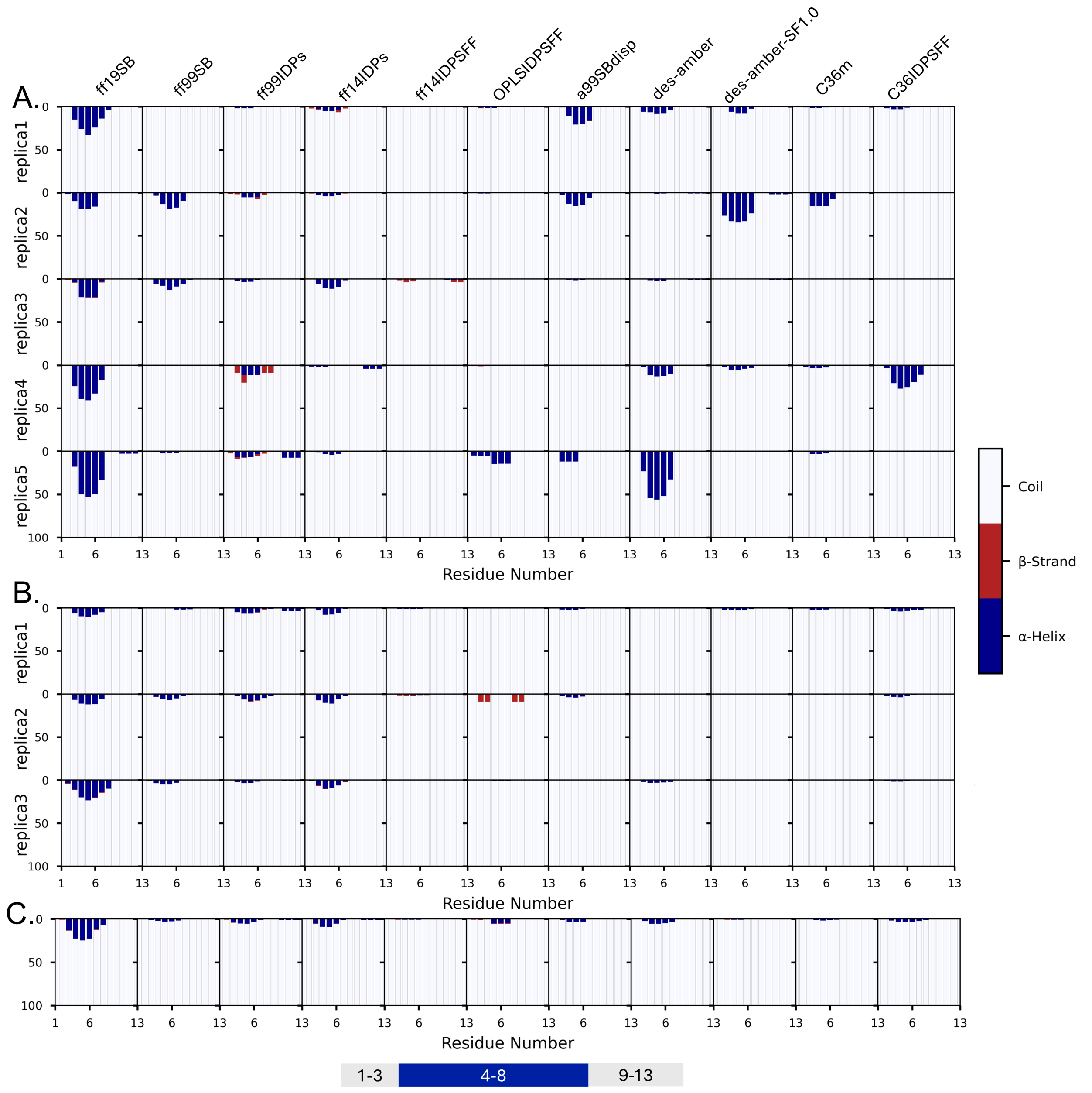


Figure S62: **Per-residue secondary structure preferences for dynorphin A (PDB ID 2N2F)** across simulations. Bars represent the percentage occurrence of each secondary structure type per residue in (A) five 200-ns replicas, (B) three 1-μs replicas, and (C) a 10-μs simulation. The secondary structure assignment in the experimentally folded structure is shown for reference. Red denotes $\beta$-strand, and blue denotes $\alpha$-helix or 3-10 helix, and the absence of color corresponds to coil or turn regions.

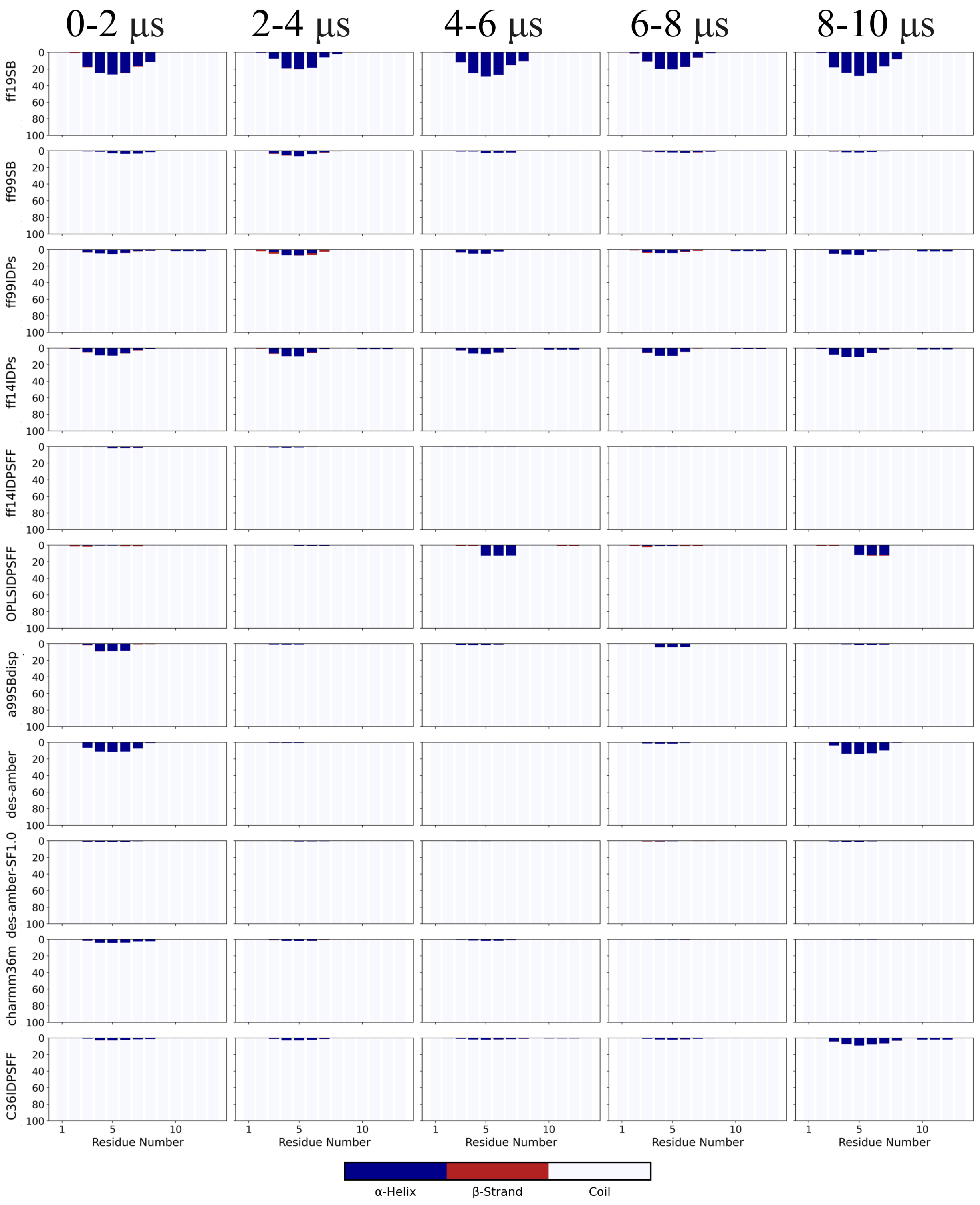


Figure S63: **Per residue secondary structure preferences for dynorphin A (PDB ID 2N2F) during the 10 $\mu$s simulation, shown as blocks of 2 $\mu$s intervals.** Bars represent the percentage occurrence of each secondary structure type per residue. Red indicates $\beta$-strand, blue indicates $\alpha$-helix or 3-10 helix, and the absence of color corresponds to coil or turn regions.

### Disordered in water, ordered in solvent mixtures

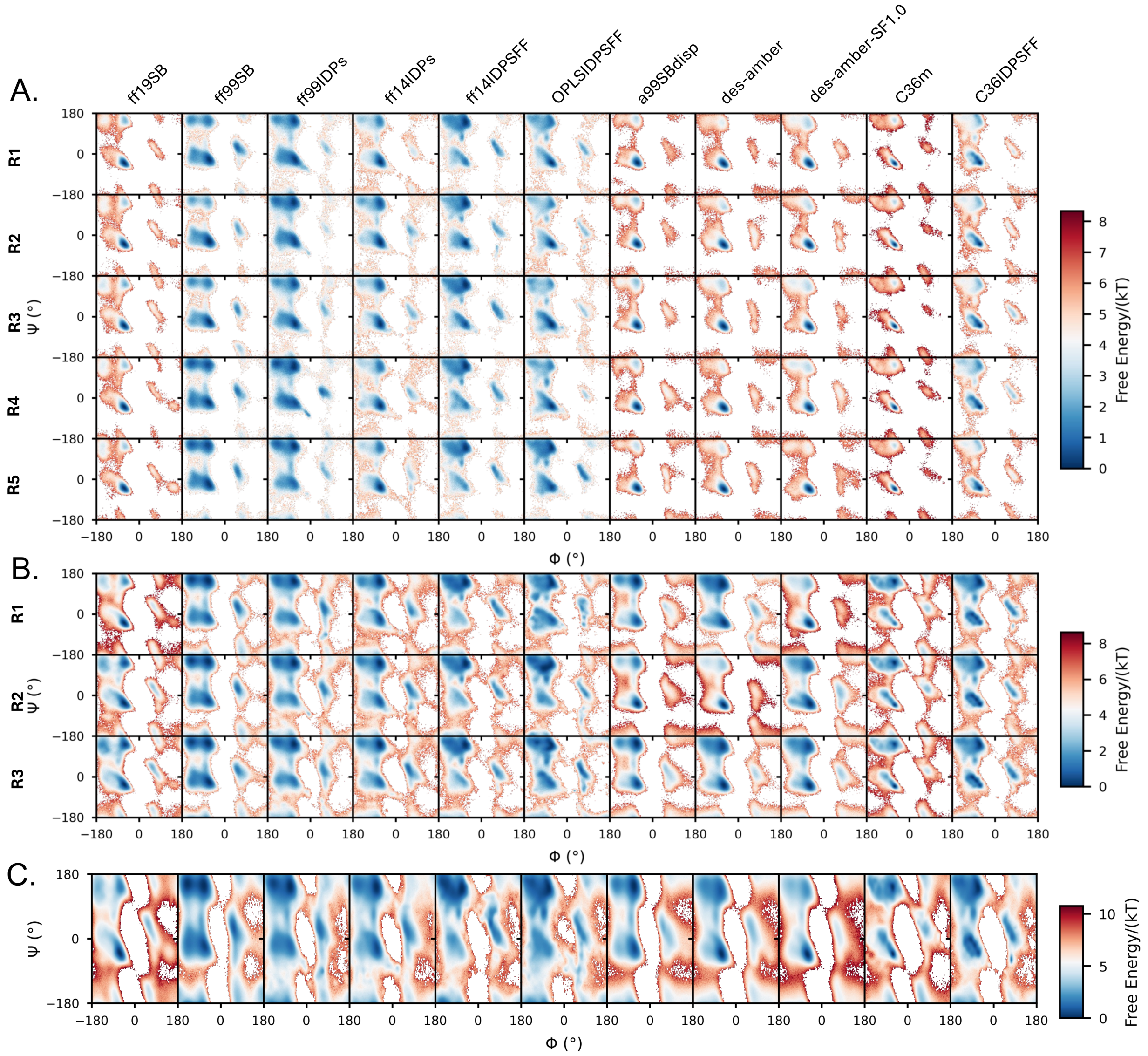


Figure S64: **Ramachandran plots for GLP-1 (PDB ID 1D0R)**. (A) Five 200-ns simulation replicas initiated from the experimentally folded structure. (B) Three 1-µs simulation replicas initiated from the linear peptide sequence. (C) 10-µs simulation initiated from the linear peptide sequence.

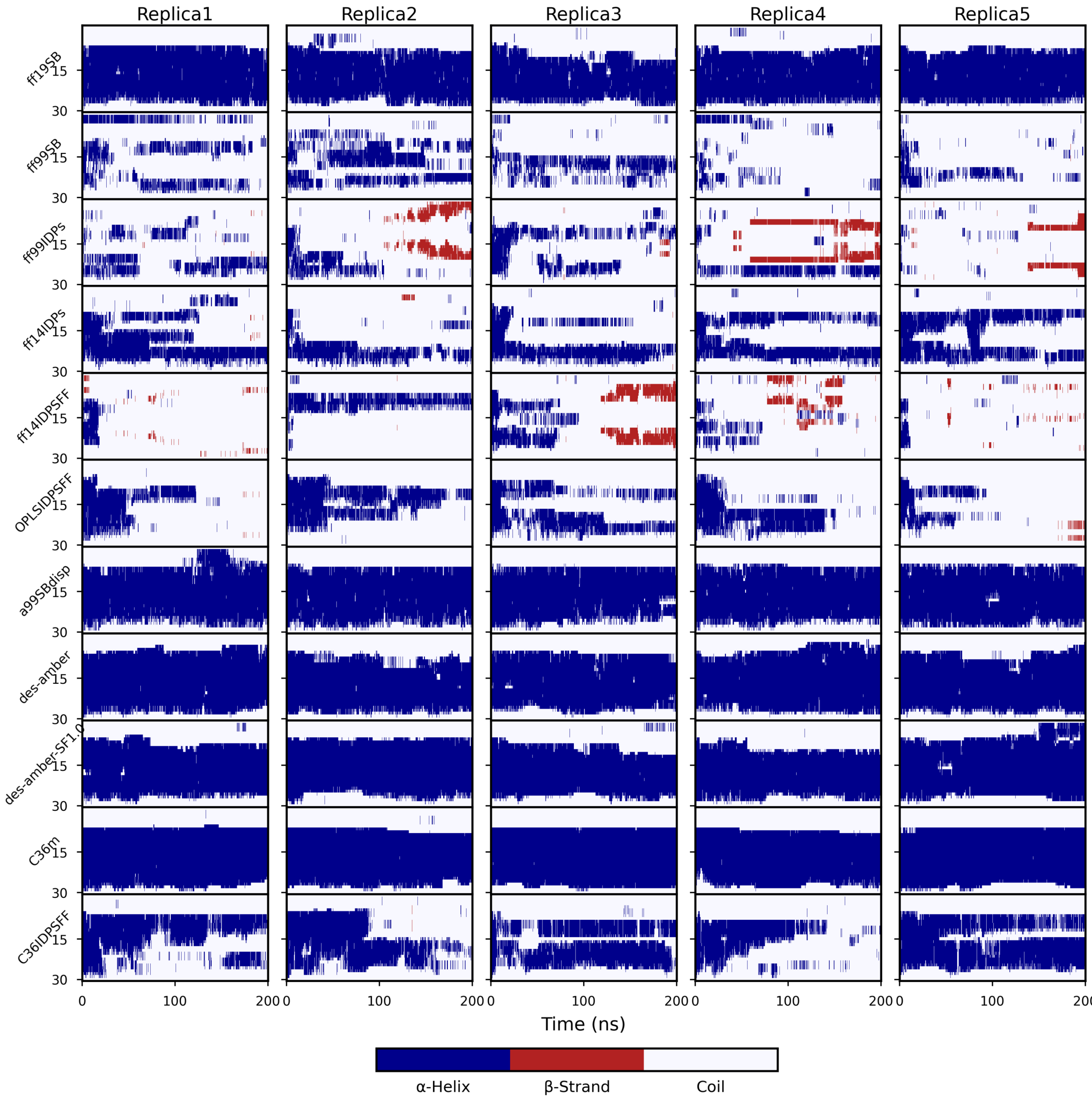


Figure S65: **Per-residue secondary structure time series for GLP-1 (PDB ID 1D0R)** for five 200-ns simulation replicas initiated from the folded structure. Red denotes $\beta$-strand, and blue denotes $\alpha$-helix or 3-10 helix, and the absence of color corresponds to coil or turn regions.

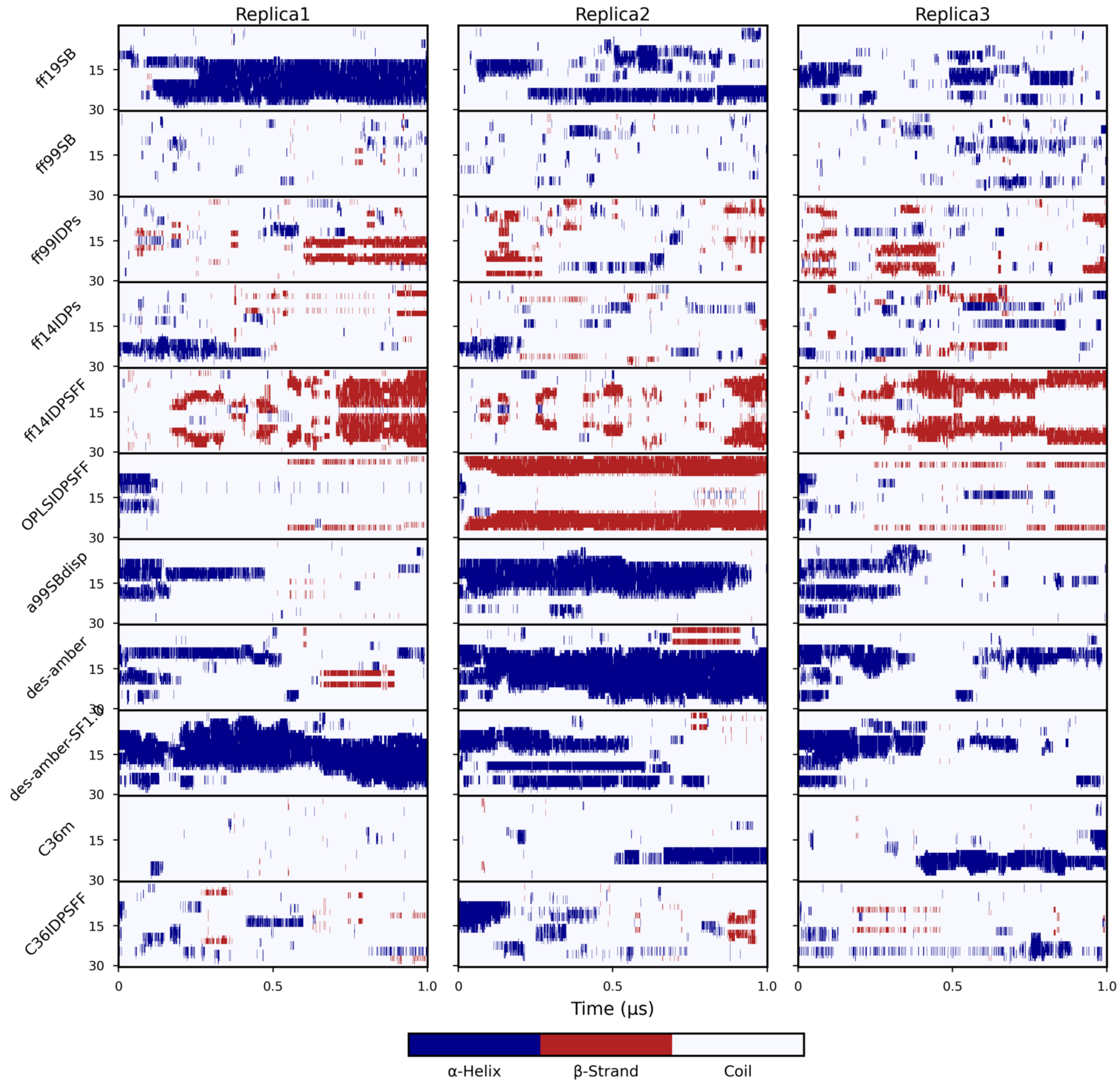


Figure S66: **Per-residue secondary structure time series for GLP-1 (PDB ID 1D0R)** for three 1 $\mu$s simulation replicas initiated from the folded structure. Red denotes $\beta$-strand, and blue denotes $\alpha$-helix or 3-10 helix, and the absence of color corresponds to coil or turn regions.

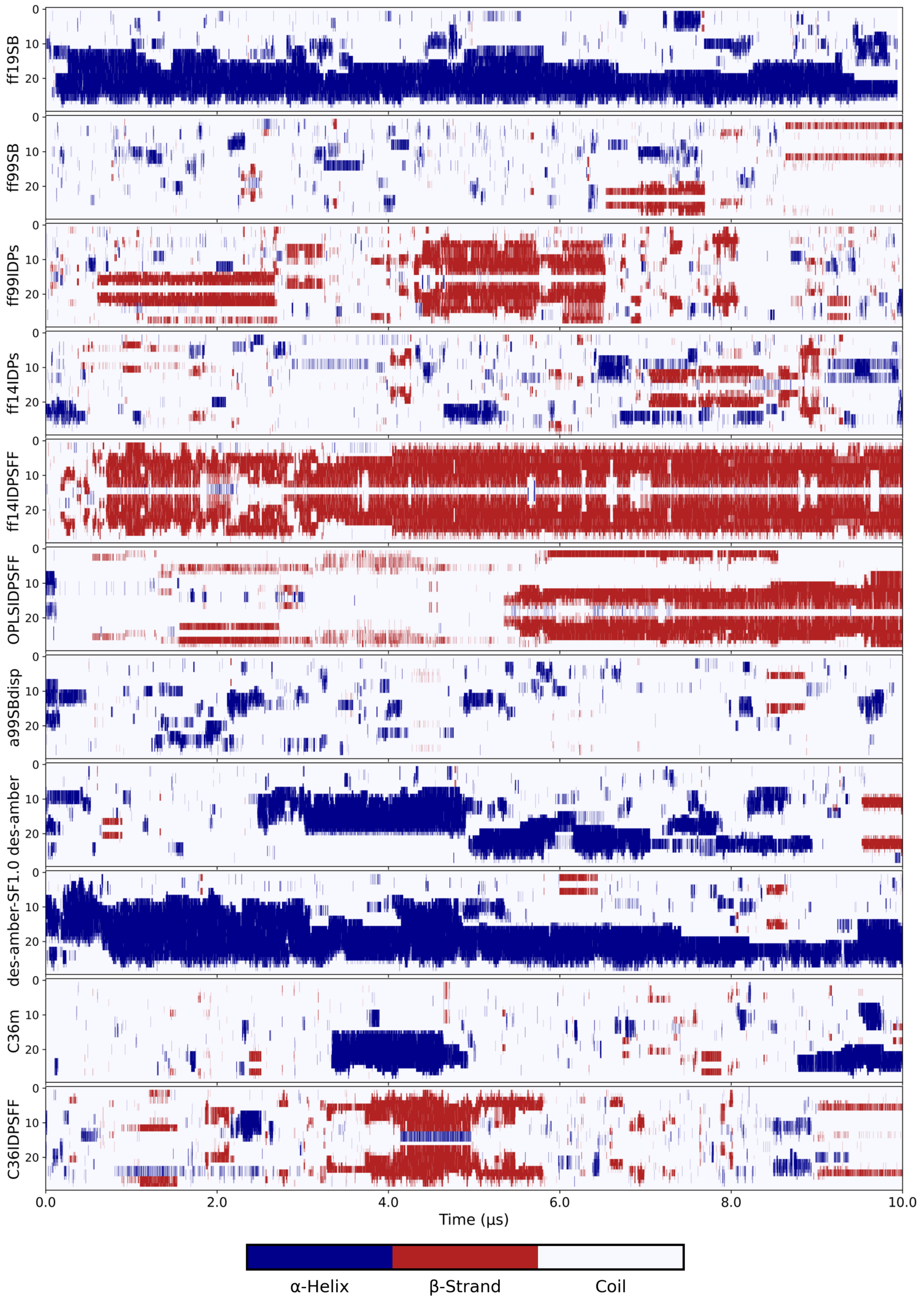


Figure S67: **Per-residue secondary structure time series for GLP-1 (PDB ID 1D0R)** for the 10 $\mu$s simulation replicas initiated from the folded structure. Red denotes $\beta$-strand, and blue denotes $\alpha$-helix or 3-10 helix, and the absence of color corresponds to coil or turn regions.

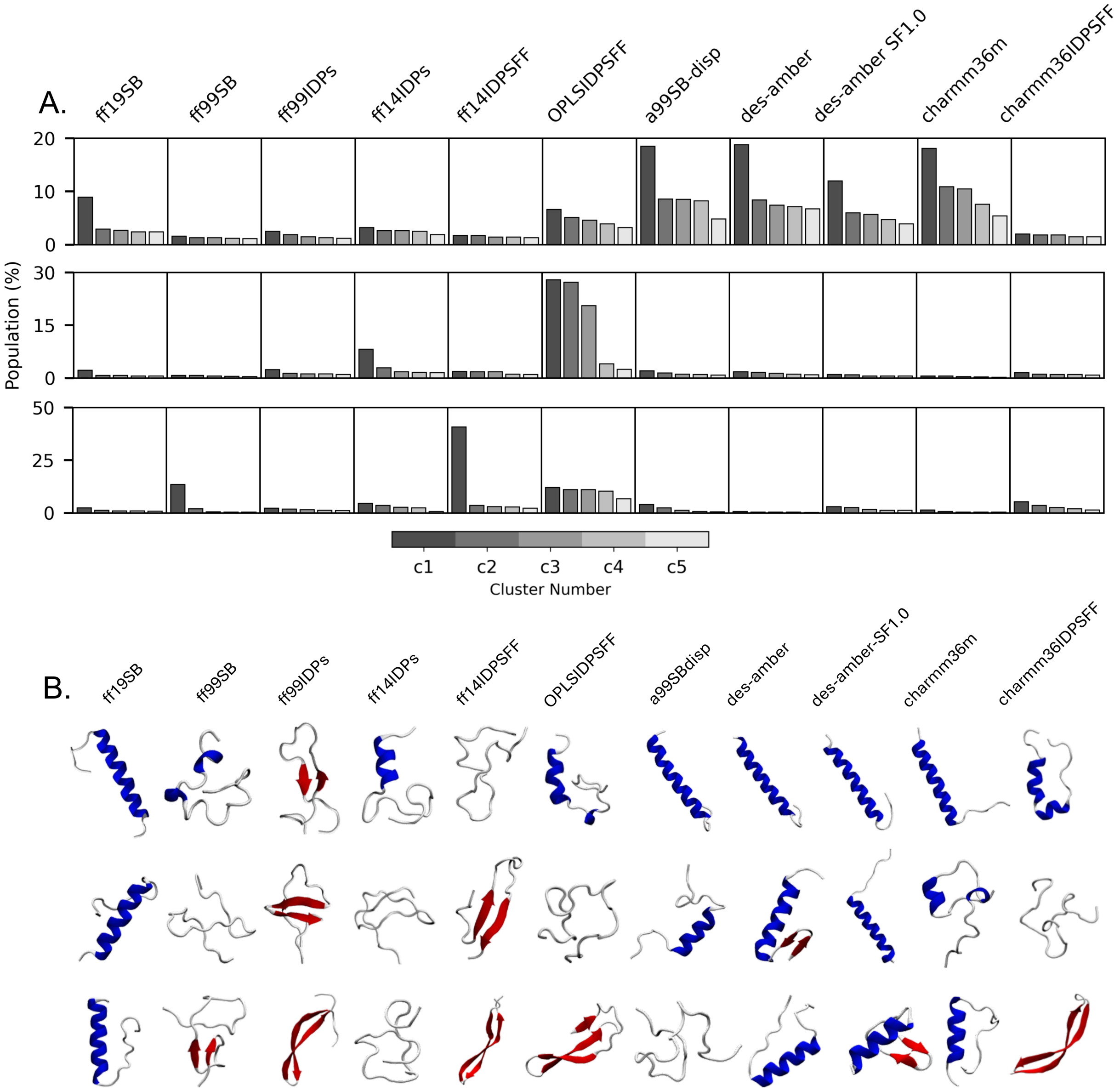


Figure S68: **Clustering results showing (A) cluster populations and (B) representative structures (centroids) of the most populated cluster for GLP-1 (PDB ID 1D0R).** The top panel shows clustering from five 200-ns stability simulations initiated from the folded structure; the middle panel shows clustering from three 1-μs simulation replicas initiated from the linear peptide sequence; and the bottom panel shows clustering from a 10-μs folding simulation initiated from the extended peptide conformation. Red indicates $\beta$-strand, blue indicates $\alpha$-helix or 3-10 helix, and white indicates coil or turn regions.

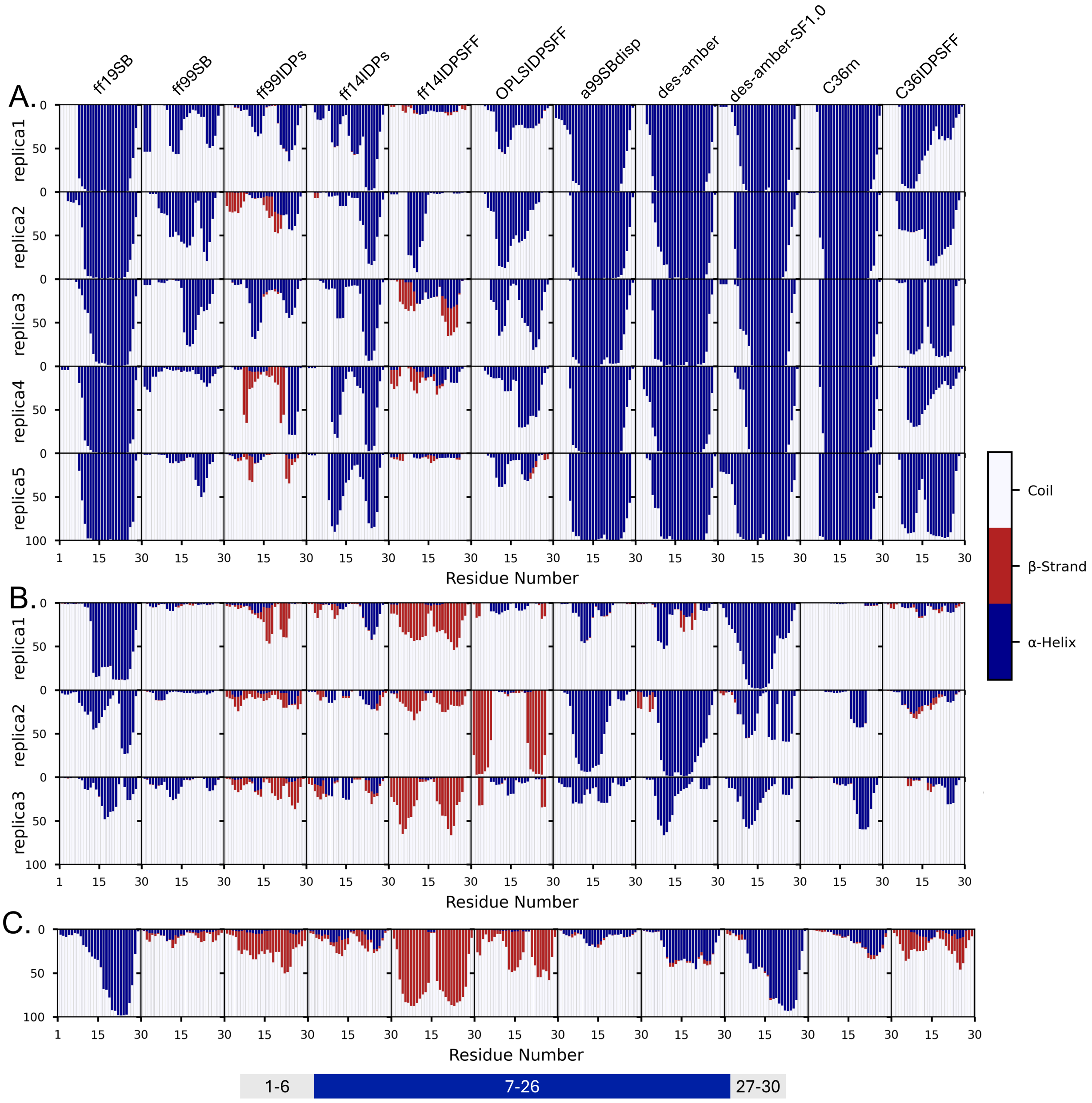


Figure S69: **Per-residue secondary structure preferences forGLP-1 (PDB ID 1D0R)** across simulations. Bars represent the percentage occurrence of each secondary structure type per residue in (A) five 200-ns replicas, (B) three 1-μs replicas, and (C) a 10-μs simulation. The secondary structure assignment in the experimentally folded structure is shown for reference. Red denotes $\beta$-strand, and blue denotes $\alpha$-helix or 3-10 helix, and the absence of color corresponds to coil or turn regions.

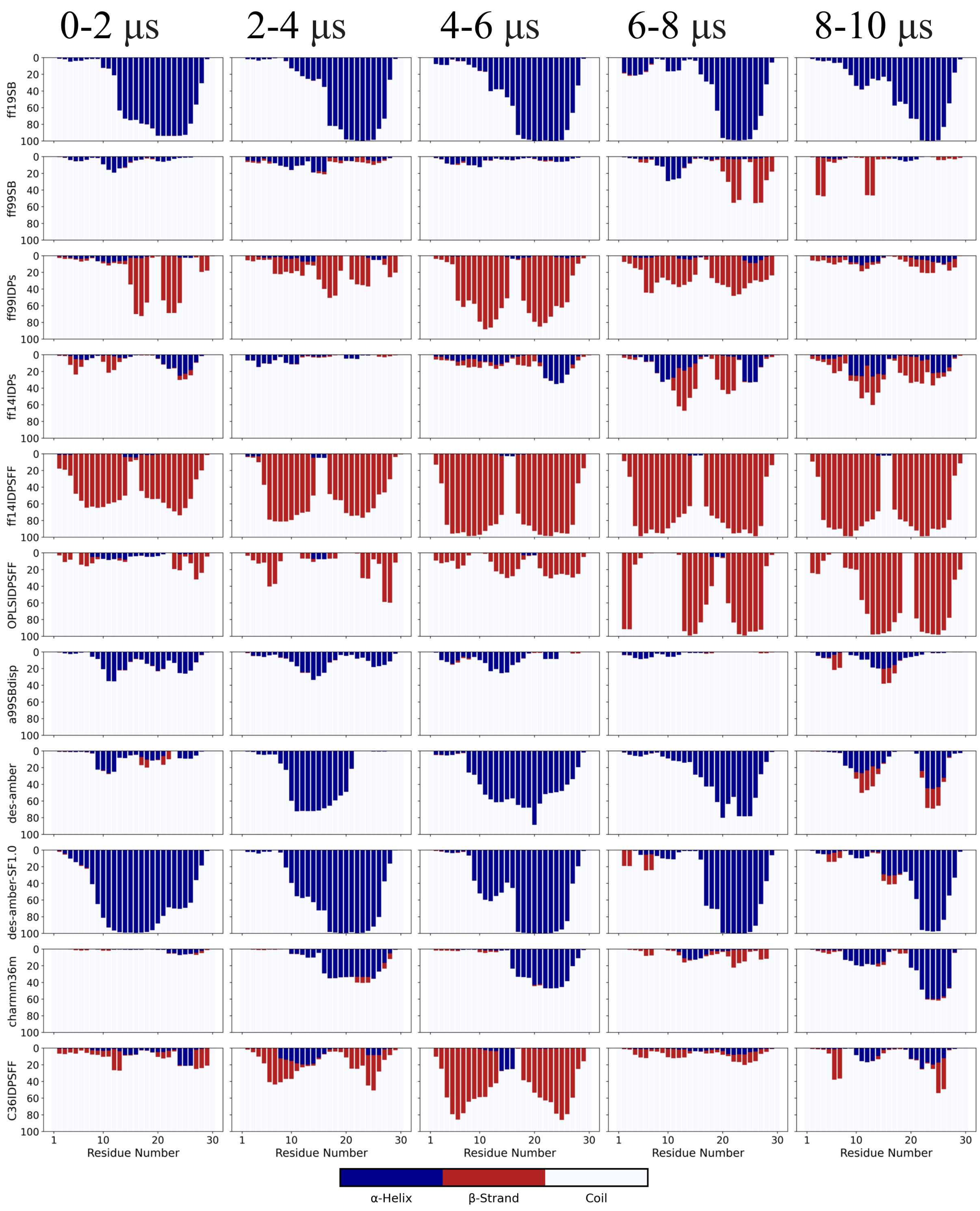


Figure S70: **Per residue secondary structure preferences for GLP-1 (PDB ID 1D0R) during the 10 $\mu$s simulation, shown as blocks of 2 $\mu$s intervals.** Bars represent the percentage occurrence of each secondary structure type per residue. Red indicates $\beta$-strand, blue indicates $\alpha$-helix or 3-10 helix, and the absence of color corresponds to coil or turn regions.

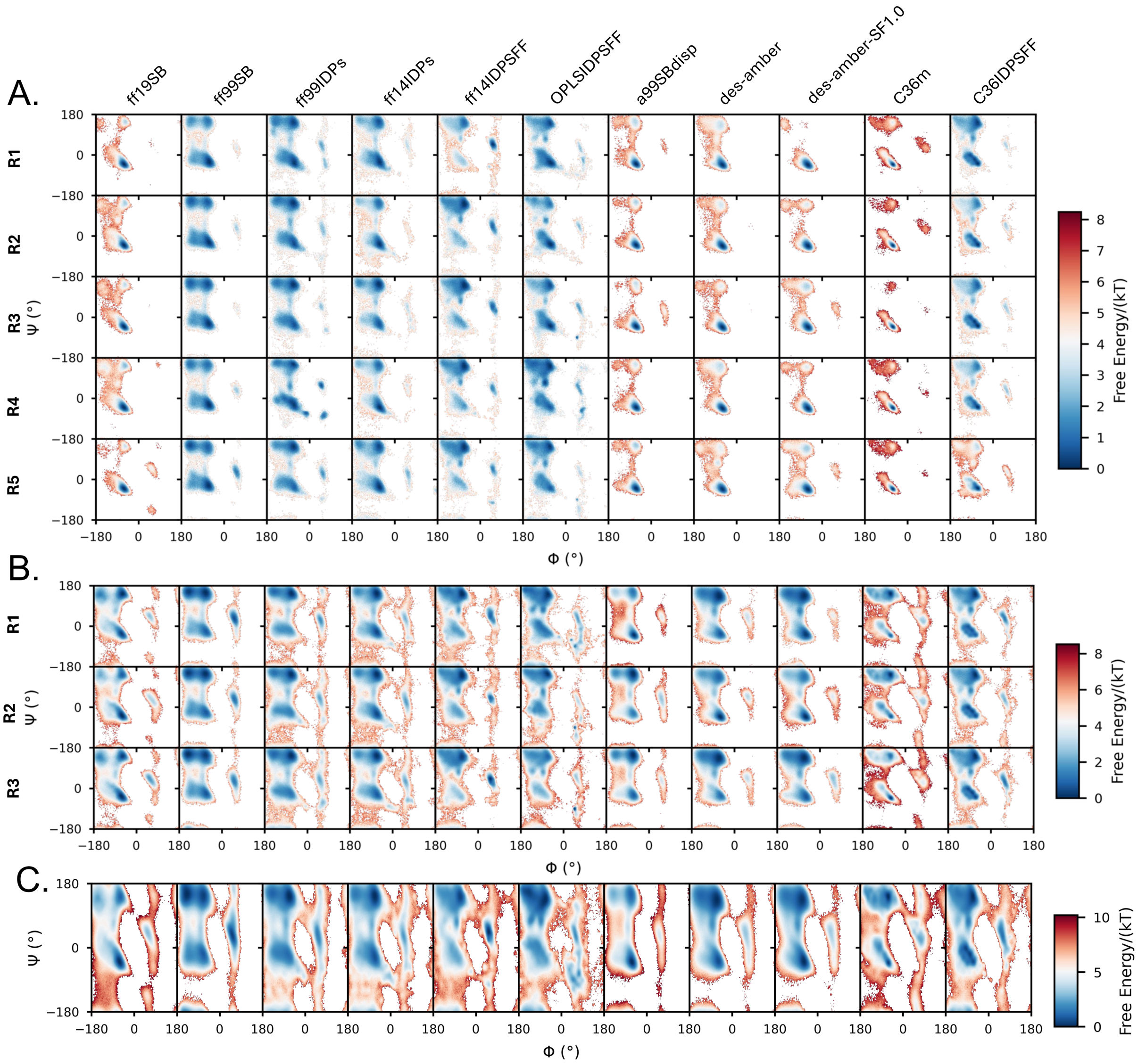


Figure S71: **Ramachandran plots for proadrenomedullin N-20 terminal (PDB ID 2FLY)**. (A) Five 200-ns simulation replicas initiated from the experimentally folded structure. (B) Three 1-μs simulation replicas initiated from the linear peptide sequence. (C) 10-μs simulation initiated from the linear peptide sequence.

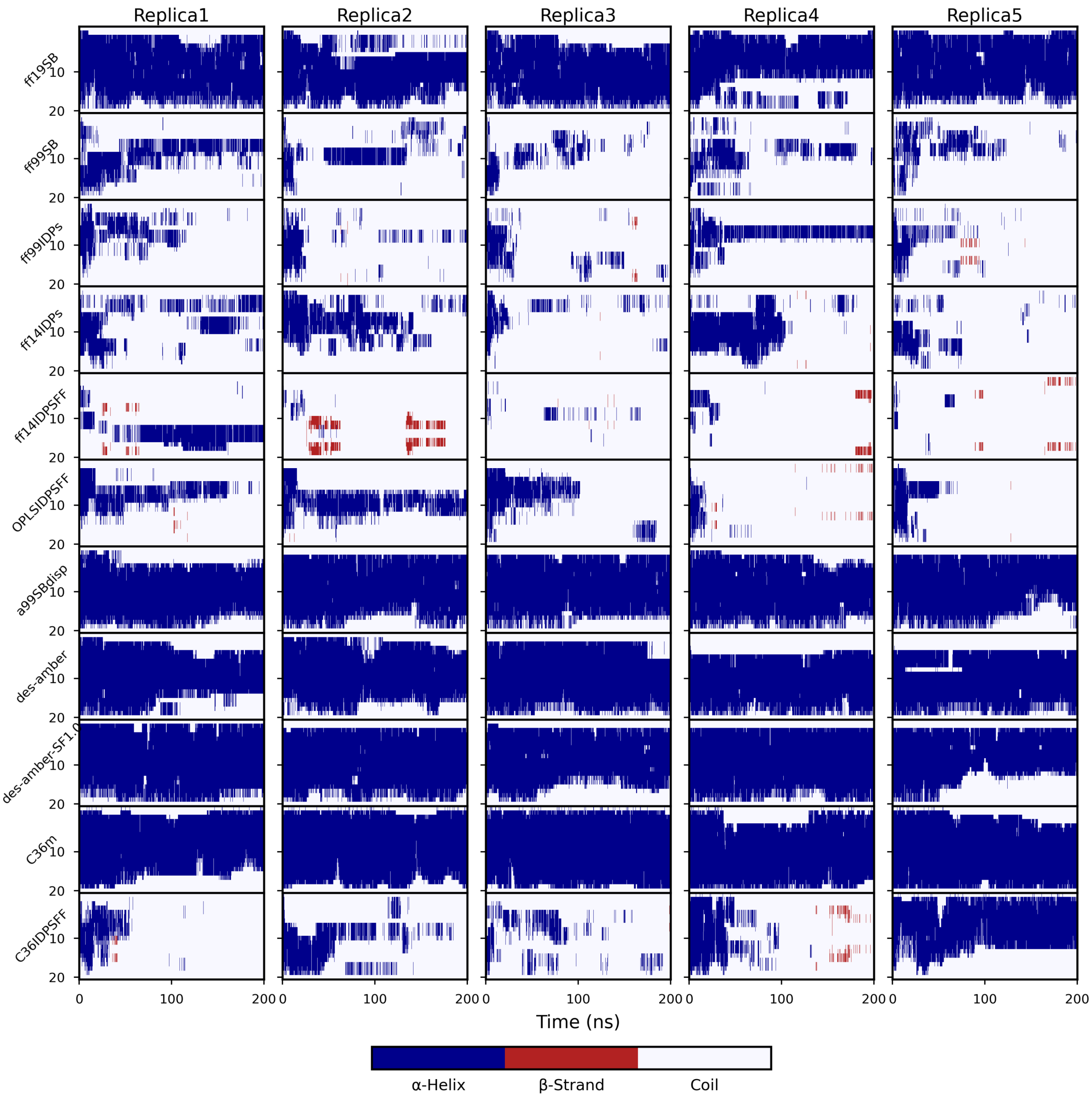


Figure S72: **Per-residue secondary structure time series for proadrenomedullin N-20 terminal (PDB ID 2FLY)** for five 200-ns simulation replicas initiated from the folded structure. Red denotes $\beta$-strand, and blue denotes $\alpha$-helix or 3-10 helix, and the absence of color corresponds to coil or turn regions.

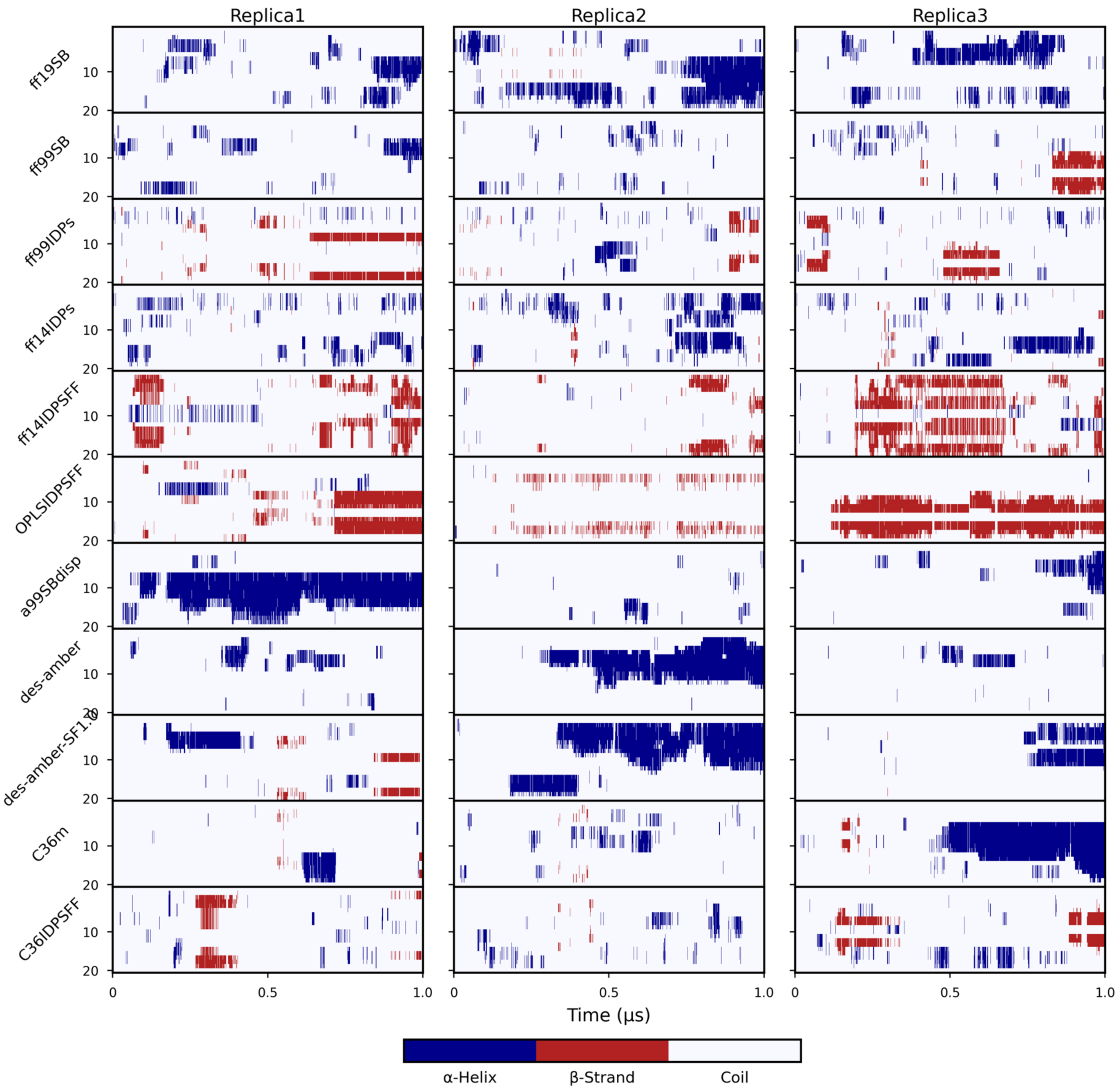


Figure S73: **Per-residue secondary structure time series for proadrenomedullin N-20 terminal (PDB ID 2FLY)** for three 1 $\mu$s simulation replicas initiated from the folded structure. Red denotes $\beta$-strand, and blue denotes $\alpha$-helix or 3-10 helix, and the absence of color corresponds to coil or turn regions.

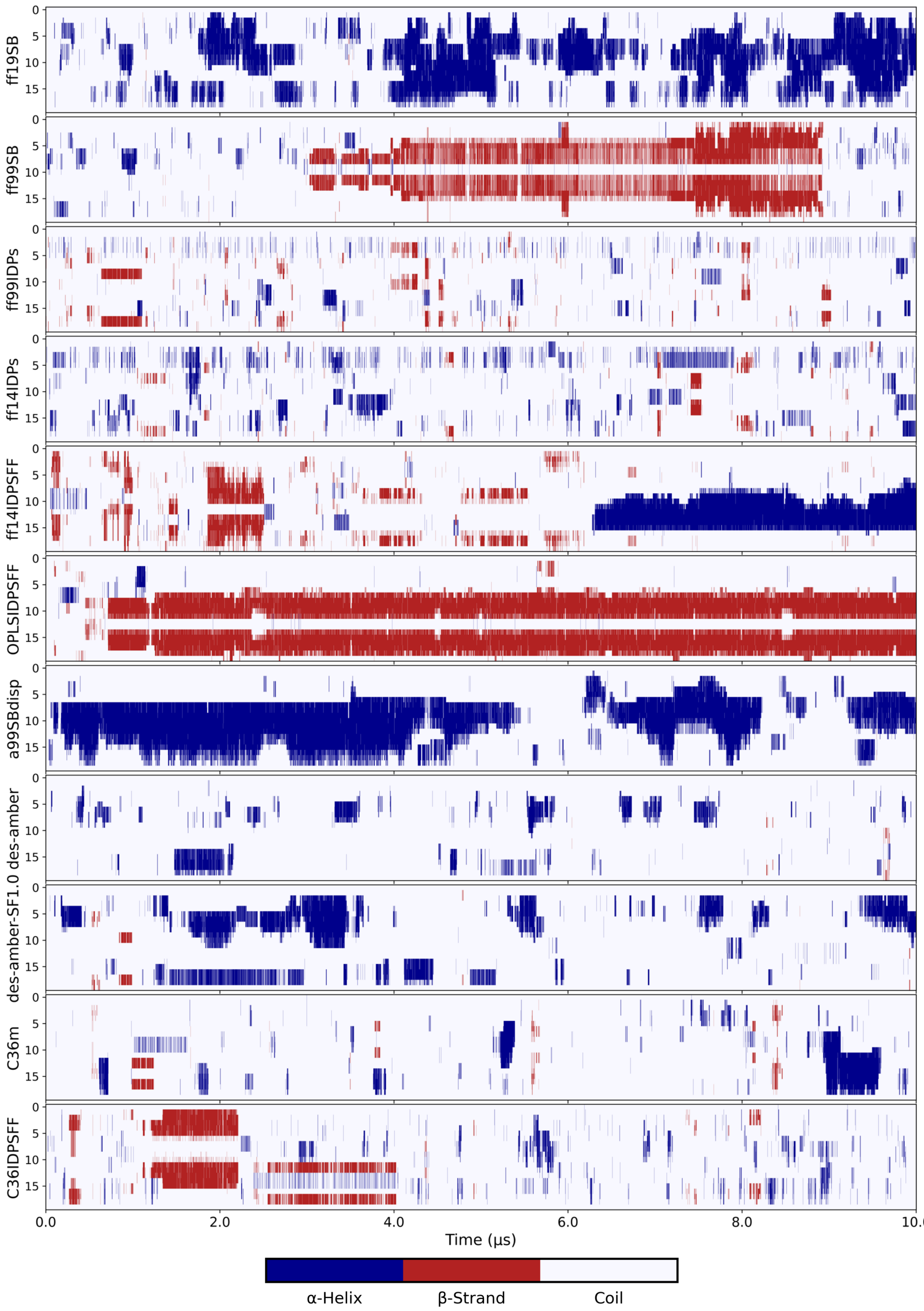


Figure S74: **Per-residue secondary structure time series for proadrenomedullin N-20 terminal (PDB ID 2FLY)** for the 10 $\mu$s simulation replicas initiated from the folded structure. Red denotes $\beta$-strand, and blue denotes $\alpha$-helix or 3-10 helix, and the absence of color corresponds to coil or turn regions.

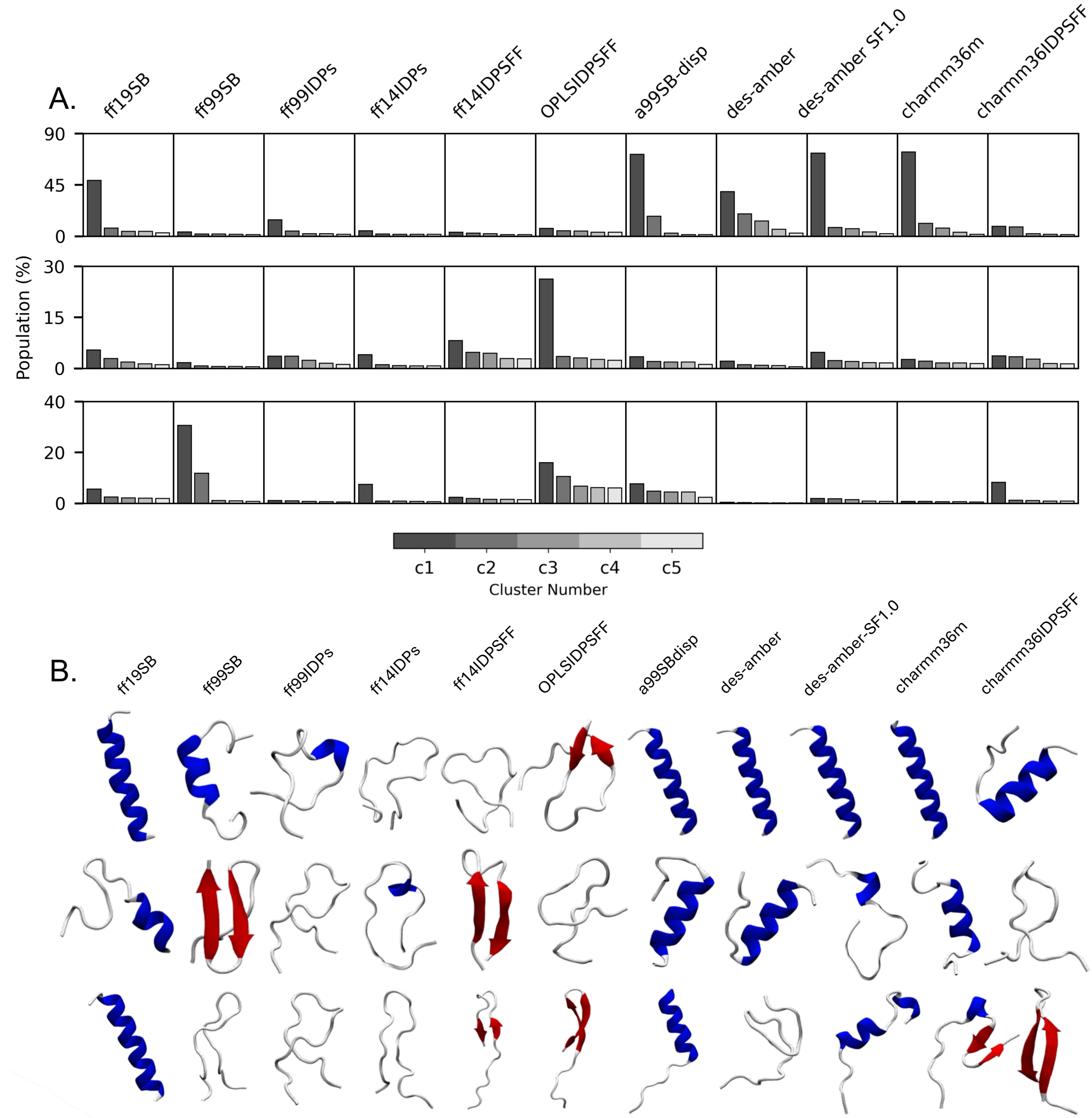


Figure S75: **Clustering results showing (A) cluster populations and (B) representative structures (centroids) of the most populated cluster for proadrenomedullin N-20 terminal (PDB ID 2FLY).** The top panel shows clustering from five 200-ns stability simulations initiated from the folded structure; the middle panel shows clustering from three 1-µs simulation replicas initiated from the linear peptide sequence; and the bottom panel shows clustering from a 10-µs folding simulation initiated from the extended peptide conformation. Red indicates $\beta$-strand, blue indicates $\alpha$-helix or 3-10 helix, and white indicates coil or turn regions.

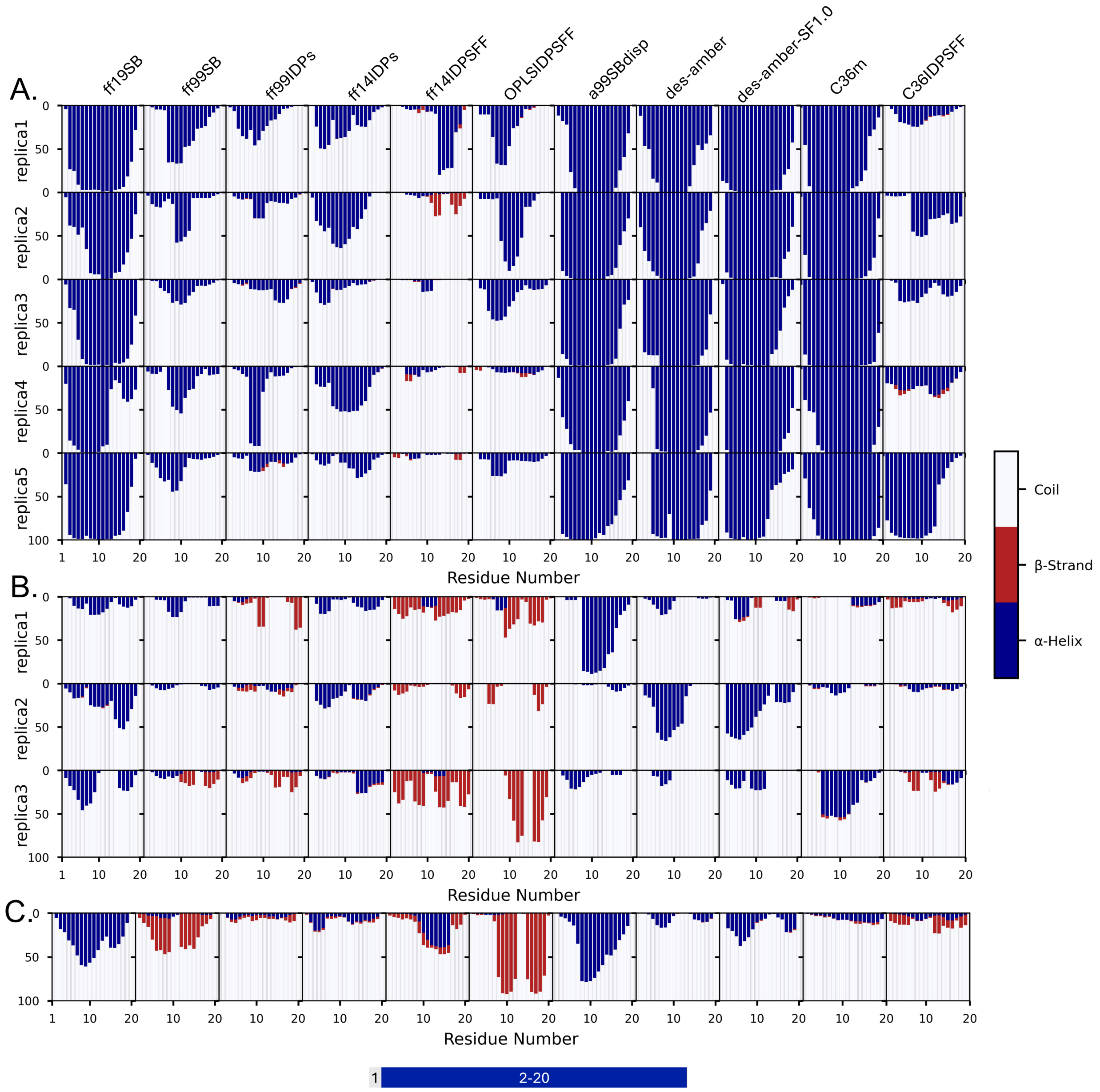


Figure S76: **Per-residue secondary structure preferences for proadrenomedullin N-20 terminal (PDB ID 2FLY)** across simulations. Bars represent the percentage occurrence of each secondary structure type per residue in (A) five 200-ns replicas, (B) three 1-µs replicas, and (C) a 10-µs simulation. The secondary structure assignment in the experimentally folded structure is shown for reference. Red denotes $\beta$-strand, and blue denotes $\alpha$-helix or 3-10 helix, and the absence of color corresponds to coil or turn regions.

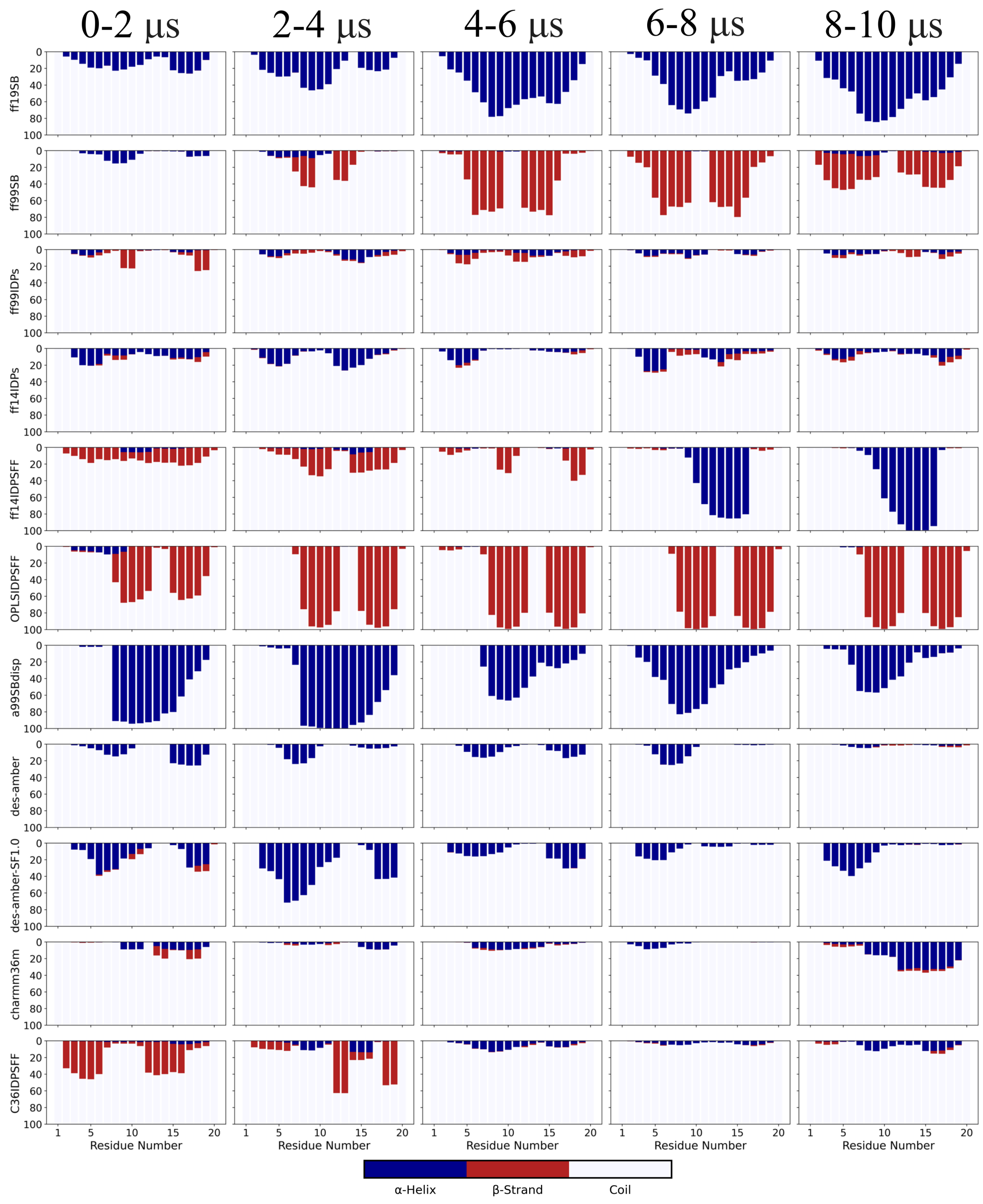


Figure S77: **Per residue secondary structure preferences for proadrenomedullin N-20 terminal (PDB ID 2FLY) during the 10 $\mu$s simulation, shown as blocks of 2 $\mu$s intervals.** Bars represent the percentage occurrence of each secondary structure type per residue. Red indicates $\beta$-strand, blue indicates $\alpha$-helix or 3-10 helix, and the absence of color corresponds to coil or turn regions.

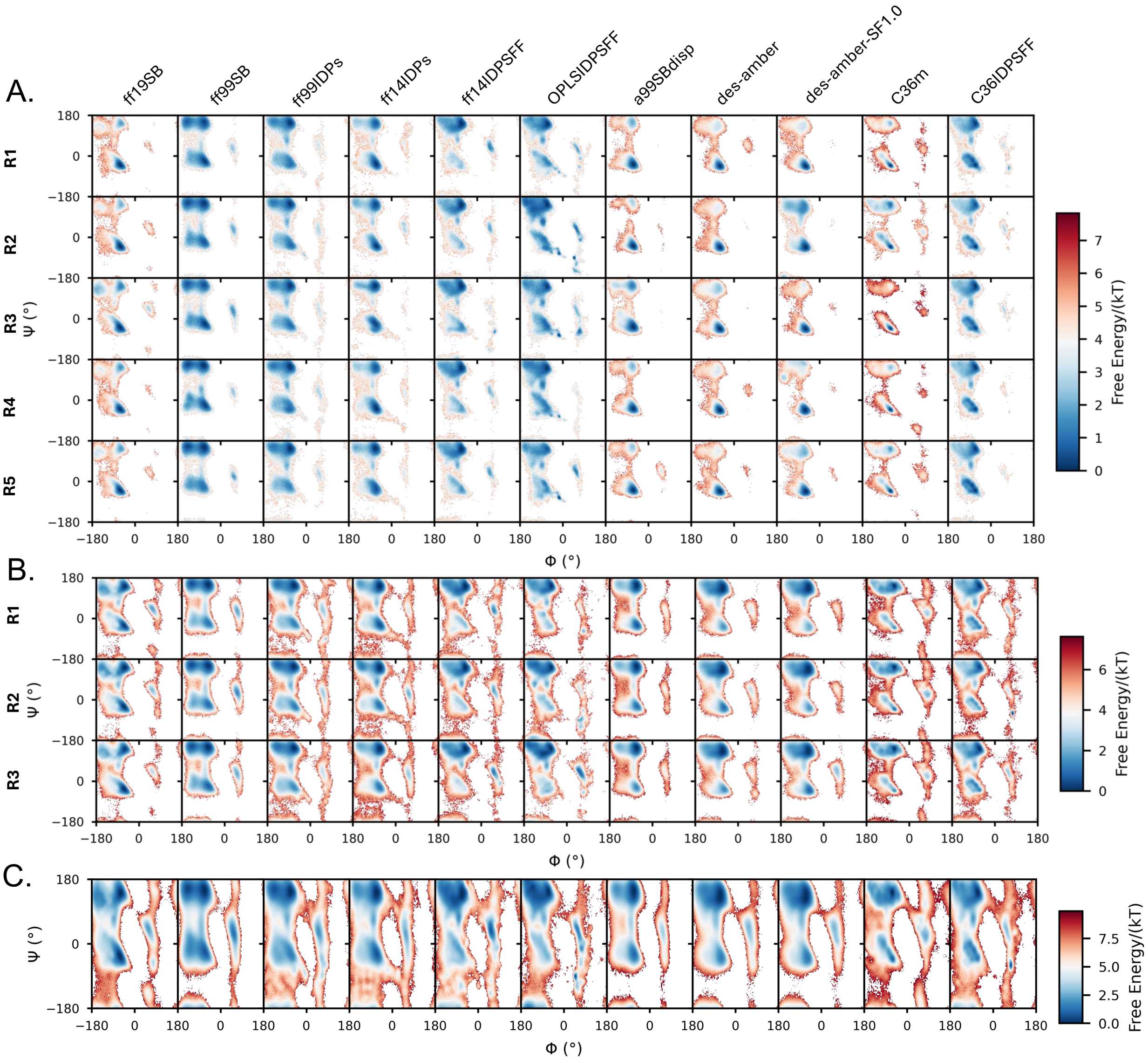


Figure S78: **Ramachandran plots for phylloseptin-1 (PDB ID 2JQ0)**. (A) Five 200-ns simulation replicas initiated from the experimentally folded structure. (B) Three 1-µs simulation replicas initiated from the linear peptide sequence. (C) 10-µs simulation initiated from the linear peptide sequence.

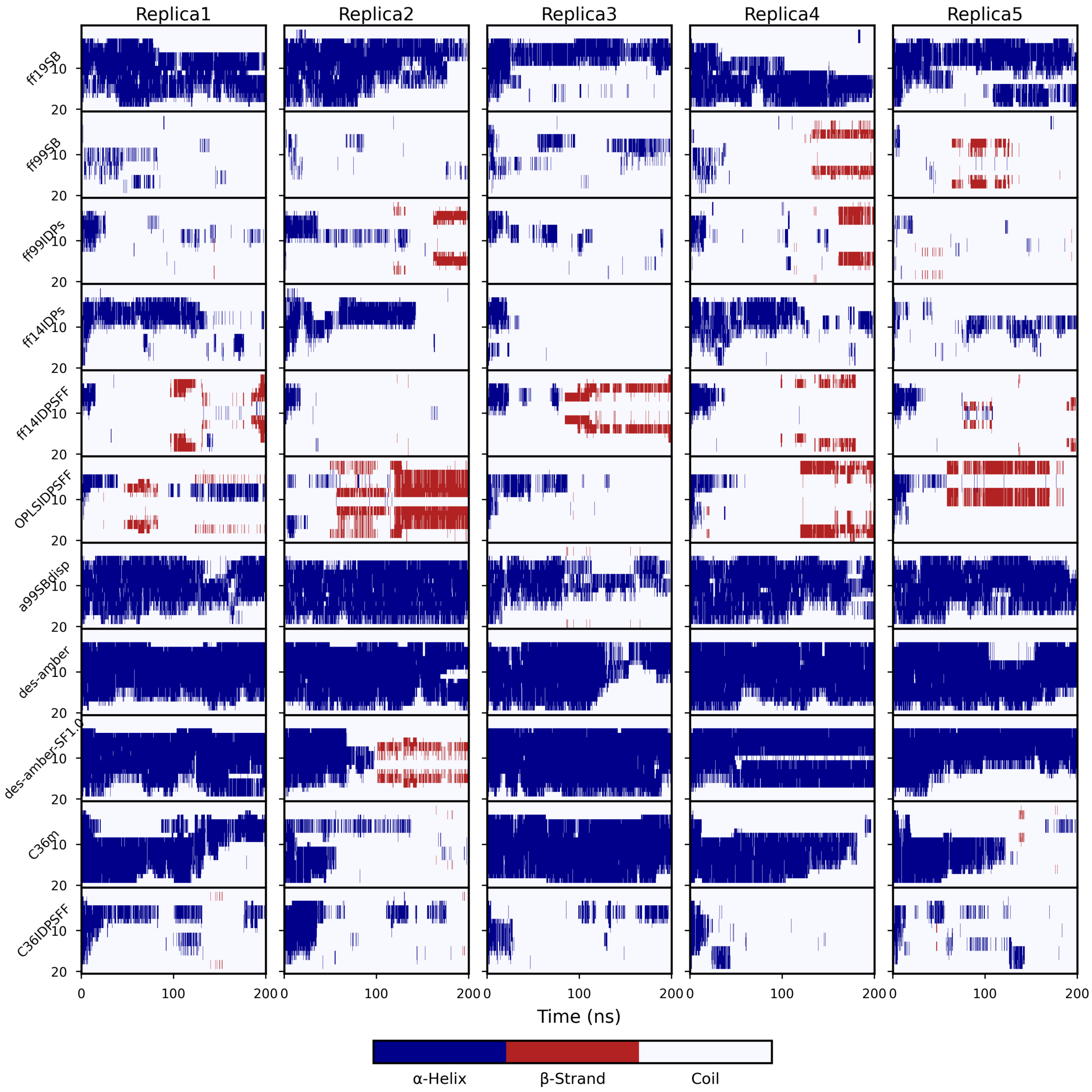


Figure S79: **Per-residue secondary structure time series for phylloseptin-1 (PDB ID 2JQ0)** for five 200-ns simulation replicas initiated from the folded structure. Red denotes $\beta$-strand, and blue denotes $\alpha$-helix or 3-10 helix, and the absence of color corresponds to coil or turn regions.

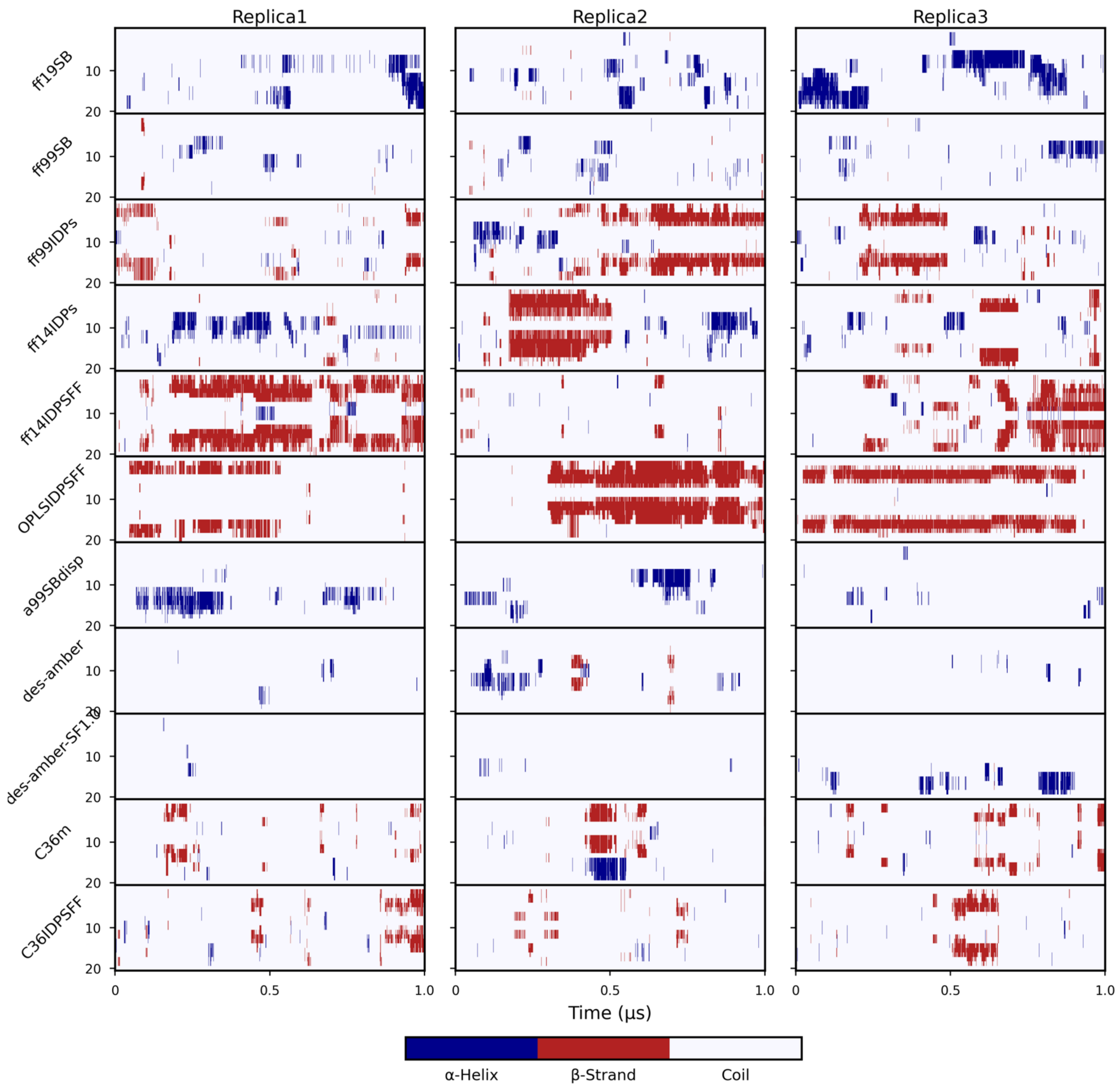


Figure S80: **Per-residue secondary structure time series for phylloseptin-1 (PDB ID 2JQ0)** for three 1 $\mu$s simulation replicas initiated from the folded structure. Red denotes $\beta$-strand, and blue denotes $\alpha$-helix or 3-10 helix, and the absence of color corresponds to coil or turn regions.

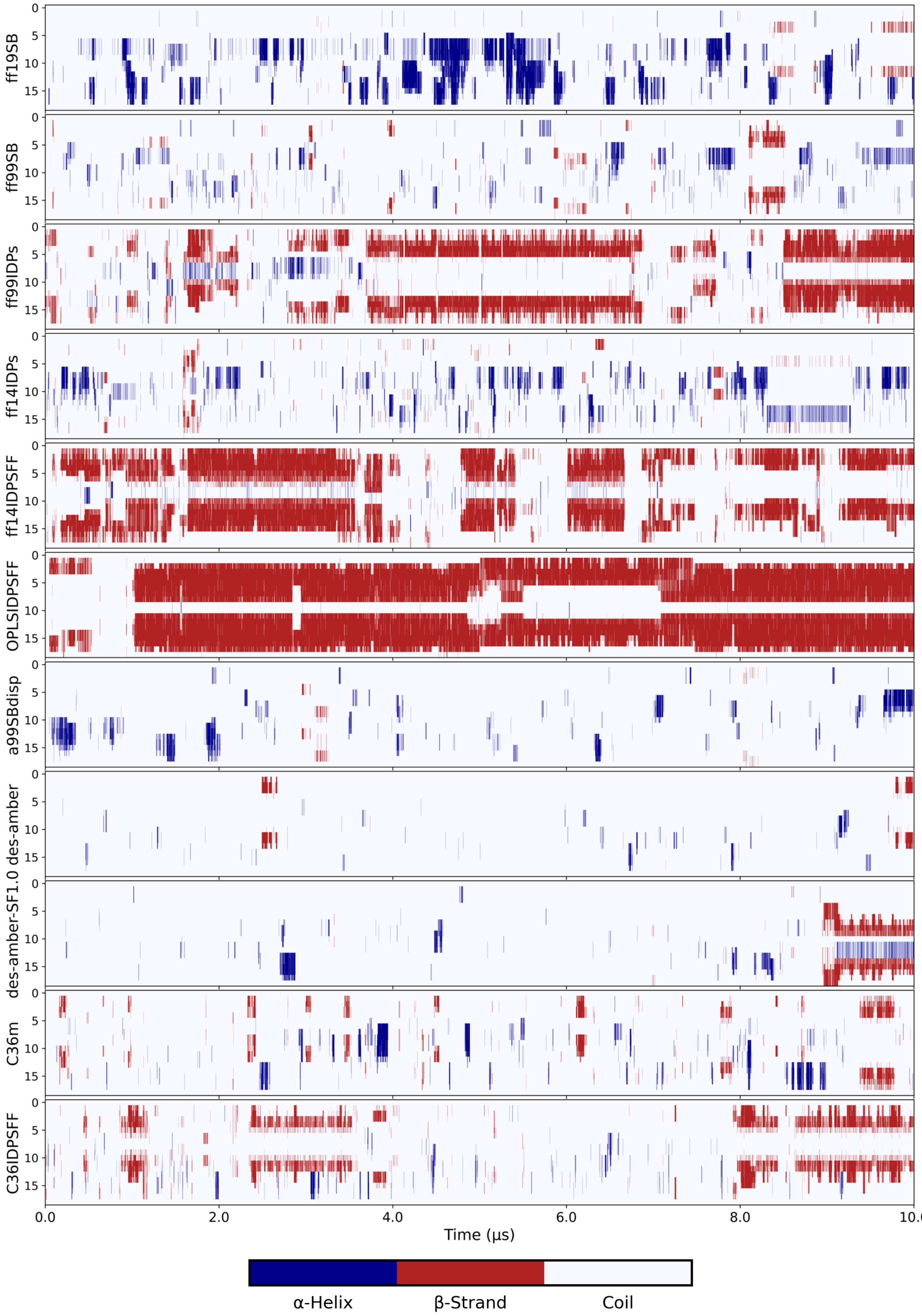


Figure S81: **Per-residue secondary structure time series for phylloseptin-1 (PDB ID 2JQ0)** for the 10 $\mu$s simulation replicas initiated from the folded structure. Red denotes $\beta$-strand, and blue denotes $\alpha$-helix or 3-10 helix, and the absence of color corresponds to coil or turn regions.

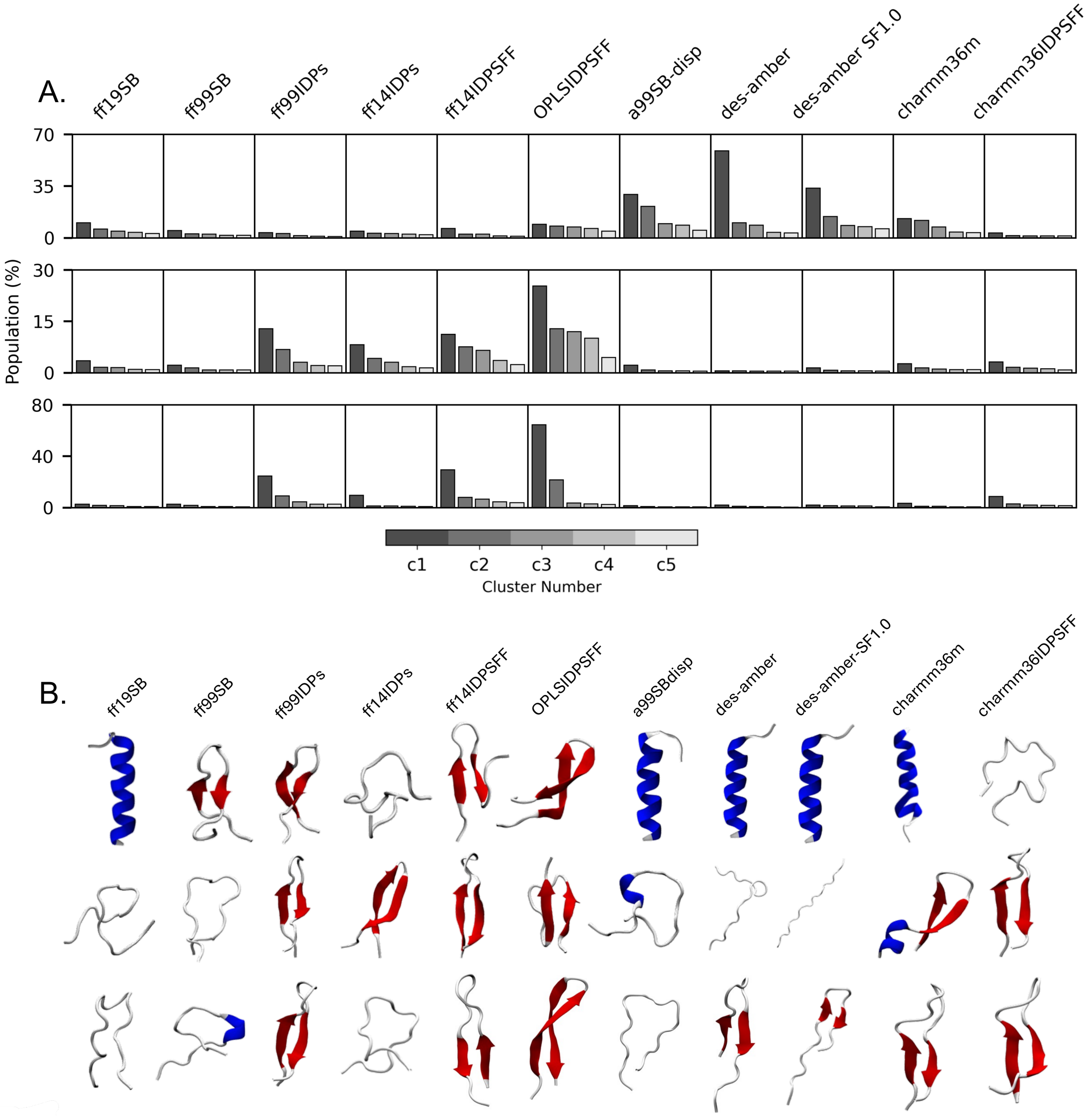


Figure S82: **Clustering results showing (A) cluster populations and (B) representative structures (centroids) of the most populated cluster for phylloseptin-1 (PDB ID 2JQ0).** The top panel shows clustering from five 200-ns stability simulations initiated from the folded structure; the middle panel shows clustering from three 1-µs simulation replicas initiated from the linear peptide sequence; and the bottom panel shows clustering from a 10-µs folding simulation initiated from the extended peptide conformation. Red indicates $\beta$-strand, blue indicates $\alpha$-helix or 3-10 helix, and white indicates coil or turn regions.

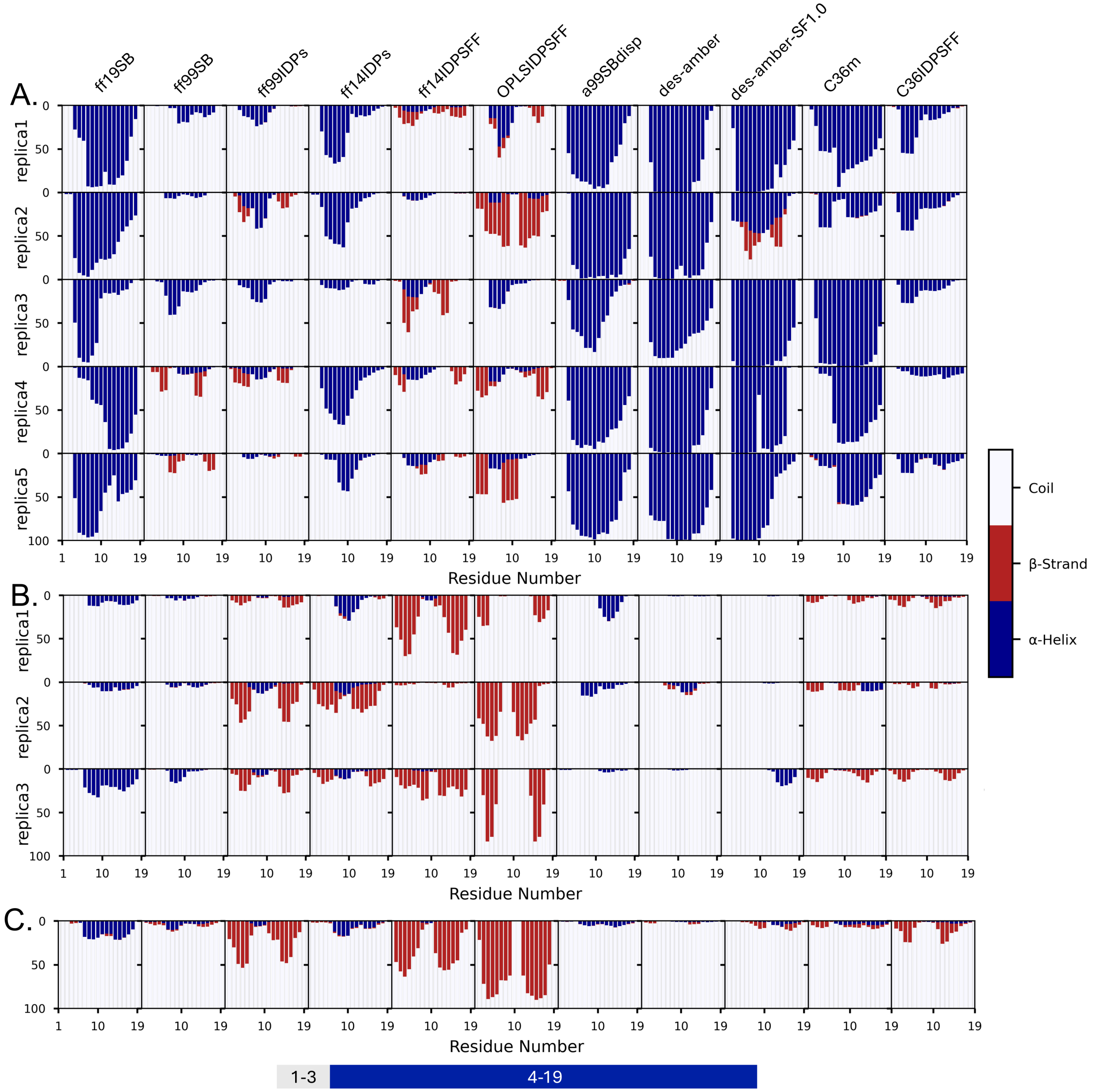


Figure S83: **Per-residue secondary structure preferences for phylloseptin-1 (PDB ID 2JQ0)** across simulations. Bars represent the percentage occurrence of each secondary structure type per residue in (A) five 200-ns replicas, (B) three 1-µs replicas, and (C) a 10-µs simulation. The secondary structure assignment in the experimentally folded structure is shown for reference. Red denotes $\beta$-strand, and blue denotes $\alpha$-helix or 3-10 helix, and the absence of color corresponds to coil or turn regions.

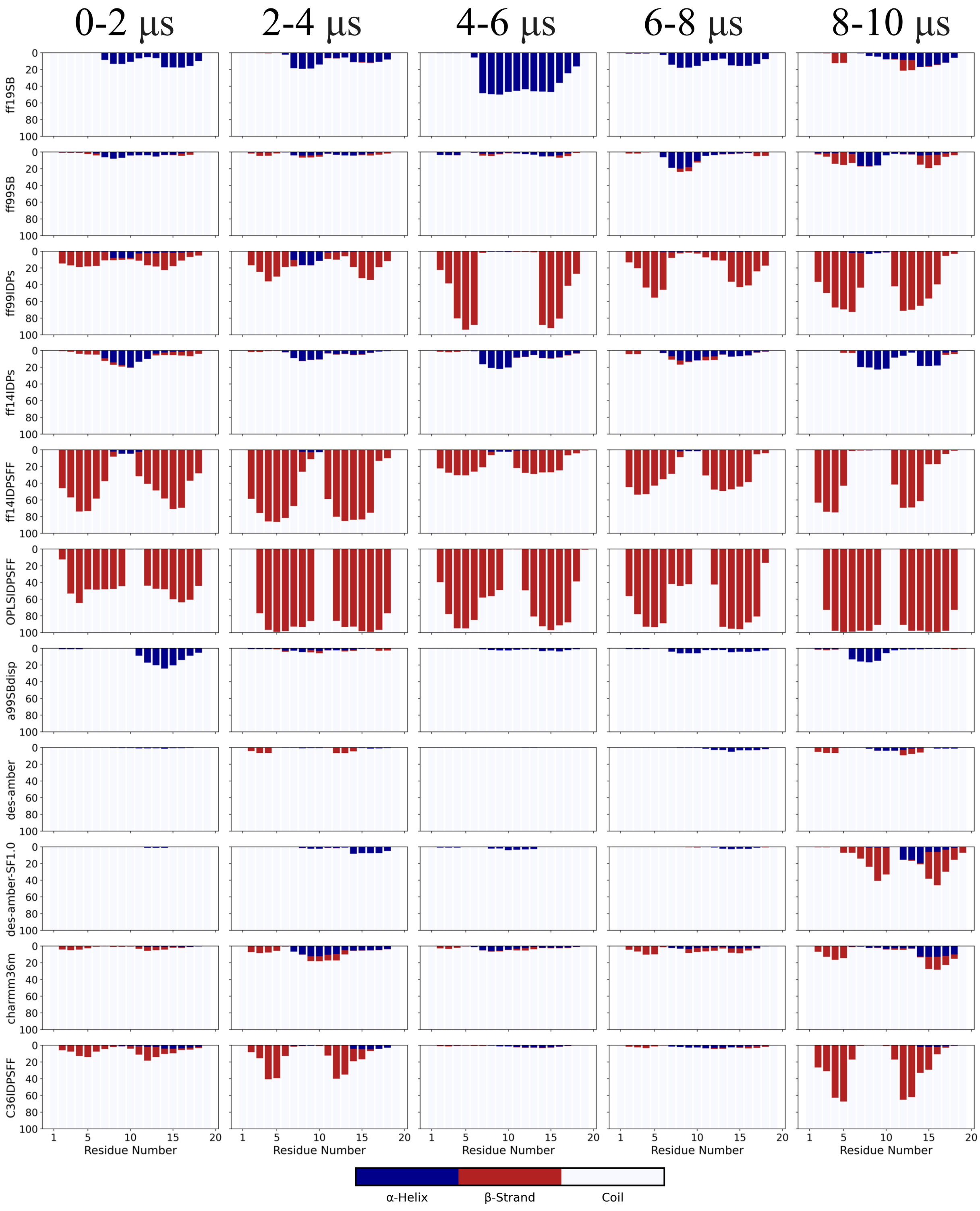


Figure S84: **Per residue secondary structure preferences for phylloseptin-1 (PDB ID 2JQ0) during the 10 $\mu$s simulation, shown as blocks of 2 $\mu$s intervals.** Bars represent the percentage occurrence of each secondary structure type per residue. Red indicates $\beta$-strand, blue indicates $\alpha$-helix or 3-10 helix, and the absence of color corresponds to coil or turn regions.

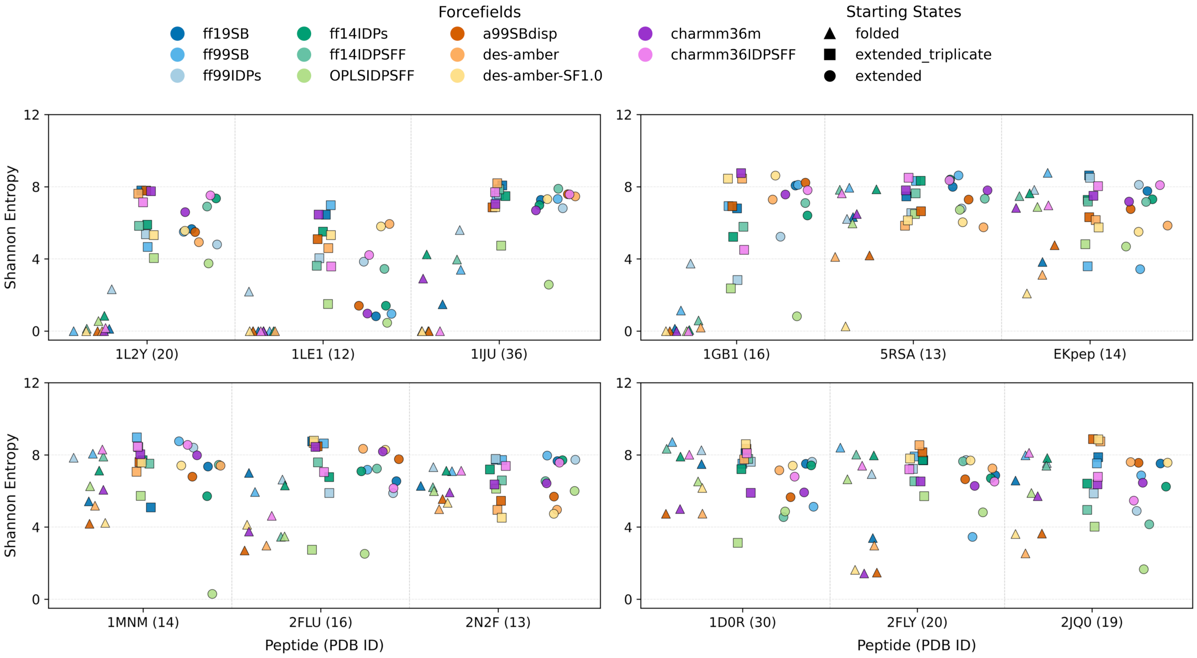


Figure S85: Comparison of system-wise Shannon entropy from five 200 ns simulations initiated from experimentally folded structures (triangle markers), three 1$\mu$s replica simulations initiated from linear peptide sequences (square markers), and 10$\mu$s simulations initiated from linear peptide sequences (circle markers). Numbers in parentheses indicate the sequence length of each peptide.

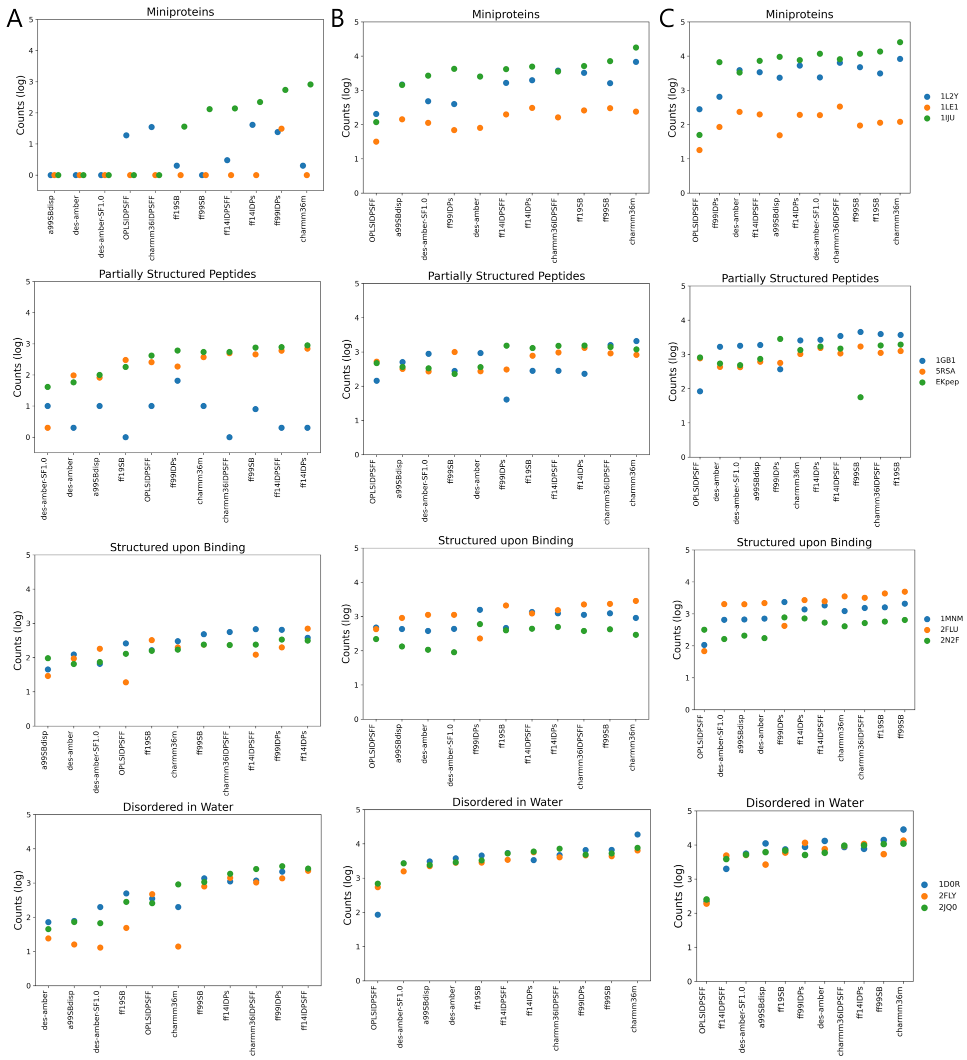


Figure S86: **Log-transformed cluster sizes for each force field and system.** (A) 200 ns simulations (five replicates) initiated from experimentally folded structures. (B) 1 $\mu$s simulations (triplicates) initiated from linear peptide sequences. (C) 10 $\mu$s simulations initiated from linear peptide sequences. Systems within each category are ordered by ascending cluster size.
